# From Stability to Complexity: A Systematic Review of Long-term Divergence Exponents in Nonlinear Gait Analysis

**DOI:** 10.64898/2025.12.18.695288

**Authors:** Jeremy Torrent, Roxane Coquoz, Philippe Terrier

**Author notes:** Corresponding Author: Philippe Terrier, Haute-Ecole Arc Santé, Espace de l’Europe 11, CH-2000 Neuchâtel, Switzerland. These authors contributed equally to this work (Jeremy Torrent, Roxane Coquoz).

## Abstract

Divergence exponents (DEs), also termed maximum Lyapunov exponents, computed from stride-to-stride fluctuations have traditionally been interpreted as measures of gait stability. We evaluated empirical evidence for reinterpreting long-term DE as a measure of gait complexity and automaticity. We systematically searched Web of Science databases through January 2026 for studies applying Rosenstein’s algorithm to human gait, extracting experimental conditions, and participant characteristics from each study. Study quality was assessed across analytical rigor, outcome reporting, and sample size adequacy. We conducted a meta-analysis examining correlations between long-term DE and detrended fluctuation analysis (DFA) scaling exponents, and synthesized evidence from perturbation, cueing, and between-subject studies. Sixty-two studies published between 2000 and 2026 met inclusion criteria, with 44% achieving high overall quality scores. Meta-analysis from six datasets (209 participants) revealed a positive correlation between long-term DE and DFA scaling exponents (r=0.64, 95% CI 0.34 to 0.82; I²=82%). Perturbation studies consistently increased short-term DE while decreasing long-term DE by up to 51%. Auditory and visual cueing reduced long-term DE (up to −86%) while minimally affecting short-term DE. Between-subject comparisons revealed heterogeneous patterns, with clinical populations exhibiting pathology-dependent long-term DE changes. Converging meta-analytic and experimental evidence supports reinterpreting long-term DE as the Attractor Complexity Index—a measure of gait complexity and automaticity rather than stability. Reduced long-term DE during controlled gait conditions reflects suppression of low-frequency variability, which constrains phase space exploration and accelerates divergence curve saturation. The measure’s sensitivity to prefrontal cortex engagement and attentional demands indicates that long-term DE reflects gait automaticity: lower values mark a shift from subcortical to executive control. This attention-dependent reorganization explains patterns observed in aging, clinical populations, and experimental perturbations, and establishes long-term DE as a complementary biomarker for motor-cognitive aspects of gait control, with implications for fall risk assessment, disease monitoring, and rehabilitation evaluation.

## Introduction

Walking is a complex motor activity that plays an important role in daily functioning [1]. Gait—the pattern of limb movement—can serve as a significant biomarker, providing insights into an individual’s physical functioning [2,3], cognitive abilities [4], and level of independence [5]. Slower speeds and deviations from normal gait patterns have been shown to correlate with adverse outcomes, including an increased risk of falls [6,7], cognitive decline [7,8], and mortality [9]. For this reason, gait analysis is an essential component of clinical evaluation across medical specialties [10,11].

Nonlinear gait analysis is a methodological approach that examines the seemingly unpredictable dynamics of human gait using mathematical and statistical tools from nonlinear systems theory [12]. Nonlinear dynamical systems are characterized by outputs that are not proportional to changes in inputs, where the behavior cannot be described by simple linear differential equations. In the context of physiological signals, nonlinearity reflects how small changes in one component can lead to disproportionate or emergent effects throughout the system. These complex interactions arise from the integration of multiple regulatory mechanisms—such as the interplay between the autonomic nervous system and neuroendocrine factors in heart rate regulation—forming a dynamic network whose behavior cannot be fully understood by analyzing individual components in isolation [13,14]. Nonlinear analysis techniques have become an integral part of gait analysis, providing advanced methods such as maximum Lyapunov (or divergence) exponents for assessing resilience to perturbations [15], fractal dynamics for investigating the complexity and statistical persistence of time series [16], and entropy measures for quantifying signal irregularity [17]. These methods facilitate the detection of gait disorders [18–20], and support the development of innovative rehabilitation strategies [21]. By going beyond traditional kinematic and kinetic parameters, nonlinear approaches allow for a more comprehensive assessment of gait variability [22].

Nonlinear gait analysis also provides a window into the underlying neural control mechanisms [23], including the involvement of executive function in gait [4]. Attention, as a component of executive function, refers to the ability to allocate cognitive resources among tasks, including gait, that are performed concurrently [4,24]. Gait automaticity reflects the extent to which executive function contributes to gait control, with highly automatic gait requiring little attentional resources and less automatic gait requiring increased executive involvement [25]. One particularly powerful approach for investigating the interplay between executive function and gait is the use of dual-task paradigms. In these experimental setups, participants are asked to perform a cognitive task while walking, which allows researchers to assess how the allocation of attentional resources affects gait parameters [4]. Nonlinear measures applied to dual-task walking data can reveal how the complexity and stability of gait are modulated by concurrent cognitive demands, providing valuable information about the adaptive capacity of the motor control system and the degree of gait automaticity [26,27]. This approach has proven useful in understanding age-related changes in gait control [28] and in identifying early markers of cognitive decline [29] and fall risk [30] in various clinical populations.

Beyond dual-task paradigms, various factors (including sensory inputs, environmental challenges, and individual physiological and psychological states) can influence the extent of executive function involvement during walking, and nonlinear gait analysis provides valuable tools for exploring these dynamics. For example, the influence of auditory cues on gait automaticity can be observed through rhythmic auditory stimulation, where external rhythms entrain motor rhythms, potentially bypassing impaired internal gait control mechanisms in conditions such as Parkinson’s disease [31,32]. Exposing older adults to visual perturbations significantly reduces the complexity of their gait, a response not observed in younger adults. This may be indicative of an increased reliance on visual feedback and a decrease in gait automaticity with age [33]. Environmental constraints, such as treadmill walking, can alter motor control strategies. Research has shown that stride-speed fluctuations exhibit statistical anti-persistence during treadmill walking, indicating tight control over this gait parameter. This tight control is further enhanced when maneuverability is restricted, suggesting that the nervous system actively minimizes speed deviations [34]. These findings are consistent with the idea that the need for continuous speed adjustments on a treadmill may increase the demands on executive function to maintain a steady gait [35]. Individual physiological and psychological states, such as pain, attentional focus, or anxiety, may also influence gait automaticity. For example, research suggests that individuals with chronic low back pain exhibit altered gait complexity, possibly due to attentional strategies focused on managing pain [36]. In addition, fear of falling is associated with reduced local dynamic stability of gait [37], which may be explained by an increased reliance on executive functions to maintain balance and adapt to perceived threats during walking.

Among nonlinear measures, the maximum Lyapunov exponent [divergence exponent (DE)] method stands out as a prominent measure for assessing dynamic stability and fall risk [20,38,39]. This prominence is due to its ability to quantify local dynamic stability—a system’s sensitivity to small perturbations [15,40,41]. Modeling studies using dynamic walker systems have validated its predictive validity, demonstrating strong correlations between short-term DE and simulated fall probabilities [42,43]. These computational validations, combined with clinical observations of elevated DE in fall-prone elderly, position it as an important tool for assessing gait stability and fall risk in both research and clinical applications [20]. However, recent systematic reviews have shown heterogeneity in the implementation of the DE method, with different parameterizations and measurement systems across studies [20,44], highlighting the need for further research to standardize methodologies and facilitate cross-study comparisons.

A persistent methodological challenge in DE-based gait analysis arises from the differential responsiveness of short- and long-term DEs computed by Rosenstein’s algorithm. Mathematically, Rosenstein’s method embeds the time series in a state space and identifies for each point a nearest neighbor such that their initial separation *d*(0) evolves with time *t* as *d*(*t*) = *d*(0)*e*^λ*t*^, where λ is the DE characterizing the intensity of the exponential divergence, which is also known as the “butterfly effect” [45]; taking the natural logarithm gives *ln*(*d*(*t*)) = *ln*(*d*(0)) + λ *t*, and hence 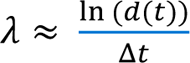 [15,40]. The characteristic logarithmic divergence curve is computed to assess the average progressive separation. This curve-based formulation is specific to Rosenstein’s algorithm: alternative approaches such as Wolf’s compute a single Lyapunov exponent through trajectory renormalization rather than a divergence curve [46]. Short-term DE is computed as the average rate of divergence by linear regression of the initial segment of the curve (0-1 step or stride), capturing local dynamic stability through immediate corrective mechanisms. Conversely, long-term DE is derived from later intervals (4-10 strides) and reflects long-term system behavior. While elevated short-term DE consistently correlates with fall risk and other variability metrics [47], long-term DE exhibits paradoxical responses to experimental manipulations, decreasing during cognitively demanding walking [33,48] and showing weak, or even opposite, associations with stability measures [47,49].

Therefore, while long-term DE was initially developed to measure gait stability, accumulating evidence suggests that it captures aspects of gait control distinct from those reflected by short-term DE. Several studies have demonstrated strong associations between long-term DE and the correlation structure of stride intervals, as quantified by scaling exponents that characterize stride-to-stride fluctuation complexity [40,50,51]. This relationship is particularly evident in experimental conditions that modify gait automaticity. Metronome walking, which requires conscious sensorimotor synchronization and increased attentional demands, substantially decreases both long-term DE and gait complexity while leaving short-term DE unchanged [48,52]. Furthermore, it has been shown that gait acceleration signals with identical shapes, but modified stride interval complexity exhibit different long-term DE values [53]. These findings have led to its reconceptualization as the “Attractor Complexity Index” (ACI), reflecting its role in quantifying gait complexity and automaticity rather than stability [53].

The primary objective of this systematic review is to examine the use of long-term DE in nonlinear gait analysis, with particular focus on its reinterpretation as a measure of gait complexity and automaticity rather than stability. A secondary objective is to characterize the substantial methodological heterogeneity observed across studies, including variations in measurement systems, signal processing techniques, and parameterization of Rosenstein’s algorithm, with the aim of identifying best practices for future research. By integrating these conceptual and methodological perspectives, this synthesis seeks to facilitate more reliable and comparable applications of long-term DE in both research and clinical contexts.

## Methods

### Reporting and transparency

The systematic review followed the PRISMA guidelines to ensure transparency and comprehensive reporting [54]. However, it is important to note that the protocol for this review was not registered in PROSPERO or any other repository. Methodological reviews focusing on tool development or reporting practices without direct clinical applicability are typically excluded from PROSPERO registration requirements [55]. Despite the lack of formal registration, efforts were made to maintain transparency by publishing the protocol as a non-peer-reviewed preprint [56]. The readers are invited to consult the published protocol for further details about the review procedures.

### Study eligibility criteria

The review included original peer-reviewed research articles focusing on human locomotion using the maximum Lyapunov exponent method with Rosenstein’s algorithm to compute long-term DE. Database searches applied no language restrictions to ensure comprehensive retrieval of potentially relevant literature. Eligible studies investigated gait in healthy participants or patients across all age groups and were published between 2000 and 2024. Observational, experimental, and intervention studies were included if they reported sufficient methodological details on DE computation. Studies were excluded if they: focused on modes of locomotion other than walking (e.g., running or jumping), relied exclusively on short-term DE or alternative analyses (e.g., Wolf’s algorithm), were pure modeling or simulation studies lacking empirical human data, or did not adequately describe the DE computation parameters. Reviews, letters, editorials, and conference abstracts were also excluded. Full-text screening was restricted to articles published in English or French based on the linguistic competencies of the review team. Articles identified in languages other than English or French underwent title, abstract, and methods section screening using automated translation tools to assess eligibility against inclusion criteria.

### Updated search (2024–2026)

To ensure the currency of the evidence base, an updated search was performed in January 2026 using the same databases and the identical search strategy, with the publication period restricted to 2024–2026. The updated screening applied the same two-phase selection process as the initial search. Two reviewers independently screened titles and abstracts, then assessed full texts for eligibility using the original inclusion criteria. Disagreements were resolved through discussion between the two reviewers.

### Information sources and search strategy

We performed a systematic search using the Web of Science platform, including MEDLINE and the Web of Science Core Collection databases. The search strategy combines themes related to gait analysis, nonlinear dynamics, and Lyapunov exponents. Boolean operators and truncations were used to ensure comprehensive coverage. The search keywords are described in the supporting information (S1 Text).

### Study selection process

The study selection process was conducted in two phases, the results of which are summarized in a PRISMA flowchart (Fig 1). Initially, two independent reviewers screened the titles and abstracts of all retrieved articles using broad inclusion criteria, as specific references to long-term divergence exponents were most often missing from abstracts. Articles potentially involving nonlinear gait analysis were advanced to the second phase, where their full texts were thoroughly reviewed to confirm the use of long-term divergence exponents calculated with Rosenstein’s algorithm. Disagreements between reviewers were resolved through discussion, with arbitration by a third reviewer when necessary [56].

**Fig 1.**
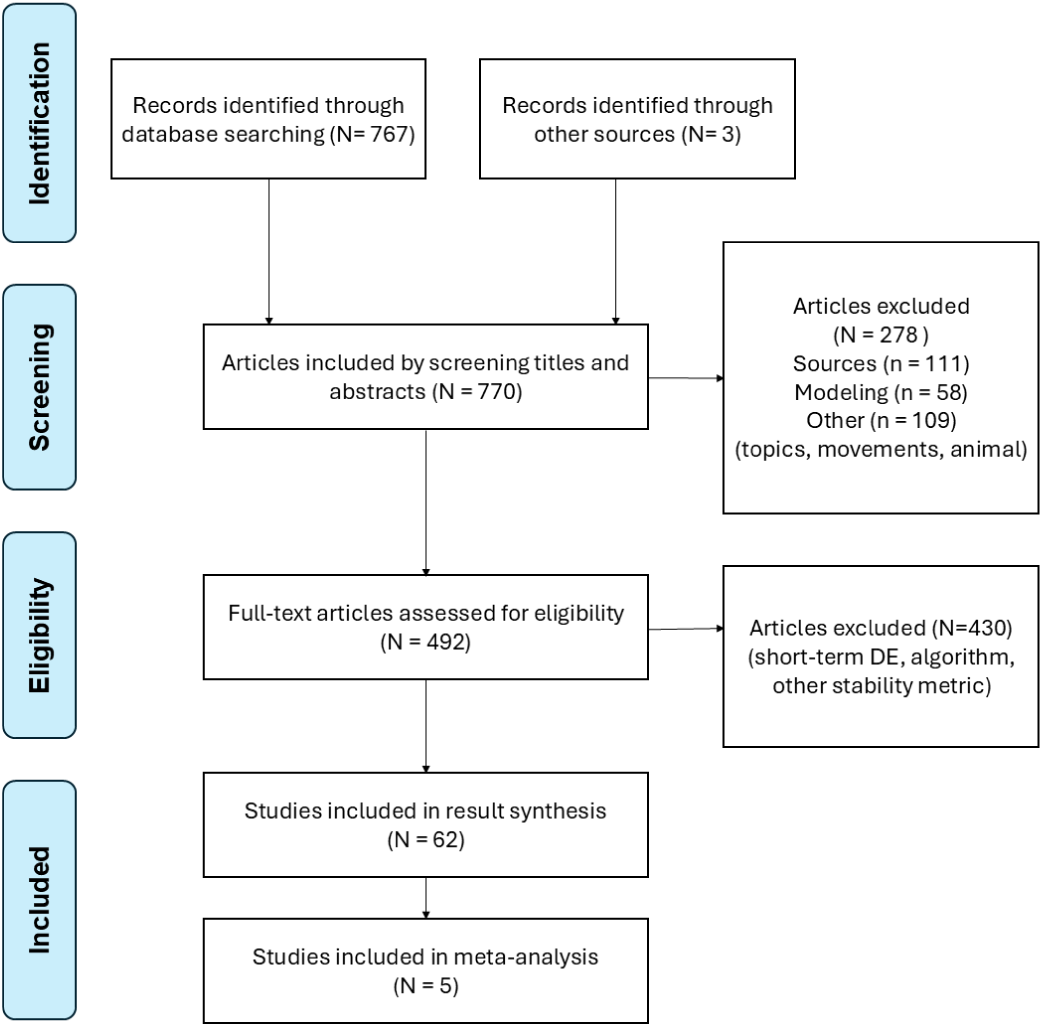
PRISMA flowchart of systematic review study selection. Flow diagram showing identification, screening, eligibility assessment, and final inclusion of studies examining long-term divergence exponents in human gait analysis.

### Data collection process and items

#### Main outcomes

Study outcomes were collected using standardized extraction tables as specified in the protocol [56], and the extracted results are available in the supporting information for detailed reference (S1 Text). For each included study, three main categories of information were extracted: study characteristics, methodological specifications, and outcomes related to short-term and long-term DEs. Study characteristics included publication year, authorship, sample size, participant demographics (age, health status), and research context (e.g., methodological studies or clinical evaluations). Methodological specifications included details on measurement methods (e.g., primary measurement system, signal type, sampling rate), walking conditions (overground or treadmill), signal processing procedures, and specific computational parameters for DEs (e.g., time delay, embedding dimension). Finally, the extracted findings focused on study objectives, conclusions, and interpretations of DE results.

To enable comparison of findings across studies, we computed relative percent change for each reported contrast, defined as the difference between the experimental and reference condition divided by the reference condition value, multiplied by 100 (Tables 1–3). Three considerations motivated this standardization. First, this standardization was necessary because absolute DE values depend on the measurement system, signal type, and anatomical location, rendering raw values incommensurable across the diverse methodological approaches represented in the corpus. Expressing results as percent change removes this baseline dependency while preserving the direction and relative magnitude of effects within each study. Second, percent change functions as a practical effect size metric, which is preferable to relying solely on statistical significance in this literature: many included studies used sample sizes that may be underpowered for detecting between-condition differences in long-term DE, and reporting effect magnitudes allows consistent identification of response patterns across studies regardless of whether individual comparisons reached significance. Third, because many numerical values were extracted from published figures using a digital extraction tool (WebPlotDigitizer, https://automeris.io/), whose intercoder reliability and validity for this purpose have been formally established [57], percent changes are inherently more robust than absolute values to the small imprecision of graphical data extraction, as extraction error in numerator and denominator partially cancels out in the ratio. Overall, consistent with the exploratory and methodologically heterogeneous nature of the synthesized literature, our interpretation focuses on the direction and ordinal magnitude of effects rather than on exact point estimates.

**Table 1.** Results of the perturbation studies.

| Study | Measurement | Perturbation | Short-term DE change | Long-term DE change |
| --- | --- | --- | --- | --- |

|  | method |  | with perturbation | with perturbation |
| --- | --- | --- | --- | --- |
| Manor et al. ID 15 [67] | 2D motion capture, sagittal plane hip, knee, and ankle angles. | Ice-induced plantar desensitization | **+8%** | **+44%** |
| McAndrew et al. ID 22 [68] | 3D motion capture, 3D velocity of the cervical spine. | Mechanical and visual | Mechanical:<br>**+10% to +72%**<br>Visual:<br>**+3% to +86%** | Visual:<br>**-7% to -51%**<br>Mechanical:<br>**+5% to -38%** |
| Van Schooten et al. ID 26 [49] | 3D inertial sensor, accelerations and angular velocities of the lower back. | Vestibular | **+8% to +10%** | **-4% to -20%** |
| Sinitksi et al. ID 30 [69] | 3D motion capture, mediolateral velocity of the cervical spine. | Mechanical and visual | Mechanical:<br>**+30% to +34%**<br>Visual:<br>**+19% to +34%** | Mechanical:<br>**-27% to -30%**<br>Visual:<br>**-9% to -35%** |
| Rábago et al. ID 42 [70] | 3D motion capture, mediolateral velocity of the cervical spine. | Cognitive, mechanical, and visual | Cognitive:<br>**-3% to 0%**<br>Mechanical:<br>**+8% to +28%**<br>Visual:<br>**+10% to +30%** | Cognitive:<br>**+6% to -3%**<br>Mechanical:<br>**-16% to -39%**<br>Visual:<br>**-17% to -41%** |
| Reynard and Terrier ID 43 [71] | 3D accelerometer. Mediolateral acceleration of the upper trunk (sternum). | Vision removal | **-2% to +7%** | **0% to -20%** |
| Franz et al. ID 41 [33] | 3D motion capture. Right and left heel positions relative to the sacrum position. | Visual | Young:<br>**+6%**<br>Older:<br>**+22%** | Young:<br>**-13%**<br>Older:<br>**-17%** |
| Qiao et al. ID 47 [72] | 3D motion capture, 3D velocity of the cervical spine. | Visual | Healthy young controls:<br>**+19% to +24%**<br>Elderly non-fallers:<br>**+20% to +23%**<br>Elderly fallers:<br>**+19% to +28%** | Healthy young controls:<br>**-7% to -32%**<br>Elderly non-fallers:<br>**+2% to -49%**<br>Elderly fallers:<br>**-7% to -44%** |

Perturbation types included sensory manipulation (ice-induced plantar desensitization), mechanical disturbances (platform translations), visual alterations (moving room, optic flow), vestibular stimulation (galvanic stimulation), and cognitive tasks (dual-task paradigms). Percentage changes represent relative differences between perturbation and baseline conditions. Positive values indicate increased divergence exponent (DE) while negative values indicate decreased DE. Ranges reflect different measurement axes, experimental groups, or perturbation levels within individual studies. ID refers to study identification number in the systematic review database (S1 text in supplementary material).

**Table 2.**
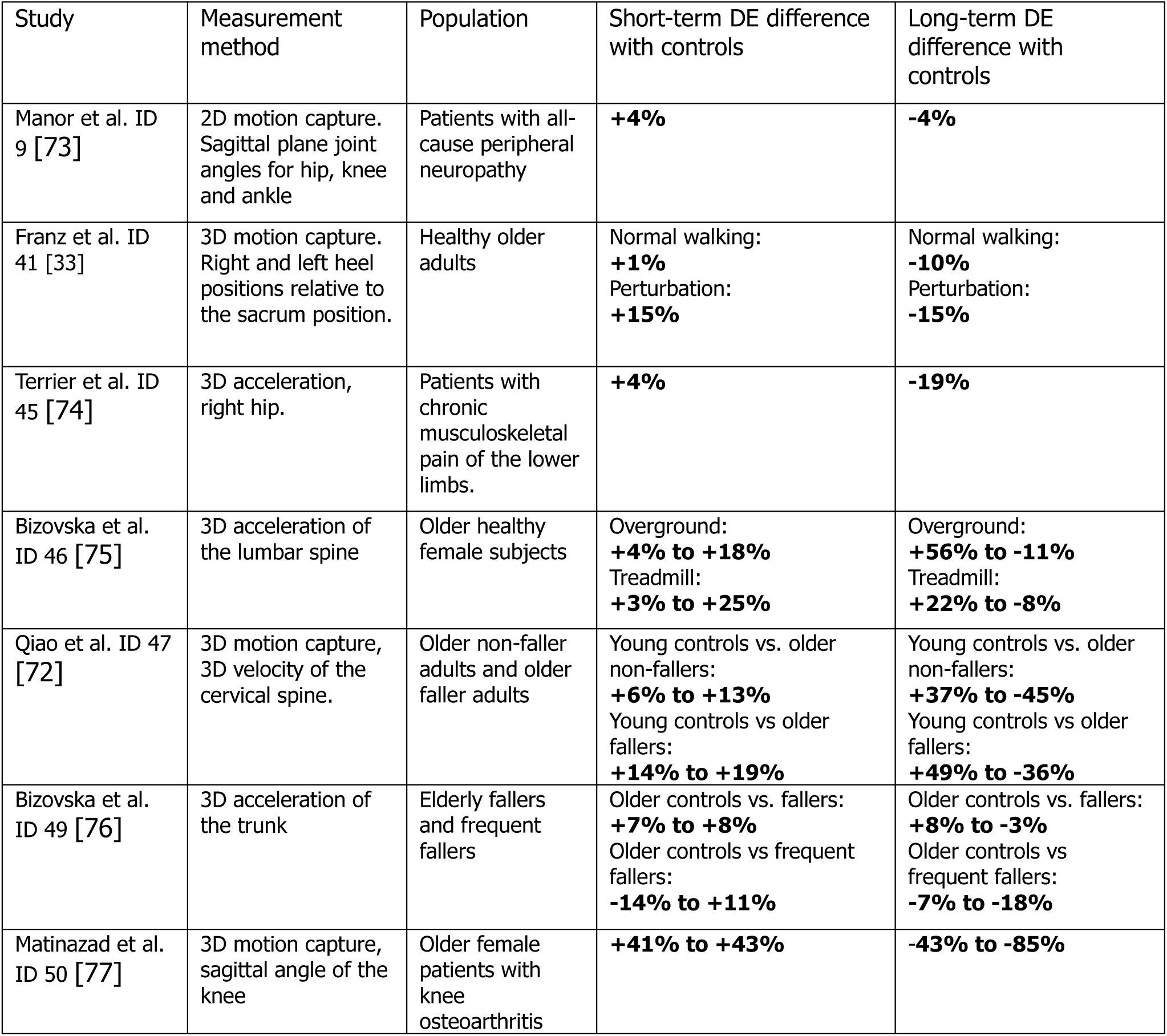
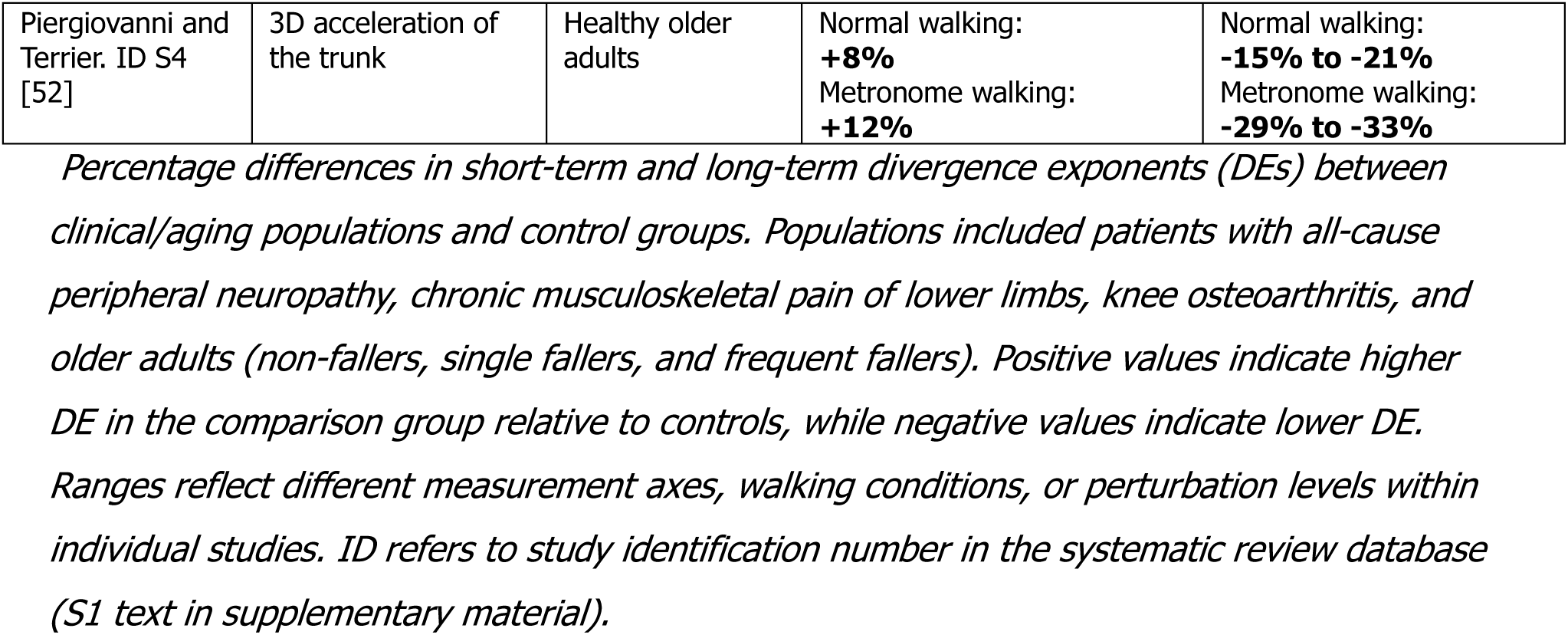
Results of the between-subject studies.

**Table 3.** Results of the externally paced walking studies.

| Study | Measurement method | Cueing | Short-term DE change with cueing | Long-term DE change with cueing |
| --- | --- | --- | --- | --- |
| Dingwell & Cusumano ID 3 [65] | Electrogoniometers (hip, knee, ankle) + sternum tri-axial accelerometer. | Treadmill speed constraint (vs overground). | Trunk acceleration: **-11% to -13%**<br>Lower limb kinematics: **-11% to -16%** | Trunk acceleration: **-11% to +12%**<br>Lower limb kinematics: **-11% to -20%** |
| Terrier & Dériaz ID 25 [58] | 3D accelerometer on lower back (lumbar), trunk accelerations. | Treadmill speed constraint (vs overground). | **-8% to -9%** | **-5% to -18%** |
| Sejdić et al. ID 29 [48] | Force-sensitive insole (heel strikes) + lower-back 3D accelerometer, trunk accelerations. | Rhythmic sensory cues during overground walking, step duration constraint: auditory, visual, and tactile. | Visual / step duration: **-4% to +3%**<br>Auditory / step duration: **-11% to -4%**<br>Tactile / step duration: **-8% to 0%**<br>Combined / step duration: **-12% to -8%** | Visual / step duration: **-10% to -18%**<br>Auditory / step duration: **-26% to -40%**<br>Tactile / step duration: **-15% to -23%**<br>Combined / step duration: **-34% to -43%** |
| Terrier & Dériaz ID 35 [40] | Instrumented treadmill (foot-pressure array); center-of-pressure trajectory. | Rhythmic auditory cueing (metronome), step duration constraint, during treadmill walking. | **-6% to +7%** | **-61% to -86%** |
| Bizovska et al. ID 46 [75] | Lumbar 3D accelerometer, trunk accelerations. | Treadmill speed constraint vs overground. | Young: **-19% to -24%**<br>Older: **-16% to -21%** | Young: **+24% to +156%**<br>Older: **0% to + 100%** |
| Terrier ID 51 [59] | Instrumented treadmill with force platform; center-of-pressure trajectory. | Visual (step length constraint) and auditory (step duration constraint) synchronization cues, during treadmill walking. | Visual / step length: **-4% to -2%**<br>Auditory / step duration: **-3% to + 14%** | Visual / step length: **-37% to -66%**<br>Auditory / step duration: **-43% to -78%** |
| Piergiovanni and Terrier. ID S4 [52] | 3D acceleration of the trunk | Rhythmic auditory cueing (metronome), step duration constraint, during overground walking. | Young: **+1%**<br>Older: **-3%** | Young: **-13% to -15%**<br>Older: **-26% to -30%** |

*Percentage changes in short-term and long-term divergence exponents (DEs) during* *externally paced walking compared to self-paced conditions. Interventions included treadmill* *speed constraint (imposing constant velocity versus self-paced overground walking) and* *rhythmic sensory cueing (auditory metronome, visual pacing, tactile stimulation). Negative* *values indicate reduced DE with external pacing, while positive values indicate increased DE.* *Ranges reflect different cueing modalities, measurement signals, or experimental conditions* *within studies. Combined cueing indicates simultaneous presentation of multiple sensory*
modalities. ID refers to study identification number in the systematic review database (S1 Text in supplementary materials).

#### Divergence curve characterization

In addition to extracting numerical outcomes, we systematically identified studies that published graphical representations of divergence curves. Divergence curves provide a visual representation of how the DEs are calculated. For each study that published divergence curves, we documented the figure location and extracted relevant information about the visual presentation of the method. For articles available through open-access licenses, we reproduced the published figures along with their complete legends. For studies behind paywalls where figure reproduction was not allowed, we documented the figure legends and descriptions to catalog the types of divergence curve presentations.

#### Quality assessment

We assessed study quality using a structured evaluation framework across three domains. Each study was scored on a 15-point scale comprising Domain 1: analytical rigor, 8 points; Domain 2: outcome reporting, 4 points; and Domain 3: sample size, 3 points. Domain 1 assessed methodological transparency through four criteria: experimental protocol description, measurement system reporting, signal processing clarity, and algorithm application details. Domain 2 evaluated the quality of statistical reporting, specifically whether descriptive statistics were clearly reported, and whether appropriate statistical analyses with effect sizes were provided. Domain 3 considered sample size adequacy relative to each study’s research question. This structured approach allowed for systematic comparison of methodological quality across the corpus while recognizing the exploratory nature of the field and the developmental stage of long-term DE methodology. Item-level and total quality scores for all included studies are provided in S3 Table (XLSX).

#### Synthesis method: meta-analysis

We identified five studies (contributing six independent study sets) that examined associations between long-term DE and scaling exponent (DFA) [40,50,52,58,59] for synthesis via meta-analysis. Among the studies included in the systematic review, two publications [51,52] used overlapping samples: the first reported correlation results for 60 older adults, while the second extended the analysis to include 42 additional younger participants. Because the older adult sample in [51] is entirely subsumed within the larger dataset of [52], only the latter was included in the meta-analysis. These studies reported multiple correlation coefficients, corresponding to different measurements (accelerometer axes, body positions). Therefore, to synthesize their results, we performed a meta-analysis using a multilevel random-effects approach in accordance with current methodological recommendations for handling dependent effect sizes [60]. We implemented a correlated and hierarchical effects model using the metafor package in R [61]. This model accounts for both between-study heterogeneity and within-study correlation structures. We assumed within-study correlation of 0.8 between effect sizes from the same study, as the different correlation coefficients consisted of repeated measures in the same subjects. Within each study, repeated correlation coefficients were aggregated to a single study-level estimate using a generalized least-squares average that respects the assumed within-study correlation. Robust variance estimation (RVE) with small-sample correction was applied using the clubSandwich package [62] to account for potential misspecification of the working model and to generate valid inferences despite the dependence structure of the data. Fisher’s Z-transformed correlations were used for analysis and back-transformed to correlation coefficients for interpretation. We also estimated heterogeneity and summarized it with I². Results were visualized using forest plots displaying individual and overall estimates with their respective 95% confidence intervals.

#### Protocol deviations

We acknowledge five protocol deviations that occurred during the conduct of this review. First, the eligibility criteria for study inclusion originally specified a publication date range from 2001 to 2024. However, it became apparent that seminal articles relevant to the analysis of DE in nonlinear gait analysis were published as early as 2000. Consequently, the inclusion criteria were adjusted to include publications from 2000 to 2024. Second, although not explicitly stated in the original protocol, we decided during the data extraction phase to identify and report studies using the detrended fluctuation analysis (DFA) method in parallel to long-term DE. Third, we systematically documented divergence curve presentations to capture how these visualizations demonstrate the extraction of short-term and long-term DEs and reveal characteristics of the underlying gait dynamics. Fourth, the protocol specified a narrative quality assessment approach. However, upon initiating the assessment process, we recognized that a more structured framework would facilitate systematic evaluation and comparison across studies. We therefore developed a formal quality assessment grid with explicit scoring criteria across three methodological domains, enabling quantitative characterization of reporting quality while maintaining the focus on methodological transparency rather than risk of bias. Fifth, an updated search was conducted in January 2026 to capture studies published after the original search date of February 2024. This update, which was not anticipated in the protocol, was motivated by the extended time required to complete data extraction, quality assessment, manuscript preparation, and the publication process.

## Results

Our systematic search retrieved 673 records from database queries and 3 additional records from references of other articles, yielding 676 unique records for title-abstract screening. Of these, 242 were excluded: 102 for non-relevant sources (proceedings, reviews), 51 for modeling studies, and 89 for irrelevant topics or non-walking gaits (running). Among the excluded records, 4 articles published in languages other than English or French (two Spanish, one German, one Chinese) underwent screening based on English abstract; none met the inclusion criteria. This left 434 full-text articles for eligibility assessment, all of which were in English. The full-text review resulted in the exclusion of 378 articles, mainly due to reporting only short-term DE. Other secondary reasons for exclusion were the use of alternative DE algorithms (Wolf) or other stability metrics.

An updated search conducted in January 2026 retrieved 94 additional records from the same databases. Following the same screening procedure, 36 records were excluded at the title-abstract stage (9 for non-relevant sources, 7 for modeling studies, and 20 for irrelevant topics or non-walking gaits), and 58 full-text articles were assessed for eligibility. Of these, 52 were excluded, and 6 new studies met the inclusion criteria, one of which contributed data to the meta-analysis.

In total, the combined searches yielded 770 records, of which 62 studies were included in the outcome synthesis and 5 in the meta-analysis (Fig 1).

### Methodological outcomes

#### Study characteristics, methodological scope, and sample sizes

Fig 2 summarizes the characteristics of the 62 included studies, based on the data provided in the supporting information (Table B in S1 Text). Panel a illustrates the annual number of publications from 2000 to 2026, showing a consistent upward trend starting in 2008, followed by sustained activity through the mid-2010s and a resurgence of four publications in 2024. Panel b categorizes the type of research by participant category. Exact definitions for each research type are provided in supporting information (Table A in S1 Text). In terms of populations studied, studies involving only healthy adults predominate, with 42 studies. Within this category, most studies compared gait patterns under different experimental conditions and aimed to develop, validate, or assess the reliability of DE methods. Research on age-related gait changes and fall risk comprises 14 studies, while the remaining 6 studies investigated gait patterns associated with other pathologies. Only a few studies have used long-term DE for clinical investigations (4). Only one study was interventional and longitudinal, i.e. it analyzed the gait patterns of patients before and after surgery. Aside from studies aimed at predicting fall risk, we identified no studies that evaluated the use of long-term DE as a diagnostic tool for detecting gait abnormalities. Panel c shows that across the study samples, the median sample size was 23 participants (interquartile range 13-36), with most studies enrolling fewer than 50 individuals; only a handful exceeded 100, and the full range extended from 9 to 187 participants, indicating a generally modest sample size framework in the field. Note that one article was counted as two different studies because it reported two separate experiments with two different groups of participants.

**Fig 2.**
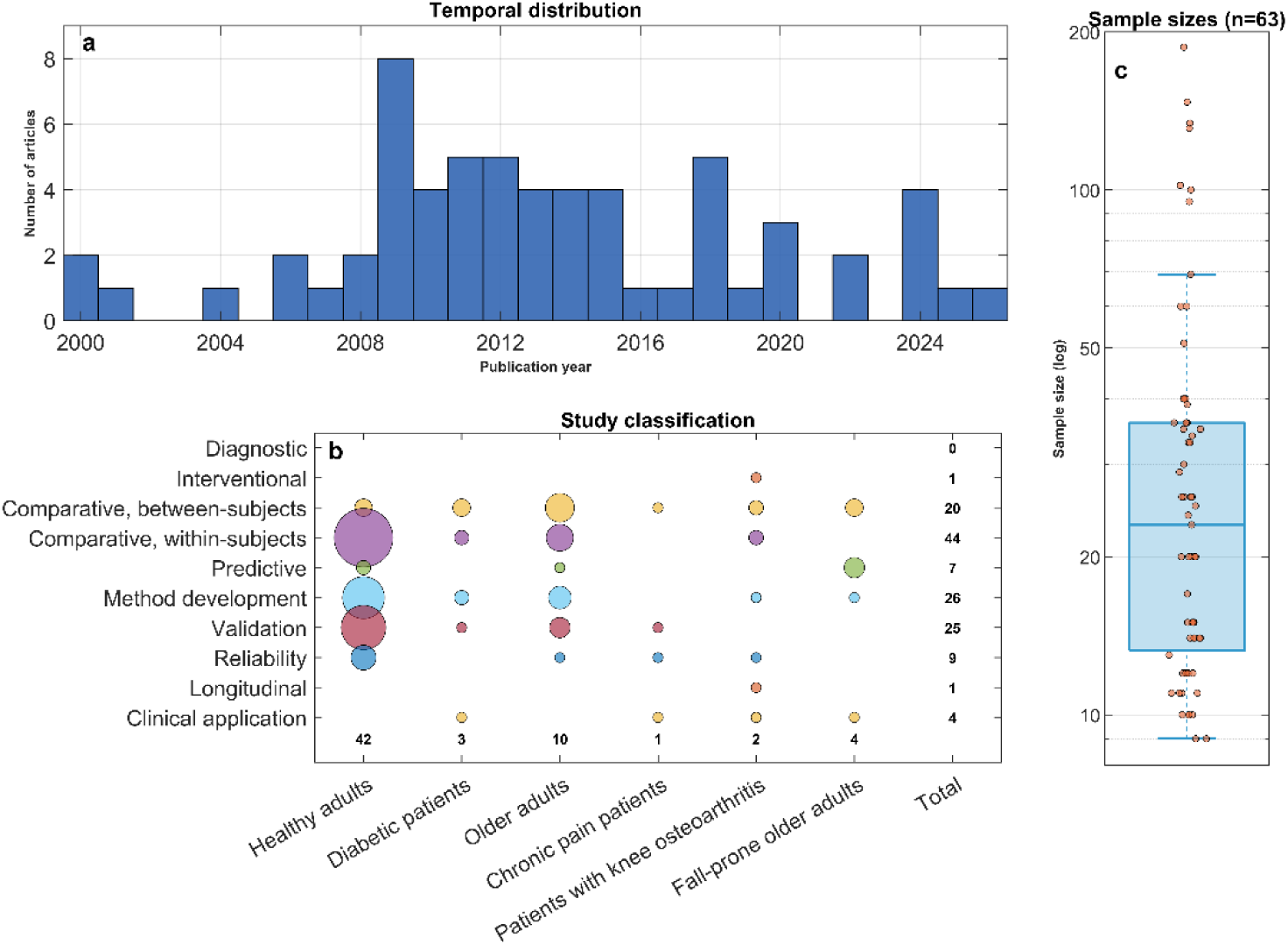
Study characteristics. **a)** Annual publication trends. b) Distribution by research type and participant category. c) Sample size distribution (n = 63 because one study reported two independent experiments with separate participant groups). DE, divergence exponent.

#### Experimental settings and sensor modalities

Per-study methodological outcomes for all 62 studies are provided in Table C in S1 Text. Measurement environments were overwhelmingly laboratory-bound (Fig 3a). Forty-two of the 62 studies (68%) acquired data on a motorized treadmill in an indoor laboratory setting, typically in conjunction with 3-D motion-capture systems. Thirteen studies (21%) relied solely on overground walking, but even here the majority preserved a controlled indoor context (indoor circuits or short walkways). Only three overground protocols departed from this paradigm by recording prolonged outdoor or quasi-natural bouts. Six further articles (10%) combined treadmill and overground sessions within the same experiment, whereas one study (2%) captured free-living gait during a week-long monitoring campaign. Instrumentation choices mirrored these environmental constraints (Fig 3b). Optical motion-capture was used in 31 of the 62 studies (50%). Wearable inertial measurement units (IMUs) ranked second, appearing in 19 studies (31%). Dedicated instrumented treadmills (equipped with embedded force or pressure arrays) were the unique measurement device in 3 studies (5%). Although single-sensor protocols remained the norm, a minority of nine studies (15%) employed hybrid instrumentation, most commonly electrogoniometers coupled with a trunk accelerometer (4/9, 44%), followed by optical motion-capture combined with body-worn IMUs (3/9, 33%) and motion capture and surface electromyography (EMG, 2/9, 22%). Most studies prioritized measurement at the trunk level (Fig 3c): 34 of the 62 studies (55%) analyzed pelvis, lumbar or thoracic spine kinematics. Lower-limb–only set-ups (hip, knee, ankle kinematics or EMG) appeared in 11 studies (18%). Eleven studies (18%) combined trunk and lower-limb measurements within the same experiment. Foot-pressure–only configurations were rare (3/62, 5%), corresponding to the instrumented-treadmill studies. Other designs include head, upper-limb, or composite whole-body recordings (3/62, 5%).

**Fig 3.**
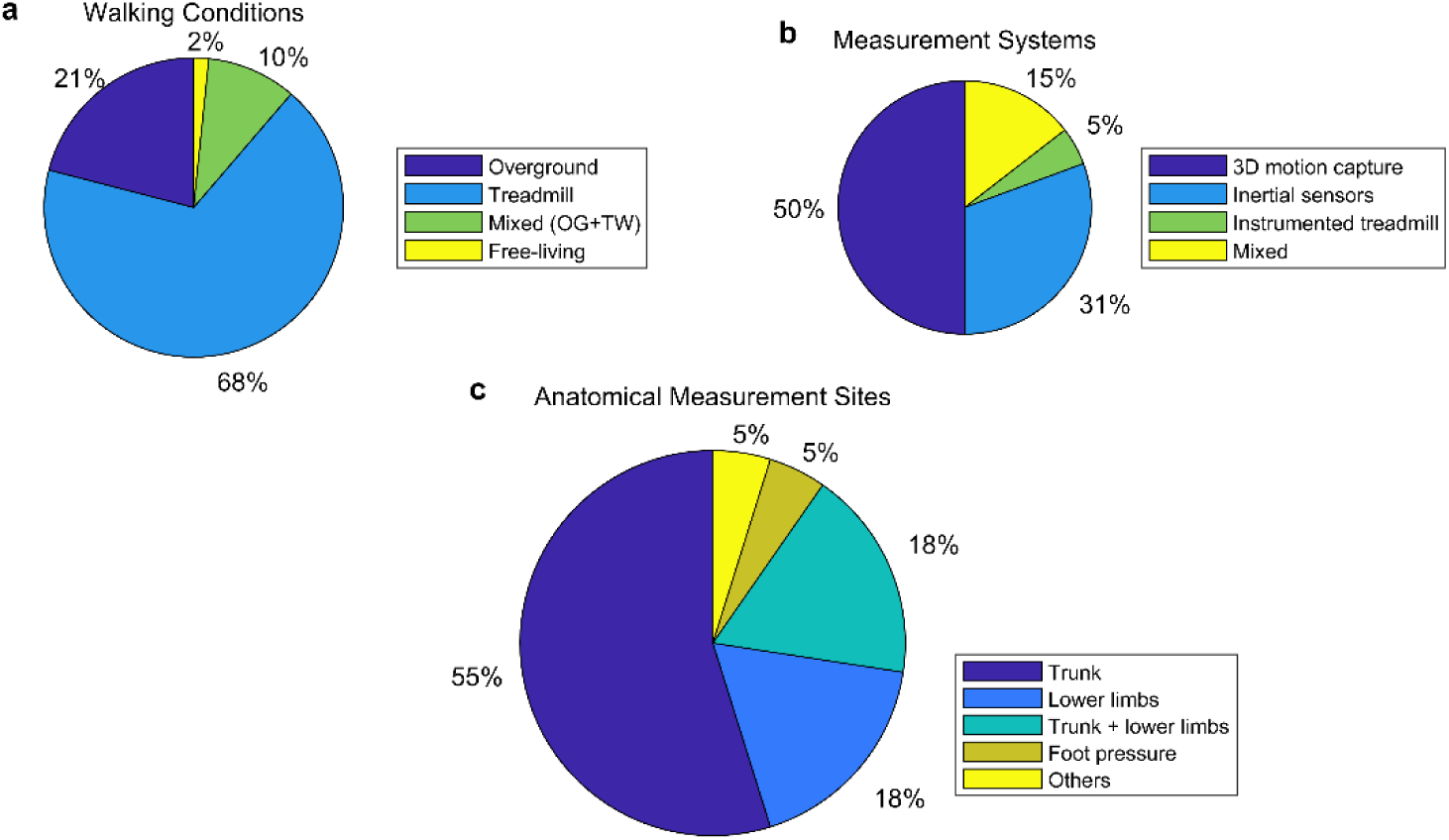
Methodological characteristics. **a)** Walking conditions. **b)** Measurement systems. **c)** Anatomical measurement sites. IMU, inertial measurement unit.

#### Measurement lengths

Because Rosenstein’s algorithm requires a single continuous signal, we extracted the duration corresponding to one analyzed walking bout per experimental condition, rather than aggregating multiple bouts. Reporting practices across the included studies varied substantially: approximately one-third specified bout length exclusively in minutes, another third exclusively in strides, and the remainder provided both units. Due to this methodological heterogeneity, direct quantitative synthesis of durations was not feasible. We therefore adopted an ordinal categorization to harmonize duration data into four empirically defined classes (Fig 4). The threshold for the “very short” category (≤ 2 min or ≤ 100 strides) was anchored to the median cadence observed in studies reporting both units (∼55 strides·min⁻¹), yielding approximately 100 strides over two minutes—a practical lower limit below which long-term DE estimates may become unreliable [63]. The upper limit for the “short” category (> 2–5 min or 101–250 strides) aligns with the most commonly adopted experimental protocol across studies (3–5 minutes of treadmill or overground walking). The “moderate” category (> 5–10 min or 251–500 strides) encompasses studies employing somewhat longer walking bouts, while the “long” category (> 10 min or > 500 strides) includes the few studies focused on endurance-oriented walking conditions. Applying this ordinal scale to the extracted data resulted in the following distribution: “short” (30 studies; 48%), “very short” (22 studies; 35%), “moderate” (6 studies; 10%), and “long” (4 studies; 6%). Finally, five studies computed DE from averaged signals of multiple short walking bouts within the same experimental condition; for these, we categorized duration based on a single constituent bout, although averaging multiple bouts may enhance the reliability of DE estimates [63].

**Fig 4.**
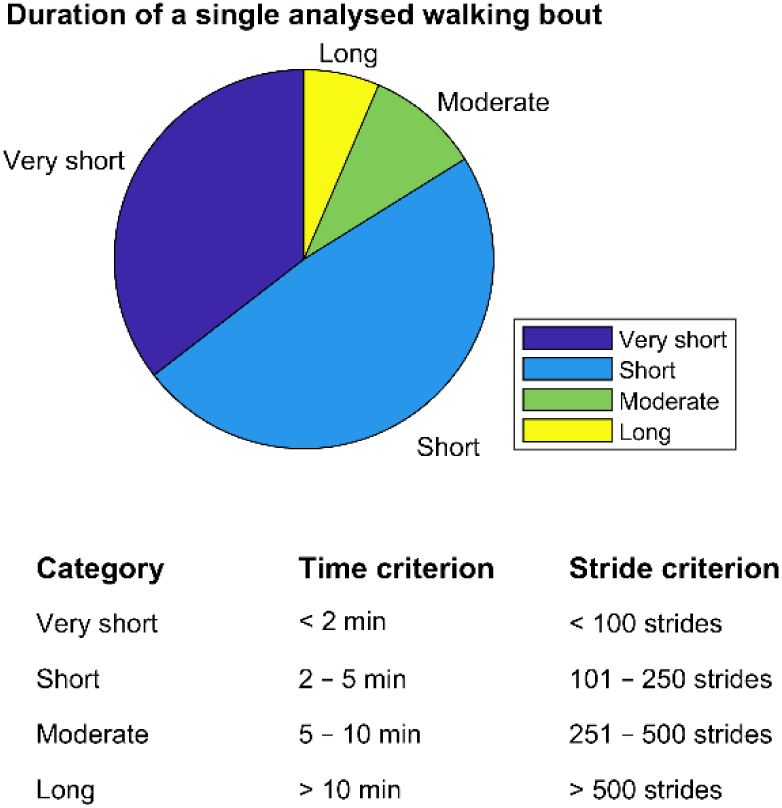
Measurement duration. Distribution of walking bout durations across four duration categories.

#### Signal processing

Signal sources driving the computation of long-term DE were clustered into distinct categories (Fig 5a). Half of the studies (31 out of 62, 50%) utilized kinematic data exclusively, acquired via optical motion capture or electrogoniometers. Within these kinematic studies, angular joint trajectories (joint angles) were predominant, representing most cases. Additionally, linear positional data (e.g., cervical or thoracic spine positions) and derived velocity data were also commonly used. Accelerometric data from inertial sensors constituted the second most frequent signal type, with 18 studies (29%) employing linear accelerations exclusively. Only two studies (3%) incorporated combined inertial measurements of linear acceleration and angular velocity from gyroscopes. No studies used gyroscopic (angular velocity) signals alone. Three studies (5%) derived DEs from COP trajectories obtained from instrumented treadmills. Surface EMG voltage signals alone were used in a single study (2%). Mixed signal sources appeared infrequently: six studies (10%) combined joint-angle kinematics with trunk acceleration, and one study (2%) combined whole-body kinematics with multi-channel EMG. Reporting of raw-signal filtering was inconsistent across the corpus. Twenty-one of the 62 studies (34%) explicitly described applying a digital filter before computing the long-term DE, almost always a zero-lag low-pass Butterworth between 8 Hz and 20 Hz. Twenty-two papers (35%) stated unambiguously that no filtering was performed. For the remaining nineteen studies (31%) no filtering procedure was mentioned, making it probable, though not certain, that the raw signals were analyzed without additional pre-processing.

**Fig 5.**
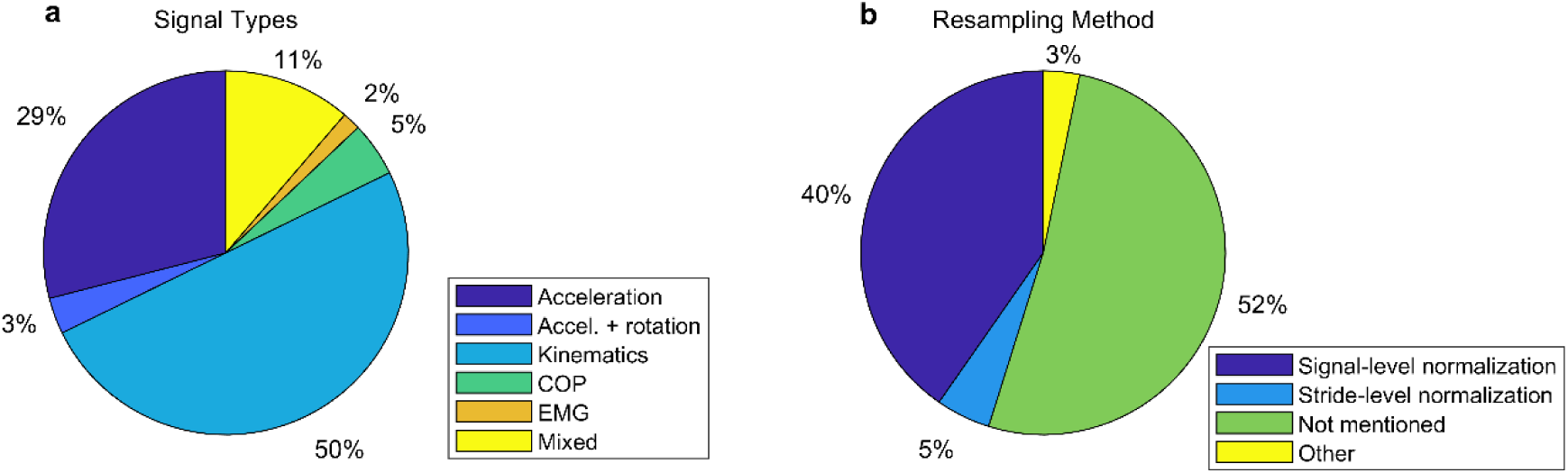
Signal processing characteristics. **a)** Signal sources used for computing DEs. **b)** Resampling methods to normalize signal durations. DE, divergence exponent; COP, center of pressure; EMG, electromyography.

The next steps in DE computation typically involve time normalization of raw data. The rationale is to maintain a consistent number of steps and data points across participants and conditions. This approach prevents potential bias arising from variations in DE values associated with sample length [63]. Panel b of Fig 5 shows the handling of temporal resampling. Explicit time normalization, typically realized as linear or spline interpolation to a constant number of samples, was reported in 25 of the 62 investigations (40%). Three studies (5%) used an alternative method consisting in resampling separately each stride to a fixed number of samples. In contrast, 32 papers (52%) gave no information on resampling, very likely using raw, unequal-length signals, and / or signals with a variable number of steps. The remaining two articles (3%) explicitly did not perform time normalization.

#### Algorithms

The subsequent processing involves reconstructing the state-space on which DE is computed. Among the 62 studies, 54 (87%) used the classical Takens approach (see Appendix A in [40]), generating a *d_E_-*dimensional trajectory from a single signal and its *T*-delayed copies [64]. By contrast, 8 studies (13%) constructed ad-hoc, multi-channel attractors by concatenating measured variables (typically linear plus angular trunk kinematics, or acceleration combined with gyroscope signals) before any delay embedding. Within the subset of studies using the classical approach (54 studies), the tuning procedure for the delay *T* and embedding dimension *d_E_* was reported heterogeneously. The standard pair (average mutual information (AMI) for *T* and global false-nearest-neighbors (GFNN) for *d_E_*) was used in 34 studies (63%). Seven studies (13%) substituted the autocorrelation criterion for *T* while still using GFNN, and the remaining thirteen (24%) gave no explicit or unclear justification for either parameter.

Finally, after reconstructing the state-space and computing divergence curves, the long-term DE is determined by performing a linear fit on a specific segment of the divergence curve [40]. The interval used for this linear fit was initially defined in seminal studies by Dingwell et al. [41,65]. Most studies (55 out of 62, 89%) adopted this canonical interval, 4 to 10 strides. Among the seven studies (11%) that selected alternative ranges, four used intervals between 0.5 and 10 strides (including one study that explored multiple sub-intervals), while three recent studies employed ranges shifted toward later strides (5–12 and 6–8 strides), two of which evaluated multiple range/length combinations to optimize responsiveness and reliability of the long-term DE estimate.

#### Divergence curve visualization

Divergence curves (graphical representations plotting the average logarithmic divergence of neighboring trajectories against time) demonstrate how short-term and long-term DEs are extracted as linear slopes from distinct temporal regions. In addition, their shape may allow visualizing when divergence ceases to grow as trajectories reach bounded regions of phase space. Among the 62 included studies, 30 (48%) published divergence curves in their figures. We systematically categorized the purpose for which divergence curves were presented across these studies. Eighteen studies (60%) used divergence curves exclusively for methodological illustration, displaying representative curves to demonstrate the calculation procedure without comparing experimental conditions. Seven studies (23%) employed divergence curves exclusively as comparative results, presenting curves to highlight differences between groups or experimental conditions. Five studies (17%) included both types of presentations, incorporating separate figures for methodological illustration and comparative outcomes. Among studies presenting comparative results, typical comparisons included different walking speeds, perturbation versus control conditions, young versus older adults, and with versus without external cueing. The compiled divergence curves and associated figure descriptions are provided in S2 Figures.

#### Quality Assessment

Quality assessment was completed using the three-domain scoring framework (maximum 15 points). Overall methodological quality was moderate to high, with a mean score of 11.56 (SD = 1.78, range 8-15) out of 15 possible points. Twenty-seven studies (44%) achieved high quality (12-15 points), while 35 studies (56%) demonstrated moderate quality (8-11 points). No studies scored in the low-quality range (0-7 points). Complete quality assessment scores for all studies are provided in S3 Table.

Domain-specific analysis revealed distinct patterns across methodological areas. Domain 1 (analytical rigor) demonstrated the strongest performance, with studies averaging 7.60 ± 0.66 out of 8 points (95%), and 43 studies (69%) achieving perfect scores (8/8). This indicates robust methodological rigor in experimental protocols, measurement systems, signal processing, and algorithm implementation across the included literature. In contrast, Domain 2 (outcome reporting) showed moderate performance (mean 2.31 ± 1.15 out of 4 points; 58%), with only 13 studies (21%) achieving full credit for comprehensive statistical reporting. Domain 3 (sample size) similarly revealed limitations, averaging 1.66 ± 0.81 out of 3 points, with most studies (40%) demonstrating moderate sample size justification and only 10 studies (16%) providing adequate sample size justification.

### Long-term DE results

#### Perturbation studies

Among the 62 studies, 14 studies experimentally destabilized gait using a range of perturbation modalities, including mechanical disturbances (e.g., compliant surfaces, outdoor surfaces, segmental loads, unexpected slips, platform instability), sensory manipulations (e.g., galvanic vestibular stimulation, blindfolding, optical-flow distortions, foot desensitization), or cognitive disturbance (dual-tasking). By design, these studies compare each participant to themselves across conditions, holding measurement system, signal processing, and algorithm parameters constant; between-condition differences in DE values therefore reflect the response to the experimental manipulation itself. In addition, these experimental paradigms are particularly relevant for testing the hypothesis that long-term DE reflects the degree of gait automaticity. When gait is perturbed or destabilized, it is reasonable to assume that individuals allocate greater attentional resources to gait control in order to manage the imposed challenges. However, none of the 14 perturbation studies explicitly tested this hypothesis, as they interpreted long-term DE in the same manner as short-term DE, that is, as a direct index of gait stability sensitive to the magnitude of external perturbation. We summarize the results of the retained perturbation studies in Table 1. Because long-term DE is sensitive to signal length (number of data points or strides), valid comparisons require similar data lengths across conditions. In treadmill perturbation paradigms, where walking speed is held constant between baseline and challenge, data length remains comparable even without time normalization. Therefore, we excluded studies with very-short bouts or overground designs in which walking speed (and thus signal length) differed between conditions, retaining only treadmill experiments with sufficient bout duration and matched speeds. We also excluded one study that used stride-level temporal normalization [66] which suppresses long-term divergence [53] and renders values non-comparable to those obtained with signal-level resampling (Fig. 5b). Note that the study by Manor et al. analyzed a reduced number of consecutive strides (45), but reported results averaged across three different walking speeds, which may enhance the reliability of the DE assessment.

Study designs differed in the nature and number of perturbation conditions, which explains why we report ranges rather than single effect sizes. For most studies, numeric values were extracted from figures using WebPlotDigitizer (https://automeris.io/). Manor et al. [67] desensitized the plantar arch using ice to diminish sensory feedback from the feet, anticipating subsequent gait destabilization. Unlike other perturbation studies that analyzed trunk motion, Manor et al. focused on lower-limb kinematics. Their findings indicated an increase in both short-term (+8%) and long-term (+44%) DE values, with results aggregated across different walking speeds and joints. In McAndrew et al. [68], participants experienced both mediolateral and anteroposterior platform oscillations and visual-field perturbations while DE was computed along the AP, ML, and vertical axes, yielding short-term DE increases of 10–72% (mechanical) and 3–86% (visual) and long-term DE decreases of 5–38% and 7–51%, respectively. Sinitski et al. [69] systematically varied the amplitude of both mechanical and visual perturbations, observing short-term DE rises of 30–34% (mechanical) and 19–34% (visual) alongside long-term DE drops of 27–30% and 9–35%, respectively. Rábago et al. [70] compared cognitive, mechanical, and visual perturbations: cognitive load had negligible effects on short-term DE (−3% to 0%) with near-zero long-term changes (−3% to +6%), while both mechanical and visual perturbations produced a similar pattern of increased short-term DE (+8% to +30%) accompanied by decreased long-term DE (−16% to −41%). Van Schooten et al. [49] and Reynard & Terrier [71] compared walking at multiple speeds under galvanic vestibular stimulation and blindfolded conditions. They found short-term DE increases of 8–10% (GVS) and –2 to +7% (blindfold), and long-term DE decreases of 4–20% and 0–20%, respectively. Franz et al. [33] analyzed younger and older subjects, reporting short-term DE increases of 6% (young) and 22% (older), and long-term DE decreases of 13% (young) and 17% (older) under visual perturbation; they also found a reduction in the scaling exponent reflecting gait complexity (DFA method) by 6% in young and 26% in older participants. The study by Qiao et al. assessed the effects of varying intensities of optical-flow distortions on young and older adults. Here again, opposite responses were observed between short-term DE, which increased (+19% to +28%), and long-term DE, which mostly decreased (+2% to −48%). Across all studies reviewed, the majority of perturbation-induced changes were statistically significant (p < 0.05), though certain exceptions occurred, notably for long-term DE measures that exhibited greater inter-individual variability.

#### Between-subject studies

Fifteen between-subject studies compared cohorts with presumed differences in gait stability and potentially in gait automaticity, including clinical groups (e.g. diabetic or peripheral neuropathy, knee osteoarthritis, chronic pain, fallers vs non-fallers) and demographic cohorts (healthy older vs younger adults, inexperienced vs experienced military cadets). As above, these are within-study contrasts that hold measurement and analytical parameters constant across compared groups, although inter-individual variability reduces statistical power relative to repeated-measures designs.

However, most of these studies based their DE analysis on too short walking bouts or on non-time normalized signals. After careful evaluation of study methodologies, eight studies met the inclusion criteria for a valid comparison (Table 2).

Although Terrier et al. analyzed very short walking bouts (35 strides, 1 min), they aggregated numerous bouts collected over one week under free-living conditions (on average 77 bouts in patients and 34 in healthy controls). This aggregation enhances the reliability of the average DE values, mitigating concerns related to short individual signal durations.

Study methodologies differed in signal processing approaches and reported outcomes. Manor et al. [73] averaged their analyses across joint angles and walking speeds, reporting a single DE value per group (patients with peripheral neuropathy vs. healthy controls). Similarly, Terrier et al. [74] computed the vector magnitude (norm) from three axes of hip acceleration, yielding a single representative DE per group (chronic musculoskeletal pain patients vs. controls). Using 3D motion capture during treadmill walking, Franz et al. [33] examined trunk motion responses to visual perturbations. Similarly, the visual perturbation study of Qiao et al. [72] examined cervical spine velocity under three increasing perturbation levels. Bizovska et al. [75] reported separate DE outcomes for each axis of a 3D accelerometer under two distinct walking conditions (overground vs. treadmill), resulting in a range of observed differences between older and younger healthy females. Similarly, Piergiovanni & Terrier [52] reported long-term DE across three axes of a lower-back accelerometer under normal and metronome walking conditions, comparing older to younger adults. In their second study, Bizovska et al. also analyzed DEs across accelerometer axes, comparing a control group of healthy older adults with groups of older individuals who fell infrequently or frequently. Finally, Matinazad et al. studied knee kinematics at two different speeds.

Short-term DE demonstrated a pattern of limited increases in most populations compared to their respective controls, though with heterogeneity in magnitude. The most pronounced differences were observed in patients with knee osteoarthritis (41-43% higher). Clinical populations with neuropathy and chronic musculoskeletal pain showed consistent increases compared to controls (+4%). Regarding age comparison studies, Franz et al. reported minimal differences during normal walking (1% higher in older adults as compared to younger controls) but substantially larger increases under visual perturbations (15% higher in older adults). Piergiovanni & Terrier [52] found a similar pattern, with 8% higher short-term DE in older adults during normal walking and 12% higher during metronome walking. Qiao et al. [72] found higher short-term DE in older adults versus younger adults, with this difference amplified by perturbation intensity, especially in older fallers. Bizovska et al. [75] found short-term DE differences ranging from 4% to 18% higher in older compared to younger females during overground walking, and from 3% to 25% higher during treadmill walking. However, it is worth noting that the older group had a mean age of only 57.5 years (±4.8), representing middle-aged rather than elderly adults. In their second study focused on fall risk, Bizovska et al. [76] observed short-term DE increases of 7-8% when comparing elderly non-fallers to single fallers, while comparisons between non-fallers and frequent fallers showed highly variable results ranging from 14% decreases to 11% increases depending on measurement axis.

In contrast to the relatively consistent short-term increases, long-term DE exhibited heterogeneous patterns. The most substantial reductions were observed in clinical populations: patients with knee osteoarthritis showed decreases ranging from 43% to 85% compared to controls, while those with chronic musculoskeletal pain demonstrated 19% reductions. Neuropathy patients showed minimal reductions of 4%. Franz et al. [33] reported consistent long-term DE reductions in older adults during both normal walking (−10%) and under visual perturbations (−15%). Piergiovanni & Terrier [52] found comparable reductions during normal walking (−15% to −21% across axes), which were further amplified during metronome walking (−29% to −33%). Despite reporting heterogeneous results, Qiao et al.’s figures [72] show a trend toward lower long-term DE (−36%) in older participants under high visual perturbation. The two studies by Bizovska et al. revealed higher variability depending on measurement axis and experimental conditions. In their age-comparison study [75], long-term DE differences ranged from increases of 56% to decreases of 11% when comparing older and younger females during overground walking, and from increases of 22% to decreases of 8% during treadmill walking. Their fall-risk study [76] showed long-term DE differences ranging from increases of 8% to decreases of 3% when comparing non-fallers to single fallers, and from decreases of 7% to 18% when comparing non-fallers to frequent fallers. Overall, three of the four age-comparison studies showed consistent long-term DE reductions in older adults; only the Bizovska data, based on a relatively young older cohort (mean age 57.5 years), diverged from this pattern.

#### External cueing studies

Among the 62 studies, seven within-subject experiments examined either externally imposed gait synchronization or a fixed-speed treadmill constraint. As in the perturbation studies above, this design ensures that between-condition differences in DE values reflect the experimental manipulation itself rather than variation in measurement system, signal processing, or algorithm parameters. As these interventions likely shift the balance between automatic and voluntary control of walking [78], they enable evaluation of the sensitivity of long-term DE to variations in gait automaticity. Three studies implemented synchronization cues: Sejdić et al. [48] applied visual, auditory, tactile, and combined step-duration cues during overground walking using heel-strike detection with a force-sensitive insole and lumbar accelerometry; Terrier & Dériaz [40] and Terrier [59] imposed metronome-based step-duration or visual step-length targets during treadmill walking while recording center-of-pressure trajectories on an instrumented treadmill. Piergiovanni & Terrier [52] applied auditory metronome cueing during overground walking in both younger and older adults, recording trunk accelerations from a lower-back accelerometer. Three studies compared constant-speed treadmill with unconstrained overground walking: Dingwell & Cusumano [65] recorded trunk accelerations (sternum) and lower-limb joint angles via electrogoniometers, and Terrier & Dériaz [58] used lower-back accelerometry; Bizovska et al. [75] extended this comparison in young and older women with lumbar accelerometry. Trial durations typically ranged from 3 to 10 min per condition (yielding ∼100–500 strides), with speed held constant in cueing paradigms; overground–treadmill contrasts occasionally differed in bout duration or speed (e.g., 5 min overground vs 3 min treadmill in [75]).

Across studies that imposed external synchronization cues (auditory, visual, tactile), short-term DE changed little or inconsistently (approximately −11% to +14%), whereas long-term DE showed large, systematic reductions (≈ −10% to −86%). In the overground study using rhythmic sensory cues [48], short-term DE was minimally affected (visual: −4% to +3%; tactile: −8% to 0%) or modestly decreased under auditory and combined cueing (auditory: −11% to −4%; combined: −12% to −8%). By contrast, long-term DE fell substantially (visual: −10% to −18%; tactile: −15% to −23%; auditory: −26% to −40%; combined: −34% to −43%). Similarly, a substantial decrease in gait complexity measured from force-sensitive insoles (scaling exponent, DFA) was observed (−50% to −13%). In the second overground cueing study, Piergiovanni & Terrier [52] observed the same dissociation: auditory metronome cueing left short-term DE essentially unchanged (young: +1%; older: −3%) while producing substantial long-term DE reductions that were more pronounced in older adults (−26% to −30%) than in younger participants (−13% to −15%). Two treadmill experiments that enforced step timing/length converged on the same pattern. Using an instrumented treadmill and center-of-pressure (COP) trajectories, Terrier & Dériaz [40] reported very large long-term DE decreases (−61% to −86%) with only small, direction-dependent short-term changes (−6% to +7%). In a mixed visual–auditory cueing paradigm [59], short-term DE remained near baseline (visual/step-length: −4% to −2%; auditory/step-duration: −3% to +14%), whereas long-term DE dropped markedly for both constraints (visual/step-length: −37% to −66%; auditory/step-duration: −43% to −78%).

By contrast, comparing treadmill speed constraint with unconstrained overground walking yielded consistent short-term DE reductions but heterogeneous long-term responses across datasets and signal types. In three studies that contrasted fixed-speed treadmill with overground walking, short-term DE was lower on the treadmill in trunk accelerations ( [65]: −11% to −13%, [58]: −8% to −9%, and [75] −16% to −24%), and lower-limb kinematics ([65]: −11% to −16%). Long-term DE, however, varied: it mostly decreased in two datasets ([65]: trunk −11% to +12%, lower-limb −11% to −20%; [58]: −5% to −18%), but substantially increased in [75] (+24% to +156%).

#### Meta-analysis

We synthesized six independent datasets reporting the association between long-term DE and the scaling exponent from DFA. Measurement modalities spanned instrumented treadmills with center-of-pressure trajectories, trunk accelerometry, and segmental kinematics of the head and lower limbs; several experiments also varied speed (slow, preferred, fast) or imposed synchronization tasks. One publication contributed two distinct datasets because the same participants walked in separate overground and treadmill experiments [58]; these were treated as independent due to the large contextual difference between conditions. Where articles reported axis- or segment-specific associations, we retained those dependent effects (e.g., anteroposterior, mediolateral, vertical, or head vs ankle). This yielded 14 dependent correlations across the six datasets (k=14), while speed manipulations within a given protocol were not counted as separate effects when correlations were reported on the pooled observations [50].

Using a correlated-effects random-effects model with cluster-robust variance estimation, Fisher’s Z-transformed correlations were pooled and then back-transformed for interpretation. The overall association was positive, with a pooled correlation of 0.64 (95% CI 0.34 to 0.82) based on 14 effects (N=582 condition-level observations from 209 participants). Heterogeneity was considerable (I²≈82%), with estimated between-study variance τ²≈0.22 and within-study variance ω²≈0.011. Leave-one-out influence diagnostics showed that the pooled correlation remained stable in magnitude and direction. Re-estimating the model after omitting each dataset in turn yielded pooled r values between 0.57 and 0.71. Across all leave-one-out fits, the 95% confidence intervals stayed positive, and the standard error varied only modestly (0.17–0.24). The forest plot summarizes study-level estimates and the model summary (Fig 6).

**Fig 6.**
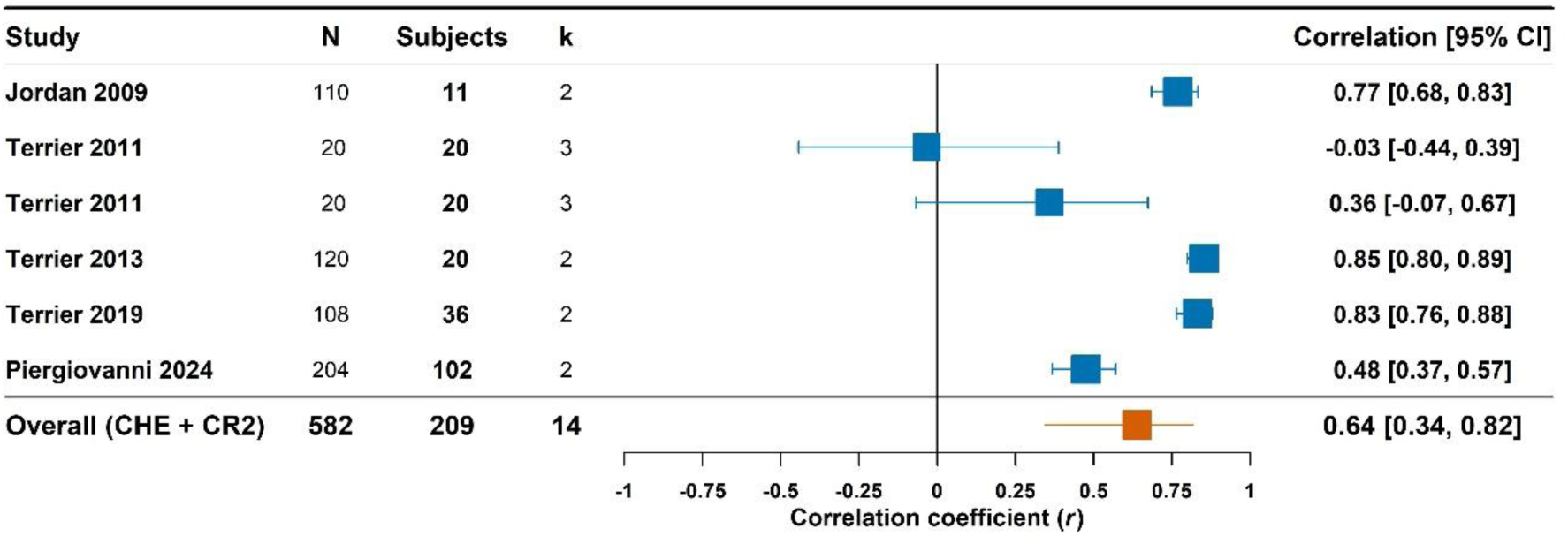
Meta-analysis of correlations between long-term DE and DFA scaling exponents. Forest plot showing study-level correlations with 95% confidence intervals and pooled estimate using correlated and hierarchical effects (CHE) model with robust variance estimation. N, total observations; k, number of correlation estimates per study; CI, confidence interval; DFA, detrended fluctuation analysis; DE, divergence exponent.

## Discussion

This systematic review synthesized evidence from 62 studies to critically examine the use of long-term divergence exponent (DE) computed by Rosenstein’s algorithm in human gait analysis, with particular focus on its reinterpretation as a measure of gait complexity and automaticity rather than stability.

### Summary of main findings

Our meta-analysis (Fig 6) is consistent with the theoretical reconceptualization of long-term DE as a complexity measure. Pooling six datasets, we observed moderate-to-substantial positive correlation (r = 0.64) between long-term DE and the DFA scaling exponent, which supports the hypothesis that these metrics capture a shared underlying construct, that is, the fractal-like correlation structure of stride-to-stride fluctuations. This synthesis must, however, be interpreted with caution: it rests on only six datasets, with substantial between-study heterogeneity (I² = 82). Two additional studies [33,48] showed parallel reductions in both long-term DE and scaling exponent during visual perturbation and external cueing, qualitatively supporting the meta-analytic correlation despite not reporting correlation statistics.

Beyond this formal meta-analytic synthesis, our review draws extensively on within-study contrasts between conditions or groups (Tables 1–3). This approach holds measurement system, signal processing, and algorithm parameters constant within each study, thereby attenuating the methodological heterogeneity that limits between-study quantitative synthesis.

The differential responsiveness of short-term and long-term DEs across experimental designs provides further support for their distinct physiological interpretations. Perturbation studies (Table 1) revealed a consistent dissociation: destabilizing interventions systematically increased short-term DE while decreasing long-term DE, confirming that these indices may reflect separable control processes underlying gait. This pattern directly contradicts the interpretation of both measures as stability metrics, since a truly destabilizing intervention should increase both if they assessed the same construct. Between-subject comparisons (Table 2) yielded more heterogeneous patterns, with clinical populations sometimes exhibiting paradoxical long-term DE responses (both increases and decreases relative to controls), further highlighting its dissociation from stability measures that should consistently elevate with pathology. Finally, external cueing studies (Table 3) confirmed this pattern with the largest effect sizes documented in the corpus: long-term DE reductions up to −86% accompanied by minimal short-term DE changes. This selective sensitivity to sensory cueing, combined with the parallel reductions observed in DFA scaling exponent [78], further support long-term DE as a marker of gait complexity rather than stability.

### Long-term divergence saturation as a signature of reduced gait complexity

#### Divergence in strange attractors

Rosenstein’s algorithm was originally developed to estimate the maximal Lyapunov exponent in deterministic chaotic systems. The method’s computational validity was established using well-characterized strange attractors, including the three-dimensional Lorenz system—a nonlinear dynamical system defined by three coupled differential equations that exhibits sensitive dependence on initial conditions [79]. When applied to such chaotic systems, the logarithmic divergence curve exhibits characteristic behavior: an initial linear region from which the maximal Lyapunov exponent is computed, followed by a plateau where further divergence ceases (see Appendix A in [40]). This saturation occurs because strange attractors, despite their complex internal dynamics, occupy a bounded region of phase space, constraining the maximum separation between initially neighboring trajectories.

#### Divergence in gait dynamics attractors

Dynamical systems models suggest that human gait may exhibit periodic limit cycle attractor dynamics with stochastic perturbations, rather than strange attractors characteristic of deterministic chaos [15,38,80], a characterization supported by both discrete stride interval analyses [81] and continuous signal phase space reconstruction [80]. This distinction justifies the preference for “divergence exponent” over “Lyapunov exponent” when describing gait dynamics [15]. The divergence curves obtained from gait attractors reflect departure from purely chaotic systems: they typically display an initial exponential divergence phase followed by gradual saturation over subsequent strides but often lack the abrupt plateau characteristic of purely deterministic chaotic systems with bounded strange attractors (S2 Figures).

#### Fractal stride patterns and phase space exploration

A gait attractor constructed using Takens theorem or an alternative approach represents the repertoire of available coordination patterns (i.e., the different kinematic combinations and their temporal organization) with phase-space exploration describing how trajectories navigate through this repertoire across consecutive gait cycles [82]. The temporal correlation structure of consecutive strides likely modulates how trajectories explore the phase space, thereby influencing divergence curve saturation and hence long-term DE values. Low-frequency variations distributed across dozens of consecutive strides may create drift through multiple regions of the attractor, delaying the saturation process and yielding higher long-term DE values.

This mechanism may explain the observed correlation between long-term DE and DFA scaling exponents (Fig. 6). In support of this hypothesis, Terrier and Reynard [53] manipulated stride interval correlation structure while preserving acceleration signal shape, demonstrating that long-term DE scaled systematically with temporal complexity while short-term DE remained unchanged. Divergence curves from correlated stride patterns exhibited delayed saturation compared to constant or random patterns (S2 Figures), providing evidence that fractal temporal structure prevents premature saturation of phase space exploration, thereby elevating long-term DE values. Consistent with this mechanism, three included studies [66,83,84] that applied stride-level time normalization by resampling each stride to a fixed number of data points (as in [53]) reported markedly suppressed long-term DE values (λL ≈ 0.00 in [83] or no significant condition effects [66,84]).

In silico evidence from mechanical walking-model simulations supports the same mechanism. A passive 2D dynamic walker [85] and a 3D walker with active lateral control [42] both produce divergence curves that plateau within two strides with long-term DE near zero, while their short-term DE remains responsive to perturbations and predicts fall risk in the models. The authors attribute this dissociation to the lack of neuromuscular variability characteristic of human gait, reinforcing the interpretation of long-term DE as a neurally-modulated complexity feature distinct from local mechanical stability and from fall risk per se.

#### Attention-dependent control mechanisms

The selective reduction of long-term DE during external pacing (Table 3) points to modifications in motor control strategy. Sensorimotor synchronization with rhythmic cues induces error correction processes that adjust each stride’s timing to match external targets [34]. This correction mode contrasts with the loosely coupled, endogenous control of self-paced walking [86]. The corrections around target pace values generate oscillations at short timescales while suppressing the low-frequency variations that characterize normal gait [87]. In fractal analysis terms (DFA), short-term oscillations manifest as anti-persistent temporal structure, whereas low-frequency variations represent the persistent, long-range correlations that define high-complexity gait dynamics. Experimental evidence shows that indeed metronome walking induces anti-persistent stride interval dynamics [78,88,89]. This transformation from persistent to anti-persistent temporal structure constrains phase space exploration (trajectories converge more rapidly within the attractor) thereby accelerating divergence curve saturation and reducing long-term DE values. This is evident in divergence curves published in [40,51,59] (see S2 Figures). The reductions in both long-term DE and DFA scaling exponents during cued walking [48,52] thus reflect a common underlying mechanism: the alteration of fractal correlation structure through attention-dependent error correction mechanisms.

#### Dual constraint effects

The minimal sensitivity of long-term DE to constant-speed treadmill constraint (Table 3) reflects parameter-specific alterations in temporal correlation structure. Evidence from DFA demonstrates that treadmill walking selectively induces anti-persistent dynamics in stride-speed fluctuations while preserving persistent correlations in both stride intervals and stride lengths [78]. This selective transformation arises because maintaining constant speed requires coordinated adjustments between stride time and stride length: participants modulate both parameters in tandem to prevent positional drift, thereby tightening speed control without imposing rigid constraints on either time series individually [34]. DEs computed from center-of-pressure trajectories or trunk acceleration signals likely capture a global fluctuation pattern, rather than the parameter-specific correlation structures revealed by DFA of individual time series. When only stride-speed correlations become anti-persistent while stride interval and stride length correlations remain persistent, this partial disruption produces moderate effects on the global attractor dynamics quantified by long-term DE. This interpretation is supported by the graded effects of constraint combinations observed in Table 3. Metronome synchronization during overground walking, which selectively modifies stride interval correlations while stride length and speed patterns remain persistent [88], produces moderate long-term DE reductions (−26% to −40% in [48], −13% to −30% in [52], Table 3). In contrast, superimposing metronome pacing on treadmill walking generates larger reductions (−66% to −86%, Table 3), reflecting combined modification of both stride interval and stride length correlation structures. This graded response demonstrates that dual constraints reorganize attractor dynamics more profoundly than either single constraint alone, consistent with the parameter-specific nature of correlation structure alterations captured by DFA.

#### Long-term divergence as an index of attractor complexity

Taken together, this evidence suggests that long-term DE reflects gait complexity rather than stability: it correlates with fractal scaling exponents, responds to manipulation of stride interval correlation structure while short-term DE remains unchanged, and exhibits graded sensitivity to constraint combinations that selectively modify correlations in different gait parameters. Walking-model simulations corroborate this picture in silico: removing the neuromuscular variability that humans produce suppresses long-term DE to near zero, while short-term DE remains responsive to perturbations. These findings support reinterpreting long-term DE as an “Attractor Complexity Index” that captures how temporal correlation structure shapes phase space exploration.

#### Long-term divergence and gait automaticity

This section examines long-term divergence exponent patterns across two complementary lines of evidence: between-subject comparisons in clinical populations, and within-subject perturbation studies that experimentally manipulate gait demands.

#### Between-subject studies

Among the studies identified in this review, only eight met methodological criteria for valid between-subject comparisons (Table 2), having employed sufficiently long walking bouts and time-normalized signals to ensure reliable long-term divergence exponent estimation. The remaining clinical comparison studies, while potentially informative, warrant cautious interpretation. Within these methodologically rigorous studies, long-term DE exhibited heterogeneous patterns across clinical populations. While short-term divergence consistently increased in patients relative to controls, long-term divergence showed both reductions and increases, which requires specific analysis separated by conditions and pathologies.

#### Pain

Two included studies examining pain-related conditions converged on a consistent pattern: patients with chronic lower-limb pain [74] and those with knee osteoarthritis [77], demonstrated reduced long-term divergence. An earlier longitudinal study of knee osteoarthritis patients ([90], N=16) awaiting replacement surgery provides additional support for pain-related complexity reductions. Using only 30 strides per analysis, the study nevertheless used global time normalization and generalized estimating equations that appropriately modeled speed-dependent effects. At a clinically relevant speed of 3 km/h, the affected limb demonstrated a dissociation between stability and complexity measures: short-term divergence exponents were unchanged between patients and controls, while long-term divergence exponents showed 19% reduction.

Pain may induce a shift toward more controlled, less automatic gait characterized by greater attentional involvement in motor planning [36]. Pain-related attentional focus on bodily sensations and movement execution likely constrains the natural stride-to-stride variability that characterizes automatic walking.

#### Peripheral Neuropathy

Only one study met methodological criteria for valid between-subject comparison (Table 2). However, the foundational work on this topic comes from the two earlier studies by Dingwell and colleagues [41,91] examining the same cohort of diabetic neuropathy patients (N=26). While these studies used sufficiently long walking bouts (10 minutes continuous), they did not use time-normalized signals. Nevertheless, they observed a consistent pattern: patients with diabetic neuropathy showed reduced long-term divergence exponents compared to controls (−26%). The pattern observed by Manor et al. [73] using 100 consecutive strides with time-normalized signals confirmed this finding with a 4% reduction in long-term DE. In contrast, the within-subject study that examined acute plantar desensitization in healthy adults showed an opposite effect: both short-term and long-term DEs increased ([67], Table 1). Such a result may point either to differential acute and habituation effects on gait dynamics or to heterogeneity in study design and measurement protocols.

Meta-analyses show that neuropathy patients walk more cautiously, with slower walking speed, shorter steps, and longer stance time [92,93]. Reduced plantar sensory information may make gait control more dependent on cognitive resources in diabetic patients compared to controls [93]. This higher cognitive demand is reflected in poorer dual-task performance [94]. Notably, these cautious gait adaptations are further amplified by cognitive impairment in diabetic patients, independent of neuropathy status [95]. The observed reduction in long-term DE may therefore reflect a compensatory shift toward a greater conscious control of gait characterized by a reduced reliance on automatic motor control.

#### Aging and fall risk

Three studies examining young versus older adults were excluded from this review due to methodological limitations. These studies generally showed trends toward increased short-term and long-term DEs in older populations [46], though the long-term DE increases rarely reached statistical significance [96,97]. Among the four methodologically valid age-comparison studies (Table 2), three converged on reduced long-term DE in older adults [33,52,72], with reductions amplified under conditions that increase attentional demands such as visual perturbations or metronome cueing. Only Bizovska et al. [75] reported variable results, though their older group (mean age 57.5 years) represented middle-aged rather than elderly adults.

Regarding fall risk, two studies comparing older fallers to non-fallers were excluded from the between-subject comparison. Toebes et al. [47] (N=134, mean age 62.4 years) used non-time-normalized angular velocity signals in composite high-dimensional state spaces, while Amirpourabasi et al. [39] (N=34, mean age 69.3 years) used non-time-normalized joint kinematics and trunk acceleration. Both studies found that short-term rather than long-term DE discriminated fall status. Nevertheless, the principal component analysis by Amirpourabasi et al. retained long-term DE along anteroposterior and vertical axes, which were independently identified as the most sensitive to complexity changes [51], suggesting that long-term DE may carry fall-related information that their analytical framework did not fully exploit. The only included fall-risk study, a prospective one-year follow-up by Bizovska et al. [76] (N=131), found no clear long-term DE difference between non-fallers and single fallers, but a more consistent reduction in frequent fallers (−7% to −18%).

Age-related decline in gait automaticity is well-documented [25], with older adults demonstrating increased prefrontal cortex activation during usual walking compared to standing [98], though patterns of cortex recruitment during cognitively demanding walking may differ from younger adults [99]. This shift from automatic to more controlled gait regulation has been attributed to age-related changes in neural networks, where diminished function in cerebellum and basal ganglia regions necessitates greater compensatory engagement of attention-related cortical networks, particularly the prefrontal cortex [100–102]. This compensatory reliance on executive resources appears even more pronounced in older adults at risk of falling: fallers exhibit greater prefrontal cortex activation than non-fallers during cognitively demanding tasks [103], and prospective evidence indicates that higher prefrontal activation during dual-task walking independently predicts incident falls [104]. The observed reductions in long-term DE in older adults, particularly under conditions that increase attentional demands such as visual perturbations, may therefore reflect this transition toward less automatic gait control.

#### Perturbation studies

Regarding perturbation studies, six studies were excluded from Table 1 due to methodological constraints. One study [66] used stride-level temporal normalization, which suppresses long-term divergence [53] and precludes any interpretation of long-term DE findings. Among the remaining excluded studies, two examined surface perturbations with contrasting results. Chang et al. [20] found significantly reduced long-term DE on a compliant foam surface compared to flat overground walking. In contrast, Emmerzaal et al. [105] reported increased long-term DE across six non-flat outdoor surfaces compared to flat-even walking. This discrepancy may partly reflect methodological limitations (short walking bouts, no time normalization). However, it may also reflect a genuine distinction between acute laboratory perturbations and familiar environmental terrain variation, which may not necessarily disrupt automatic control.

Among the eight retained treadmill studies (Table 1), a remarkably consistent pattern emerged: mechanical, visual, and vestibular perturbations increased short-term DE (+3% to +86%) while simultaneously decreasing long-term DE (−4% to −51%), with the notable exception of plantar desensitization which increased both measures. This is consistent with the hypothesis that individuals allocate greater attentional resources to gait control to manage the imposed challenges.

Neuroimaging studies provide direct support for this interpretation. Unpredictable slip perturbations during treadmill walking induce significant increases in prefrontal cortex activation as measured by functional near-infrared spectroscopy [106]. Visual perturbations modulate electrocortical dynamics during treadmill walking, revealing increased cortical processing demands under perturbed sensory conditions [107]. Last but not least, a systematic review of mobile brain imaging studies further confirmed that unpredicted mechanical perturbations elicit prominent cortical activation in frontal regions [108].

#### Long-term divergence as an index of gait automaticity

Across clinical populations, experimental perturbations, and neuroimaging evidence, a coherent pattern emerges: conditions that increase cortical involvement in gait control consistently reduce long-term DE, while short-term DE tracks dynamic stability independently. Pain, neuropathy, and aging each shift gait toward greater executive oversight, and this shift is mirrored by reduced long-term divergence. Perturbation studies confirm this relationship experimentally, with neuroimaging demonstrating that the same manipulations that decrease long-term DE simultaneously increase prefrontal and frontal cortical activation. These converging lines of evidence support the interpretation that long-term DE captures the degree of gait automaticity, that is the extent to which locomotor control operates through automatic, subcortically-mediated processes rather than attention-dependent executive control.

### Methodological considerations

#### Measurement methods

The included studies employed diverse measurement approaches (Fig 3). Despite this methodological heterogeneity, examination of response patterns across experimental manipulations reveals consistent findings, suggesting that long-term DE captures aspects of gait control that transcend specific measurement modalities. The critical prerequisite is likely that the measured signal captures the rhythmic pattern of locomotor cycles with their inherent stride-to-stride variations. Whether these dynamics are sampled through trunk accelerations, joint angle trajectories, or center-of-pressure displacements, the resulting measures converge on the same underlying phenomenon: the temporal organization of consecutive gait cycles.

Among the measurement approaches employed, trunk-level assessment offers particular advantages for capturing global gait coordination patterns. A majority of included studies (34/62, 55%) prioritized trunk measurements. Dingwell and Kang [80] emphasized that trunk movements reflect the integrated outcome of bilateral limb coordination and postural adjustments required to maintain upright progression. For assessments targeting the temporal organization of overall locomotor patterns, trunk-level measurements therefore represent the preferred approach, offering an integrated readout of stride-to-stride control dynamics. Similarly, center-of-pressure trajectories derived from instrumented treadmills (used in 3 included studies) are another global measure of locomotor control, despite being recorded at the foot-ground interface. The center of pressure reflects the net ground reaction force application point and integrates the postural and propulsive adjustments required to regulate balance and forward progression.

#### Measurement duration

Assessment of long-term DE as a complexity measure imposes specific data duration requirements that differ from stability assessments. The basis for these requirements arises from the multiscale nature of gait complexity. Low-frequency variations, which are typical of persistent temporal correlations, manifest only across extended observation windows. Truncated recordings constrain the observable correlation timescales, filtering out the very low-frequency components. This explains why studies employing shorter recordings (fewer than 50 strides) may capture local stability properties adequately but potentially mischaracterize attractor complexity.

Methodological investigations within the included studies provide empirical guidance for minimum data requirements. Bruijn et al. [109] showed that divergence exponent precision improves with recording length, identifying approximately 150 consecutive strides as necessary for adequate measurement precision. Reynard and Terrier [63] confirmed these requirements, showing that 70-stride estimates yielded substantially higher reliability than 35-stride estimates for long-term DE. Riva et al. [110] observed that long-term divergence measures required a high stride count (110+ strides for 50% reliability threshold) compared to other stability metrics.

A 2020 systematic review on DFA methodology [111] established consensus guidelines recommending at least 500-600 consecutive strides for reliable estimation of scaling exponents, acknowledging that shorter recordings may inadequately capture the long-range correlation structure defining fractal gait dynamics. The more modest stride requirements for long-term DE (150-200 strides) compared to DFA’s 500-600 stride recommendation suggest practical advantages for clinical gait assessment, requiring shorter recording durations while still capturing essential features of temporal correlation structure through phase space reconstruction [51]. Future investigations employing long-term divergence exponents as complexity measures should prioritize recording durations of at least 150-200 consecutive strides to ensure adequate precision for detecting meaningful differences while maintaining practical feasibility for clinical populations.

#### Data length effects and time normalization

Beyond precision considerations, measurement duration influences DE magnitudes. Bruijn et al. [109] demonstrated that both short-term and long-term DEs increase with data series length. As recording duration extends, the phase space trajectory explores increasingly distant regions of the attractor, and allows the algorithm to identify nearest neighbors separated by larger distances. These more distant neighbor pairs exhibit different divergence characteristics than the tightly clustered neighbors identified in shorter recordings. Reynard and Terrier confirmed this effect in a larger sample (N=95) [63]. Consequently, standardizing recording duration across experimental conditions and participant groups constitutes a critical methodological requirement, as divergence exponent differences attributable to unequal measurement lengths may confound or obscure true physiological differences between populations or conditions.

### Study quality and risk of bias

#### Analytical rigor and methodological transparency

Quality assessment revealed good methodological transparency across the included studies (S3 Table), with comprehensive reporting of experimental protocols, measurement systems, signal processing procedures, and algorithm parameters necessary for study replication. This result may indicate that the field has achieved methodological maturity, with recent studies providing sufficient detail to enable independent replication. However, it is important to acknowledge that these favorable quality metrics partly reflect our inclusion criteria, which required studies to provide adequate methodological information to confirm use of Rosenstein’s algorithm for long-term DE computation.

#### Outcome reporting: descriptive statistics

While analytical procedures were generally well-documented, the reporting of study outcomes showed considerable variability. A substantial proportion of studies, particularly those published before 2013, presented results exclusively through figures without accompanying numerical tables containing means and measures of variability. This reporting approach necessitated manual extraction of approximate values using specialized digitization software to enable quantitative synthesis and calculation of percentage changes between conditions or groups (as presented in Tables 2 and 3). The reliance on visual estimation from figures introduces measurement error and limits the precision of secondary analyses. Moreover, some studies reported only qualitative descriptions of differences or provided statistical test results without the underlying descriptive statistics, making meta-analytical synthesis unfeasible. A clear temporal improvement emerged after 2013, with recent studies increasingly providing complete descriptive statistics for all experimental conditions.

#### Outcome reporting: effect sizes and clinical significance

Beyond descriptive statistics, a more persistent limitation concerned the reporting and interpretation of effect sizes. Many studies across all time periods relied primarily on p-values and significance testing to characterize findings, with only 40% achieving full scores through comprehensive effect size reporting. Many studies reported results dichotomously as “significant” or “non-significant” without quantifying the magnitude of observed differences or discussing their potential clinical relevance. This approach limits the ability to distinguish between statistically significant but trivially small effects and meaningfully large effects that may warrant clinical attention. The absence of standardized effect size measures (such as Cohen’s d, confidence intervals, or percentage differences with precision estimates) also constrains meta-analytic synthesis and prevents assessment of whether effects are consistent or variable across studies and populations. While some recent studies have begun incorporating effect size reporting and discussing practical significance, this remains an underdeveloped aspect of reporting standards in the field.

#### Sample size considerations and recruitment practices

Domain 3 (sample size adequacy) achieved an average score of 1.7 out of 3 points, reflecting the modest sample sizes typical of exploratory biomechanical research (S3 Table). The median sample size across studies was 23 participants, with most studies enrolling fewer than 50 individuals. Given the exploratory and methodological nature of much of this research, these sample sizes are not inappropriate, particularly considering that most studies used within-subjects designs and repeated measures that enhance statistical power relative to between-subjects comparisons. However, only a minority of studies (<10%) provided explicit sample size justifications or reported formal power calculations, making it difficult to assess whether samples were adequate for detecting meaningful effects. A more substantive concern relates to recruitment practices and sample representativeness. Although rarely discussed explicitly in the reviewed articles, convenience sampling appeared widespread, with many studies likely recruiting from institutional populations such as university students or laboratory staff members. This practice may introduce systematic biases, as such samples tend to overrepresent younger, healthier, more educated, and potentially more physically active individuals compared to the general population. For studies comparing healthy controls to clinical populations, the use of convenience samples may exaggerate group differences if controls represent an unusually homogeneous and high-functioning subset of the healthy population. While these recruitment practices are understandable during the methodological development phase of a field, the transition toward clinical application requires more systematic attention to sample representativeness, demographic diversity, and explicit justification of sample sizes relative to the precision needed for clinical decision-making.

### Limitations of the systematic review

#### Study identification and selection limitations

This systematic review has several limitations that should be acknowledged. First, the favorable quality assessment results partly reflect selection bias inherent to our inclusion criteria. Studies were required to provide sufficient methodological detail, resulting in exclusion of less transparently reported research during full-text screening. Second, while our database searches applied no language restrictions to ensure comprehensive literature retrieval, our protocol specified that full-text screening would be limited to English and French publications based on the linguistic competencies of our review team. The language restriction remained theoretical rather than operational, as no relevant non-English/French studies required exclusion. Nevertheless, we acknowledge that future updates to this review would benefit from broader linguistic inclusion with appropriate translation resources to ensure maximum comprehensiveness. Third, our focus on peer-reviewed journal articles resulted in exclusion of grey literature including doctoral dissertations, conference abstracts, and technical reports, which may contain relevant methodological developments or findings not yet published in traditional venues. Fourth, publication bias may affect our synthesis, as studies reporting null findings or unsuccessful applications of the method may be less likely to appear in peer-reviewed literature. This bias may have been particularly pronounced because most studies interpreted long-term DE as a stability measure. Paradoxical patterns, such as long-term DE reductions during destabilizing perturbations (Table 1), may therefore have been regarded as methodological failures rather than meaningful results, leading to selective non-reporting of theoretically inconsistent outcomes.

#### Quality-assessment limitations

Our three-domain, 15-point quality-assessment framework was developed as a fit-for-purpose instrument for a methodological review of an analytical technique, in line with the practice of comparable systematic reviews in this subfield. It carries three principal limitations. First, scores are not directly comparable to those produced by standardized risk-of-bias instruments and should be interpreted as relative indicators within the present corpus rather than as absolute quality estimates. Second, our domains emphasize reporting completeness and analytical rigor over clinical effect estimation. Third, the psychometric properties of the framework (inter-rater reliability, construct validity) have not been formally evaluated. The complete scoring grid is provided in S3 Table to allow readers to assess the rationale for each criterion.

#### Synthesis limitations

The scope of quantitative synthesis was constrained by several factors. Despite identifying 62 relevant studies, comprehensive meta-analysis across all research questions was not feasible due to heterogeneous outcome reporting, particularly in earlier studies that presented results exclusively through figures without numerical tables. Only five studies provided the necessary correlation data for our meta-analysis examining associations between long-term DE and DFA scaling exponents, limiting the precision of this key finding. Similarly, only eight studies met methodological criteria for valid inclusion in between-subject clinical comparisons (Table 2), as most clinical studies used insufficiently long recording durations or lacked time normalization procedures necessary for reliable complexity assessment. The diverse measurement contexts—including different walking conditions, signal types, and body locations, while potentially strengthening the generalizability of findings, precluded direct quantitative comparison of effect magnitudes across studies. These synthesis limitations reflect the exploratory developmental stage of the field rather than fundamental obstacles to future evidence accumulation, as recent improvements in reporting standards suggest that subsequent systematic reviews will be better positioned for comprehensive meta-analytic synthesis.

### Future directions and clinical translation

#### In silico validation

A natural next step is to assess the ACI hypothesis through dedicated modelling. The bipedal-robot modelling literature has extensively analyzed simple passive walkers within the framework of deterministic chaos [112]. Closer to the framework adopted here, passive dynamic walker simulations with stochastic perturbations reproduce the short-term DE response to local instability but yield near-zero long-term DE, in contrast with the sustained divergence observed across multiple strides in empirical human gait [42,43,85]. A direct test of the ACI hypothesis would therefore consist in driving a passive-walker model with biologically realistic, fractal stride-interval fluctuations. Demonstrating that such fractal driving suffices to recover the empirical long-term DE pattern would provide a mechanistic *in silico* counterpart to the empirical and quasi-experimental evidence reviewed here, and would close the conceptual loop linking gait complexity, automaticity, and long-term DE.

#### Methodological advances

Several methodological priorities emerge from this review to advance the field toward clinical application. First, standardization of measurement protocols should prioritize recording durations of at least 150-200 consecutive strides for complexity assessment. Second, theoretical development remains incomplete. Establishment of clinically meaningful change thresholds for long-term DE is essential—what magnitude of reduction indicates functionally significant loss of gait automaticity? Development of age-stratified and population-specific normative reference values would enable interpretation of individual measurements relative to expected ranges. Third, validation of the complexity interpretation requires experimental studies that directly manipulate automaticity (e.g., dual-task paradigms, attention-demanding environments) while simultaneously measuring long-term DE and neuroimaging markers of cortical engagement. Finally, longitudinal research tracking long-term DE changes across disease progression, rehabilitation, or aging trajectories would clarify whether complexity measures predict clinically relevant outcomes such as fall risk, functional decline, or treatment response. Multi-site replication studies ensuring generalizability across diverse populations and measurement contexts remain particularly valuable given the current concentration of research within specialized laboratories.

#### Clinical translation and practical applications

Long-term DE holds promise as a complementary clinical measure that captures aspects of motor control not reflected in traditional gait metrics. Its potential clinical utility spans several domains. For diagnostic applications, reduced long-term DE may identify early-stage neurological changes characterized by diminished gait automaticity before gross functional impairments become apparent. In rehabilitation contexts, long-term DE could monitor recovery of automatic motor control following neurological injury or track treatment effects where restoration of natural gait patterns represents a therapeutic goal. For fall risk assessment, integration of complexity measures with conventional stability metrics may improve risk stratification. The method offers practical advantages over alternative complexity measures such as DFA: the more modest recording requirements (150-200 versus 500-600 strides) enhance feasibility for clinical populations with limited walking tolerance. Long-term DE should complement rather than replace existing clinical gait assessment tools, contributing one element within comprehensive evaluation protocols. With these foundations established, wearable sensor technologies could enable free-living complexity monitoring, extending assessment beyond laboratory settings to capture gait automaticity in natural environments where attentional demands and environmental challenges reflect real-world function.

## Conclusion

This systematic review provides converging evidence that long-term DE measures gait complexity and automaticity rather than stability. The substantial correlation with DFA scaling exponents (r = 0.64), combined with systematic dissociation from short-term stability measures during perturbations and cueing, supports reinterpreting this metric as the Attractor Complexity Index—a marker of the temporal organization and attentional demands of locomotor control.

This paradigm shift has practical implications. Rather than duplicating information from short-term stability measures, long-term divergence may uniquely capture motor-cognitive aspects of gait control relevant to fall risk, disease progression, and rehabilitation effectiveness. Establishing this measure as a complementary biomarker for gait automaticity would enhance multidimensional assessment of locomotor function in clinical populations.

## Supporting information

Supplemental file 1 (outcome tables)

Supplemental file 2 (divergence curves)

Supplemental file 3 (Quality assessment)

## Acknowledgments

We thank Marco Pedrotti and Florence Waelchli for their assistance with administrative tasks.

## Supporting Information

**S1 Text. Systematic review database.** Complete data extraction tables with study characteristics (Table A: research types; Table B: study characteristics; Table C: methodological specifications), search strategy, and detailed information for all 62 included studies. (PDF)

**S2 Figures. Divergence curve figures.** Compiled divergence curves from 30 studies with detailed figure descriptions showing methodological illustrations and comparative results. (PDF)

**S3 Table. Quality assessment results.** Complete quality assessment scores for all 62 studies using the three-domain framework (maximum 15 points). (XLSX)

