## Supplemental file 1 (outcome tables) for "From Stability to Complexity: A Systematic Review of Long-term Divergence Exponents in Nonlinear Gait Analysis"

¶ Equal contribution

### Web of Science: Search strategy

Query date:

February 2024 (updated search, January 2026)

**Note:** Studies identified in the original search (February 2024) are numbered 1–56. Studies identified in the updated search (January 2026) are numbered S1–S6 and cited from a separate bibliography at the end of this document ("References — Update screening").

#### Search chain:

**TS=(walking OR gait OR ambul*) AND TS=(Lyapunov OR "local dynamic stability" OR "divergence exponent*") AND TS=( child* OR adult* OR people OR subject* OR person* OR participant*)**

TS: Searches for topic terms in the following fields within a record. Title, Abstract, Author Keywords, Keywords Plus® (KeyWords Plus® are index terms automatically generated from the titles of cited articles. KeyWords Plus terms must appear more than once in the bibliography and are ordered from multi-word phrases to single terms. KeyWords Plus augments traditional keyword or title retrieval)

#### Databases:

#### Web of Science Core Collection, MEDLINE.

#### Publication years:

2000-2024 (updated search 2024-2026)

COPYRIGHT 2026 THE AUTHORS. THIS FILE IS LICENSED UNDER A CREATIVE COMMONS ATTRIBUTION 4.0 INTERNATIONAL (CC BY 4.0) LICENSE.

### Results tables

#### A. Categorization criteria (research type)

| Category | Definition |
|---|---|
| Diagnostic | Identify gait abnormalities or classify gait patterns |
| Interventional | Assess effects of treatments, therapies, or interventions on gait |
| Comparative | Compare gait patterns between different populations (between subjects, BS) or conditions (within subjects, WS) or both (BS/WS) |
| Predictive | Use gait analysis to predict outcomes or classify individuals |
| Method Development | Develop or improve gait analysis techniques or technologies |
| Validation | Validate new gait analysis methods, technologies, or algorithms |
| Reliability | Assess reliability and repeatability of gait analysis methods |
| Longitudinal | Track changes in gait patterns over time |
| Clinical Application | Evaluate use of gait analysis in clinical decision-making |

#### B. Authors, sample size, and research type

| ID | Authors | Year | Sample size | Sample composition | Research type |
|---|---|---|---|---|---|
| 1 | Dingwell et al. [1] | 2000 | 26 | 14 diabetic patients (peripheral neuropathy)<br>12 matched healthy controls | Method dev.<br>Comparative BS |
| 2 | Dingwell & Cusumano [2] | 2000 | 36 | 10 young healthy adults<br>12 matched healthy controls<br>14 diabetic patients | Method dev.<br>Comparative BS/WS<br>Validation |
| 3 | Dingwell & Cusumano [3] | 2001 | 10 | 10 young healthy adults | Method dev.<br>Comparative WS<br>Validation |
| 4 | Yoshino et al. [4] | 2004 | 12 | 12 young healthy adults | Comparative BS/WS<br>Predictive |
| 5 | Dingwell & Marin [5] | 2006 | 12 | 12 young healthy adults, 1 discarded (final N=11) | Comparative WS<br>Validation |
| 6 | Kang & Dingwell [6] | 2006 | 20 | 20 healthy adults aged 18–73 | Reliability<br>Method dev. |
| 7 | Dingwell & Kang [7] | 2007 | 10 | 10 young healthy adults (same as in 3) | Comparative WS<br>Method dev.<br>Validation |
| 8 | Kang & Dingwell [8] | 2008 | 36 | 18 healthy older adults,<br>18 (17) height-, weight-, and gender-matched young adults. (one subject discarded) | Comparative BS/WS |
| 9 | Manor et al. [9] | 2008 | 24 | 12 individuals with all-cause peripheral neuropathy<br>12 age-, body mass-, and height matched healthy controls. | Comparative BS/WS<br>Clinical application |

COPYRIGHT 2026 THE AUTHORS. THIS FILE IS LICENSED UNDER A CREATIVE COMMONS ATTRIBUTION 4.0 INTERNATIONAL (CC BY 4.0) LICENSE.

| ID | Authors | Year | Sample size | Sample composition | Research type |
|---|---|---|---|---|---|
| 10 * | Jordan et al. [10] | 2009 | 11 | Enrolled: 12 female adults, recreational runners. One subject discarded | Comparative WS |
| 11 | Bruijn et al. [11] | 2009 | 9 | 9 healthy young adults | Reliability Validation |
| 12 | Bruijn et al. [12] | 2009 | 15 | 15 healthy young adults | Comparative WS Method dev. |
| 13 | Kang & Dingwell [13] | 2009 | 35 | 18 healthy older adults 17 matched young adults | Comparative BS Method dev. |
| 14 | Kang & Dingwell [14] | 2009 | 35 | 18 healthy older adults 17 matched young adults | Comparative BS/WS Method dev. |
| 15 | Manor et al. [15] | 2009 | 13 | 13 healthy young adults | Comparative WS Method dev. |
| 16 | Nessler et al. [16] | 2009 | 14 | 14 healthy young adults | Comparative WS |
| 17 | Son et al. [17] | 2009 | 40 | 40 young healthy subjects | Method dev. Validation |
| 18 | Bruijn et al. [18] | 2010 | 11 | 11 healthy young male subjects | Comparative WS Method dev. |
| 19 | Bruijn et al. [19] | 2010 | 9 | 9 healthy males | Validation Method dev. |
| 20 * | Chang et al. [20] | 2010 | 14 | 14 able-bodied adult participants | Comparative WS |
| 21 | Yakhdani et al. [21] | 2010 | 26 | 14 older patients with knee osteoarthritis in waiting list for replacement surgery 12 healthy matched controls | Interventional Longitudinal Comparative BS/WS Clinical application |
| 22 | McAndrew et al. [22] | 2011 | 12 | 12 healthy young adults | Comparative WS Validation |
| 23 * | Nessler et al. [23] | 2011 | 15 | 15 recreationally active adults | Comparative WS |
| 24 | Son et al. [24] | 2011 | 40 | 40 healthy young adults (20 males, 20 females) | Method dev. Validation Comparative BS |
| 25 * | Terrier and Dériaz [25] | 2011 | 20 | 20 healthy male adults | Comparative WS |
| 26 | Van Schooten et al. [26] | 2011 | 12 | 12 healthy young adults | Comparative WS Validation |
| 27 | McAndrew et al. [27] | 2012 | 14 | 14 healthy young adults | Comparative WS |
| 28 | Cignetti et al. [28] | 2012 | 14 | 7 healthy older adults 7 healthy matched younger adults | Validation Method dev. Comparative BS |
| 29 * | Sejdić et al. [29] | 2012 | 15 | 15 healthy young adults | Comparative WS |
| 30 | Sinitksi et al. [30] | 2012 | 11 | 11 healthy young adults | Comparative WS |
| 31 | Toebes et al. [31] | 2012 | 134 | 44 elderly fallers 90 elderly non-fallers | Predictive Comparative BS Method dev. |
| 32 | Liu & Lockhart [32] | 2013 | 25 | 25 young adults | Comparative WS Validation |
| 33* | Nessler et al. [33] | 2013 | 26 | 26 young adults | Comparative WS Method dev. |
| 34 | Sejdić et al. [34] | 2013 | 17 Exp.1 15 Exp.2 | 17 young adults in Music experiment 15 young adults in TV experiment | Comparative WS |
| 35 * | Terrier & Dériaz [35] | 2013 | 20 | 20 healthy adults | Comparative WS Validation |
| 36 | Reynard & Terrier [36] | 2014 | 95 | 100 enrolled: Ten male and 10 female healthy adults for each decade, between 20 and 69 years old | Reliability Validation |
| 37 | Riva et al. [37] | 2014 | 10 | 10 healthy young adults | Reliability Validation |
| 38 | Riva et al. [38] | 2014 | 51 | 51 healthy young adults | Method dev. Validation Reliability |

COPYRIGHT 2026 THE AUTHORS. THIS FILE IS LICENSED UNDER A CREATIVE COMMONS ATTRIBUTION 4.0 INTERNATIONAL (CC BY 4.0) LICENSE.

| ID | Authors | Year | Sample size | Sample composition | Research type |
|---|---|---|---|---|---|
| 39 | Yang & Pai [39] | 2014 | 187 | 187 community-dwelling older adults | Predictive |
| 40 | Beaudette et al. [40] | 2015 | 12 | 12 healthy young adults | Comparative WS |
| 41 * | Franz et al. [41] | 2015 | 23 | 11 healthy older adults<br>12 healthy younger adults | Comparative BS/WS<br>Method dev. |
| 42 | Rábago et al. [42] | 2015 | 20 | 20 healthy young adults | Reliability<br>Comparative WS<br>Validation |
| 43 | Reynard & Terrier [43] | 2015 | 100 | Ten male and 10 female healthy adults for each decade, between 20 and 69 years old, same sample as 36. | Comparative WS |
| 44 | Rantalainen et al. [44] | 2016 | 39 | 39 healthy young adults | Validation<br>Method dev. |
| 45 | Terrier et al. [45] | 2017 | 147 | 66 middle-aged patients with chronic pain of lower limbs after an orthopedic trauma<br>81 healthy controls | Validation<br>Reliability<br>Comparative BS<br>Clinical application |
| 46 | Bizovska et al. [46] | 2018 | 26 | 13 healthy young females<br>13 healthy older females | Comparative BS/WS<br>Validation |
| 47 | Qiao et al. [47] | 2018 | 33 | 11 healthy young adults<br>11 older non-faller adults<br>11 older faller adults | Comparative BS/WS<br>Predictive |
| 48 * | Terrier & Reynard [48] | 2018 | 69 | 69 middle aged subjects from a sample of 100 subjects, reanalysis of study #36. | Validation<br>Method dev. |
| 49 | Bizovska et al. [49] | 2018 | 131 | 81 older adults, non-fallers<br>35 older adults, fallers<br>15 older adults, frequent fallers | Predictive<br>Clinical application<br>Comparative BS |
| 50 | Matinazad et al. [50] | 2018 | 14 | 7 females with knee osteoarthritis<br>7 healthy female controls | Comparative BS/WS<br>Method dev.<br>Reliability |
| 51 * | Terrier [51] | 2019 | 36 | 36 healthy adults | Method dev.<br>Validation<br>Comparative WS<br>Predictive |
| 52 | Wodarski et al. [52] | 2020 | 40 | 40 young healthy adults | Comparative WS<br>Method dev. |
| 53 | Chinimilli et al. [53] | 2020 | 11 | 11 healthy young adults | Method dev.<br>Comparative WS |
| 54 * | Sedighi et al. [54] | 2020 | 20 | 20 healthy young adults | Comparative WS<br>Method dev. |
| 55 | Sarvestan et al. [55] | 2022 | 29 | 29 healthy young adults | Comparative WS |
| 56 | Ulman et al. [56] | 2022 | 60 | 30 inexperienced military cadets<br>30 experienced military cadets | Comparative BS/WS |
| S1 | Yamagata et al. [S1] | 2024 | 10 | 10 healthy young adult males | Comparative WS<br>Validation |
| S2* | Piergiovanni & Terrier [S2] | 2024 | 60 | 60 healthy older adults | Comparative WS<br>Reliability<br>Validation |
| S3 | Bhat & Kaufman [S3] | 2024 | 33 | 33 healthy older adults | Method dev. |
| S4* | Piergiovanni & Terrier [S4] | 2024 | 102 | 60 healthy older adults (same as S2)<br>42 healthy younger controls | Comparative WS/BS<br>Validation |
| S5 | Emmerzaal et al. [S5] | 2025 | 30 | 30 healthy adults | Comparative WS |
| S6 | Amirpourabasi et al. [S6] | 2026 | 34 | 17 healthy older women, non-fallers<br>17 healthy older women, fallers | Comparative BS<br>Predictive |

#### C. Methodological outcomes

COPYRIGHT 2026 THE AUTHORS. THIS FILE IS LICENSED UNDER A CREATIVE COMMONS ATTRIBUTION 4.0 INTERNATIONAL (CC BY 4.0) LICENSE.

| ID | Measurement system | Signal type | Duration | Walking condition | Filtering method | Resampling method | Attractor reconstruction | Stride range |
|---|---|---|---|---|---|---|---|---|
| 1 | Electrogoniometers for hip, knee, and ankle joint angles. Tri-axial accelerometer for trunk (sternum) accelerations (66.7 Hz) | Sagittal plane angles of hip, knee, and ankle. joints Tri-axial accelerations of the trunk | 10 min walking (5x2 min, ~100 str.) | OG, 200m indoor track | None | Not mentioned | T: variable (AMI) dE: 5 (GFNN) | 4-10 |
| 2 | Electrogoniometers for hip, knee, and ankle joint angles. Tri-axial accelerometer for trunk (sternum) accelerations (66.7 Hz) | Sagittal plane angles of hip, knee, and ankle. joints Tri-axial accelerations of the trunk | 10 min walking (5x2 min, ~100 str.) | OG and TW | None | Not mentioned | T: variable (AMI) dE: 5 (GFNN) | 4-10 |
| 3 | Electrogoniometers for hip, knee, and ankle joint angles. Tri-axial accelerometer for trunk (sternum) accelerations (66.7 Hz) | Sagittal plane angles of hip, knee, and ankle. joints Tri-axial accelerations of the trunk | 10 min walking (5x2 min, ~100 str.) | OG and TW | None | Not mentioned | T: variable (AMI) dE: 5 (GFNN) | 0-1, 4-10 |
| 4 | Triaxial accelerometer (low-back), 1000 Hz | Tri-axial accelerations of the trunk | 12x15min walking, 3 hour continuous walking. | OG | 1000Hz down sampled to 100Hz | Not mentioned | T: variable (autocorrelation ,time to 1/e) dE : 5 (GFNN) | 0.5-4 |
| 5 | Six-camera (infrared) motion capture system (60 Hz) tracking a single marker on thoracic spine | First difference of trunk position data (~velocity) | 15 recorded trials of 3min. (3 trials × 5 speeds). ~150 strides per trial | TW | None | Not mentioned (? Time normalization limited to MeanSD computation ?) | T: variable (AMI) dE: 5 (GFNN) | 0-1, 4-10 |
| 6 | 3D motion capture system (60 Hz) with six markers placed on trunk landmarks | 3D linear trunk motions (position, velocity) and 3D angular trunk motions (position, velocity) | 3 x 5 min walking, then separated in 1-5 min. lengths | TW | None | Not mentioned | 12-dimensional state space combining trunk linear and angular positions and velocities, demeaned and normalized to unit variance | 0-1, 4-10 |
| 7 | Electrogoniometers for hip, knee, and ankle joint angles. Tri-axial accelerometer for trunk (sternum) accelerations (66.7 Hz) | Sagittal plane angles of hip, knee, and ankle. joints Tri-axial accelerations of the trunk | 10 min. walking. No further details are given. Very likely like in 3. | OG and TW | None | Not mentioned | T: variable (AMI) dE: 5 (GFNN) | 0-1, 4-10 |
| 8 | 3D motion capture system with six markers placed on trunk landmarks. Markers on both feet to track heel-strike events | 3D linear trunk motions (velocity, acceleration) and 3D angular trunk motions (velocity, acceleration) | 10 (2x5) 5 min walking trials | TW | 10 Hz low pass, zero-lag Butterworth filter. | Not mentioned | 12-dimensional state space combining trunk linear and angular velocities and accelerations | 0-1, 4-10 |
| 9 | 2D motion capture system (60 Hz) | Sagittal plane joint angles for hip, knee and ankle | 3x3 min walking, 100 consecutive strides | TW | None | Time-series were time-normalized to 4905 data points | T: variable (AMI) dE: 5 (GFNN) | 0-1, 4-10 |
| 10 * | 3D motion analysis system (125 Hz) | Vertical oscillations of the head and ankle markers | Trials of 5 min duration. 10 walking trials at different | TW | 6 Hz low pass zero-lag second-order | Not mentioned | T: fixed, for head and ankle separately (AMI) dE: 5 (GFNN) | 4-10 |

COPYRIGHT 2026 THE AUTHORS. THIS FILE IS LICENSED UNDER A CREATIVE COMMONS ATTRIBUTION 4.0 INTERNATIONAL (CC BY 4.0) LICENSE.

| ID | Measurement system | Signal type | Duration | Walking condition | Filtering method | Resampling method | Attractor reconstruction | Stride range |
|---|---|---|---|---|---|---|---|---|
| | | | speeds. All strides included in the analysis | | Butterworth filter. | | | |
| 11 | Active 3D movement registration system (50 Hz) tracking a single marker on thoracic spine | First derivatives of the 3D trunk positions. | Samples of 30 to 300 strides were extracted from 2x20-min walking trials. | TW | None | Time-series were time-normalized to, on average, 100 samples per stride. | T: fixed to10, no method mentioned dE: 5 (GFNN) | 0-0.5, 4-10 |
| 12 | Active 3D movement registration system (50 Hz) tracking a single marker on thoracic spine | First derivatives of the 3D trunk positions. | 6 x 2 min walking. 50 strides | TW | None | Time series time-normalized via shape-preserving spline; 50 strides resampled to 5000 samples | T: fixed to10, after AMI analysis. dE: 5 (GFNN) | 0-0.5, 4-10 |
| 13 | 8-camera 3D analysis system with marker clusters on trunk, pelvis, left thigh, left shank and both feet. | Linear and angular velocities of trunk, pelvis, thigh, shank, and foot. | 2 x 5 min walking. Number of strides not mentioned. | TW | 10 Hz low-pass filtered, zero-lag Butterworth filter. | Not mentioned | 12-dimensional state space combining linear and angular velocities and their time delayed copies. T: variable (AMI) | 0-1, 4-10 (inferred from reference to older studies) |
| 14 | Surface electromyography (EMG, 1080Hz), using bipolar electrodes from 4 muscles of the left leg. Use kinematic data from a previous study. | EMG voltage with passband filtering and amplitude normalization. Measured for muscles of the leg. | 10 (2x5) 5 min walking trials | TW | 10 Hz low-pass filtered (Hamming averaging window). Down sampled to 60Hz | Not mentioned | 8-dimensional state space combining normalized EMG signals and their first derivatives. | 0-1, 4-10 |
| 15 | Single camera motion analysis system (60 Hz). | Sagittal plane hip, knee, and ankle angles. | 6 (3 x 2) min trials. Only last minute recorded. 45 strides included. | TW | Not mentioned | 45 strides time-normalized to 1930 data points with linear interpolation. | T: variable (maximum attractor volume ?) dE: 5 (GFNN) | 0-1, 4-10 |
| 16 | 6 camera optical motion capture system (120Hz), with markers over the toe, heel, lateral maleolus, lateral knee, and lateral thigh. | Ankle and knee angles, ankle 3D positions, knee vertical position. | 6 min walking (3x2 min). ~100 strides per trial. | TW | 4th order Butterworth, cutoff = 100 Hz | Normalized in time with respect to the overall mean cadence. | T: variable (AMI) dE: 6 (GFNN) | 0-1, 4-10 |
| 17 | 8 camera 3D motion capture system with 24 markers. (60Hz) | Flexion-extension angles of hip, knee and ankle. (bilateral) | 90 s walking | TW | Not mentioned | Not mentioned | T: variable (AMI) dE: 5 (GFNN) | 4-10 |
| 18 | 6 camera 3D optoelectronic system, with markers on thoracic spine and heels (50Hz). | 3D Linear and angular velocities of the trunk. | 12 2x(2x3) 5 min walking trials. 140 strides | TW | Not mentioned | Time series normalized to 140 strides using spline interpolation | 12-dimensional state space combining linear and angular velocities and their time delayed copies. T: fixed (25) | 0-0.5, 4-10 |
| 19 | 3D motion capture, optoelectronic system, with markers on thoracic spine. 3D | 3D linear accelerations and angular velocities of the trunk | 3x5 min walking trials. 150 strides. | TW | None | Time series normalized to 150 strides | 12-dimensional state space combining linear acceleration and angular | 0-0.5, 4-10 |

COPYRIGHT 2026 THE AUTHORS. THIS FILE IS LICENSED UNDER A CREATIVE COMMONS ATTRIBUTION 4.0 INTERNATIONAL (CC BY 4.0) LICENSE. 

| ID | Measurement system | Signal type | Duration | Walking condition | Filtering method | Resampling method | Attractor reconstruction | Stride range |
|---|---|---|---|---|---|---|---|---|
| | accelerometer and gyroscope fixed on thoracic spine (50Hz) | (thoracic spine), for both measurement systems. | | | | | velocities and their time delayed copies. T: fixed (25) | |
| 20 * | Force-sensitive resistor insole for heel strike detection. 3D accelerometer, lumbar spine (200Hz) | 3D linear acceleration of the trunk (lower back) | 3x15 min walking trials. Average of 3x 100 strides | OG, 34 m circular track | Not mentioned | Not mentioned | T: variable (autocorrelation function) dE: 5 (GFNN) | 0-1, 4-10 |
| 21 | 6 camera 3D optoelectronic system (50Hz) | Angular velocity of sagittal knee movements (bilateral) | 7x2 min walking trials. 30 strides per trial. Repeated at 6 weeks, 6 months and 1 year. | TW | Not mentioned | Time series resampled to on average 100 samples per stride. | T: fixed, 10. dE: 5 methods not mentioned | 0-0.5, 4-10 |
| 22 | 24 camera 3D motion capture system, 22 markers (60Hz). | 3D velocities of the cervical spine. (upper trunk) | 25 (5x5) 3 min walking trials. 150 strides per trial. | TW | Not mentioned | Time-normalized: Resampled to on average 100 samples per stride. | T: fixed for each axis (10-20), AMI dE: 5 (method not mentioned) | 0-1, 4-10 |
| 23 * | 8 camera optical 3D motion capture system. (120Hz) with markers over 5th metatarsal, heel, lateral malleolus, lateral midshank, lateral knee, and lateral mid-thigh. | knee angle, ankle angle, and ankle Y (vertical oscillation) | 10 min walking trial, from which only 3-5 min were analyzed | TW | Not mentioned | Not mentioned | T: variable (AMI) dE: 6 (GFNN) | 0-1, 4-10 |
| 24 | 8 camera 3D motion capture system (60Hz). Markers around major joints. | Flexion-extension angle of neck, shoulders and elbows (bilateral) | 90 s walking. | TW | None | Not mentioned | T: variable (AMI) dE: 5 (GFNN) | 4-10 |
| 25 * | 3D accelerometer fixed to the lower back (200Hz) | 3D trunk accelerations | 10 min OG 10 min TW. 7 min per trial used for DE analysis. | OG and TW | None, except down sampling | 200 Hz downsampled to 100Hz for accelerating computation. No time normalization | T: variable (AMI) dE: 6 (GFNN) | 0-1, 4-10 |
| 26 | 3D inertial sensor (accelerometers, gyroscopes) at lumbar spine (50Hz) | 3D linear accelerations and angular velocities of the trunk (lower back) | 3 walking trials: 3.5 min, 3 min, 2.5 min. 115 strides in each trial. | TW | None | Strides were time-normalized to the average stride length [?] for each speed level. | 12-dimensional state space combining linear acceleration and angular velocities and their time delayed copies. T: constant ¼ stride time. (AMI) | 0-0.5, 4-10 |
| 27 | 10 camera motion capture system (60Hz), markers on trunk and feet. | 3D cervical spine velocities. | 6 x 3 min walking trials. 120 strides | TW | Not mentioned | Time normalized, resampled to 12,000 data points. | T: 20, 15, 10 (axes, method not mentioned dE: ? not mentioned | 0-1, 4-10 |
| 28 | 8 camera motion capture system (60Hz), markers on lower limbs | Plantar flexion and dorsiflexion of the ankle, flexion and extension of the hip | 3 min walking trials. | TW | None | Not mentioned | T: variable (AMI) dE: variable (GFNN) | 0-1, 4-10 |
| 29 * | Force-sensitive resistor insole for heel strike detection. 3D | Time series of heel strikes = stride intervals (unilateral). 3D | 2 sessions of 5x15 min. walking. | OG | Not mentioned | Not mentioned | T:? (Autocorrelation function dE:? (GFNN) | 0-1, 4-10 |

COPYRIGHT 2026 THE AUTHORS. THIS FILE IS LICENSED UNDER A CREATIVE COMMONS ATTRIBUTION 4.0 INTERNATIONAL (CC BY 4.0) LICENSE.

| ID | Measurement system | Signal type | Duration | Walking condition | Filtering method | Resampling method | Attractor reconstruction | Stride range |
|---|---|---|---|---|---|---|---|---|
| | accelerometer fixed to the lower back. (200Hz) | accelerations of the lumbar spine. | | | | | | |
| 30 | 24 camera 3D motion capture tracking a cervical spine marker. (60Hz) | Mediolateral velocity of the cervical spine. | 18 walking trials (2x9) of 3 min. 150 strides per trial. Aggregated results (average of 2 trials) | TW | Not mentioned | Resampled to 15,000 data points ~ 100 per stride | T: 20 (AMI)<br>dE: 5 (GFNN) | 0-1, 4-10 |
| 31 | 3D inertial sensors (accelerometers, gyroscopes) fixed to the thoracic spine. | 3D linear accelerations and 3D angular velocities of the trunk | 7 min. walking trials. 150 strides | TW | low pass filtering (20 Hz, 4th order Butterworth ) | Not mentioned | T: 10 (method not mentioned.<br>dE: 5 (method not mentioned).<br>6 and 15 dimensional state spaces were also used. | 0-0.5, 4-10 |
| 32 | 3D accelerometer fixed to the lumbar spine | 3D linear acceleration of the trunk | 2x2 min walking trials. 80 strides | TW | low-pass filtering (4th order, Butterworth , 20 Hz) | Time normalized: Resampled to 8,000 data points ~ 100 per stride | T: 10 (method not mentioned.<br>dE: 5 (method not mentioned). | 0-1, 4-10 |
| 33 * | 8 camera motion capture, sagittal plane kinematic (120Hz). Markers over the 5th metatarsal, lateral malleolus, heel, lateral shank, lateral knee, and lateral thigh. | Relative phase angle between the thigh and shank and shank and foot. Knee angle, ankle angle. Ankle vertical oscillations. | 3x4 min. A variable number of strides were analyzed. | TW | Not mentioned | None (raw 4 min. data used, uniform number of data points) | T: variable (AMI)<br>dE: 6 (GFNN) | 0-1, 4-10 |
| 34 | Force-sensitive resistor insole (for heal strike detection). 3D accelerometer fixed to the lower back. (200Hz) | 3D accelerations of the trunk. | Exp. 1: 2x(3x15) min. walking.<br>Exp. 2: 2x(3x15) min walking. | OG (Exp.1)<br>TW (Exp. 2) | Not mentioned | Not mentioned | T : ? (AMI)<br>dE : ? (GFNN)<br>Ref to 20 ? | ? Likely 0-1, 4-10 as in 20. |
| 35 * | Instrumented treadmill, with a grid of foot-pressure sensors (100Hz). | 2D center of pressure trajectory. (feet) | (2x3) 5 min walking trials. 175 strides per trial. | TW | Downsampled to 50 Hz (eighth-order low pass Chebyshev Type I filter) | Time-normalized. Resampled to 10'000 points (polyphase interpolation) | T: 15 (ML) and 18 (AP), AMI<br>dE: 6 (GFNN) | 0-0.5, 4-10 |
| 36 | 3D accelerometer fixed to the sternum (200Hz) | 3D acceleration of the trunk | 2x 5 min walking. 6x35 or 6x70 strides. Outcomes aggregated (average) over 6 bouts (210 or 420 strides) | TW | Downsampled to 50Hz in (Chebyshev Type I, zero-phase filtering) | Time-normalized. Resampled to 2,500 or 5,000 points (polyphase interpolation) | T: 5 (AP), 5 (V) and 6 (ML), (AMI)<br>dE: 6 (GFNN) | 0-0.5, 0-1, 4-10 |
| 37 | 2 synchronized 3D inertial sensors fixed to lumbar spine and on the right shank. (128Hz) | 3D acceleration of the trunk | 250 m walking ~180 strides. 150 strides included | OG | Not mentioned | Not mentioned | T: 10 (method not mentioned)<br>dE: 5 (GFNN) | Not mentioned, likely 0-1, 4-10 |
| 38 | 3D inertial sensor fixed to the lumbar spine. (100Hz / 200Hz) | 3D accelerations of the trunk. Gyroscopes | 6 min walking along a 30m pathway, fast walking. 20 | OG | None | Not mentioned | T: 10 (method not mentioned)<br>dE: 5 (GFNN) | Not mentioned, likely 0-1, |

COPYRIGHT 2026 THE AUTHORS. THIS FILE IS LICENSED UNDER A CREATIVE COMMONS ATTRIBUTION 4.0 INTERNATIONAL (CC BY 4.0) LICENSE. 

| ID | Measurement system | Signal type | Duration | Walking condition | Filtering method | Resampling method | Attractor reconstruction | Stride range |
|---|---|---|---|---|---|---|---|---|
| | | used for detecting turns. | strides analyzed. Aggregated values (average). | | | | | 4-10 |
| 39 | 8 camera 3D motion capture system (120H), full body capture (28 markers). Force plates (600Hz) | 3D linear motions and rotations of the trunk. | 7 m instrumented pathway. 20 strides analyzed before slip trials | OG | Not mentioned | Not mentioned | 6D phase space of trunk kinematics (positions and angles) | 0-0.5, 4-10 |
| 40 | 3D motion capture system, rigid body tracking (3 non-collinear markers per segment). | Norm of the 3D angular positions of the lumbar spine, hip, knee, and ankle. | 4 x 5 min walking trials. | TW | low-pass filtered (fourth-order dual-pass Butterworth, 10 Hz cut-off) | Not mentioned | T: 7, 10, 8, and 5 (autocorrelation method) dE: 6, 5, 5 and 6 (GFNN) | 0-0.5, 0-1, 4-10 |
| 41 * | 3D motion capture system (100 Hz) recorded markers placed on sacrum and right and left posterior calcanei. | Time series of right and left heel positions relative to the sacrum position. | 2 x 3 min walking trials (150 strides) | TW | low-pass filtered (fourth-order Butterworth filter, 12 Hz cut-off.) | Not mentioned | T: variable, ¼ of subjects' average stride time. dE: 5 (GFNN) | 0-1, 4-10 |
| 42 | 24 camera 3D motion capture system (60Hz), full body (57 markers) | Cervical spine 3D velocity. | 3 x (4 x 3 min) walking trials. 124 strides. | TW | Zero-lag Butterworth low-pass filter, 8 Hz. | Time normalized, resampled to 12,400 data points. | T: 15, 10 and 30 (AMI) dE: 5 (method not mentioned) | 0-1, 4-10 |
| 43 | 3D accelerometer (200Hz), sternum (upper trunk) | Mediolateral acceleration of the upper trunk | 3 x 6 min walking trials.175 strides | TW | Preliminary down sampling to 50Hz | Time normalized, resampled to 10,000 data points. | T: 6, average AMI results. dE: 6 (GFNN) | 0-0.5, 4-10 |
| 44 | 3D inertial motion unit, thoracic spine, 200Hz. 12 camera 3D motion capture (200Hz), lateral malleoli marker. | Vertical acceleration signal of the thoracic spine | 60 s of walking. | TW | Zero-lag low-pass filtered at 28 Hz cut-off with a digital fourth-order Butterworth filter. | each stride was resampled to 100 data points (? method non clear) | T:10 dE: 5 Methods not mentioned | 0-0.5, 4-10 |
| 45 | 3D accelerometer fixed to the right hip (50Hz) | Vector norm of the 3D acceleration signal (trunk, pelvis level) | Unconstrained walking: variable number of 1 min walking bouts over one week. 35 strides. Aggregated average values over on average 77 1min bouts (patients) and 34 1min bout (controls) | OG (free-living) | None | Time-normalized. Resampled to 2500 samples | T: 8 (AMI) dE: 5 (GFNN) | 0-0.5, 2-6 |
| 46 | 3D accelerometer, lumbar spine (296 Hz) | 3D acceleration of the lumbar spine. | 5 min OG walking, 25m corridor. 3 min TW. 140 strides (U-turn removed) | TW and OG | 2$^{nd}$-order low-pass Butterworth filter, 50 Hz cut-off. | Time-normalized, resampled to 14,000 data points. | T: 10, 7, 9 (AMI) dE: 6 (GFNN) | 0-0.5 4-10 |
| 47 | 14 camera 3D motion capture system (100 Hz). | 3D velocity of the cervical spine. | 4 x 2 min walking. | TW | 4$^{th}$-order zero-lag low-pass | Not mentioned | T: ¼ of average stride time. dE: 4 (GFNN) | 0-1, 4-10 |

COPYRIGHT 2026 THE AUTHORS. THIS FILE IS LICENSED UNDER A CREATIVE COMMONS ATTRIBUTION 4.0 INTERNATIONAL (CC BY 4.0) LICENSE.

| ID | Measurement system | Signal type | Duration | Walking condition | Filtering method | Resampling method | Attractor reconstruction | Stride range |
|---|---|---|---|---|---|---|---|---|
| | Markers on heels, posterior sacrum, and C7 vertebrae. | | | | Butterworth filter, 8 Hz. | | | |
| 48 * | 3D accelerometer fixed to the sternum (200Hz) | 3D acceleration of the trunk | 2x 5 min walking. 250 strides. | TW | None | Time-normalized. Resampled to 25,000 data points. | T: variable ~7 (AMI)<br>dE: 5 (GFNN) | 0-0.5<br>2-4<br>4-10 |
| 49 | Three 3D accelerometers attached to the lumbar spine and both shanks. 296 Hz. | 3D acceleration of the trunk and shanks | 5 min walking. 25m corridor. 150 strides (U-turn removed) | OG | None | Time normalized. Resampled to 15,000 data points. | T: 11, 8 and 10 (trunk) and 9, 6 and 11 (shanks)<br>dE: 6 (GFNN) | 0-0.5<br>4-10 |
| 50 | 3D motion analysis system (75 Hz) capturing right leg kinematic in sagittal plane. | Knee-joint flexion-extension movement. | 75 strides | TW | None | Not mentioned | ? T: 17<br>dE: 2<br>(not clearly stated) | 0-0.5<br>4-10 |
| 51 * | Instrumented treadmill. Force platform (500Hz) | Center-of-pressure trajectory (feet), anteroposterior (AP) and mediolateral (ML) directions | 3 x 10 min walking. 500 strides | TW | low-pass filtered (18 Hz 12th order Butterworth) and down-sampled to 100 Hz. | Time-normalized. Resampled to 50,000 data points. | T: variable (AMI)<br>dE: 5 (GFNN) | 0-0.5<br>1-4<br>4-7<br>7-10 |
| 52 | Instrumented treadmill, with a grid of foot-pressure sensors (? 100Hz). | Center-of-pressure trajectory (feet) | 4 x (6 x 30 s) walking trials. | TW | Not mentioned | Not mentioned | T: variable (autocorrelation)<br>dE: ? | 0-0.5<br>4-10 |
| 53 | 10 camera 3D motion capture system (100 Hz), 16 markers on the legs. Instrumented treadmill capturing 3D ground reaction force (1000 Hz) | Hip, knee and ankle kinematic. | 5x3 min walking. 100 strides, average of 3 consecutive blocks of 33 strides. | TW | Not mentioned | Not mentioned | T: ~35 (AMI)<br>dE: 5 (GFNN) | 0-1,<br>4-10 |
| 54 * | 7 camera 3D motion capture (100Hz). Markers on trunk and lower body. | 3D incremental (difference) time series of the cervical spine. | 4 x 5 min walking trials 241 strides | TW | Not mentioned | Each stride was resampled to 101 data points (? method non clear) | T: 17, 11, 8 (Autocorrelation)<br>dE: 5 (GFNN) | 0-1,<br>4-10 |
| 55 | 6 camera 3D motion capture system (100 Hz), whole body markers (15).<br>6 electrodes EMG | 3D of the COM trajectory (trunk), angles of the pelvis, hip, knee and ankle joints. EMG data of the Gluteus Medius (GM), Rectus Femoris (RF), Vastus Medialis (VM), Biceps Femoris (BF), Medial Gastrocnemius (MG), Tibialis Anterior (TA). | 6 x 3 min walking trials. 150 strides | TW | The EMG data were band-pass filtered between 30 and 450Hz using a 4th order Butterworth recursive filter. | Time series normalized, resampled to 15'000 data points | T: variable, medians 23 (kinematic) and 29 (muscle activities). AMI<br>dE: variable, 5-8 kinematic, 7-10 muscle activities. GFNN. | 0-0.5<br>4-10 |
| 56 | 13 camera 3D motion capture system (120 Hz). | Kinematics of the pelvis (lower back) and lower | 3 x 7 min walking trials. 26 strides captured | OG (indoor rectangle circuit) | 2nd-order, low-pass, recursive, Butterworth | Not mentioned | T: variable ? (Autocorrelation)<br>dE: variable ? (GFNN) | 0-1,<br>4-7 |

COPYRIGHT 2026 THE AUTHORS. THIS FILE IS LICENSED UNDER A CREATIVE COMMONS ATTRIBUTION 4.0 INTERNATIONAL (CC BY 4.0) LICENSE.

| ID | Measurement system | Signal type | Duration | Walking condition | Filtering method | Resampling method | Attractor reconstruction | Stride range |
|---|---|---|---|---|---|---|---|---|
| | Markers on pelvis and lower limbs | limbs (left side). Hip, knee, and ankle joint angle time series in the sagittal plane | (straight section) | | filter with a cutoff frequency of 10 Hz | | | |
| S1 | 7 camera 3D motion capture system (200 Hz). 20 whole body markers. Instrumented treadmill (two force plates, 1kHz) | Ground reaction force (GRF) for gait event detection (toe off). Center of mass (COM) 3D velocities of whole body, trunk, thigh, shank and foot. Hip, knee and ankle joint in the sagittal plane. | 3 x 10 min walking trials. 450 strides captured. | TW | 4th-order low-pass Butterworth filter with zero lag and cut-off frequencies of 10 Hz | Time normalized, each stride was resampled to 100 data points (? method not clear) | T: variable (AMI)<br>dE: 6 (GFNN) | 0-0.5, 4-10 |
| S2 * | 3D accelerometer fixed to the lower back | 3D acceleration of the trunk | 4 x 205 m walking trials. 125 strides captured. | OG, indoor circuit | None | Time-normalized. Resampled to 18'750 data points. | T: variable (AMI)<br>dE: 5 (GFNN) | 0-0.5, Variable for long-term DE |
| S3 | 14-camera real-time 3D motion capture system (120 Hz) | Center of mass (COM) and center of pressure (COP) trajectories. COM movement with respect to COP movement in the mediolateral direction. Position and speed. | 5 x 10 m. 13 strides captured, 10 strides analyzed. | OG, 10 m walkway | 4th order lowpass Butterworth filter with a 10 Hz cutoff frequency | Not mentioned | T: variable (AMI)<br>dE: different method used (study aim) | 0-0.5, 6-8 |
| S4 * | 3D accelerometer fixed to the lower back | 3D acceleration of the trunk | 4 x 205 m walking trials. 125 strides captured. | OG, indoor circuit | None | Time-normalized. Resampled to 18'750 data points. | T: variable (AMI)<br>dE: 5 (GFNN) | 0-0.5, 5-12 |
| S5 | 5 inertial sensors in the lower extremities and lower back (100Hz) | Gait event from shank sensor. 3D acceleration signals of the lower back | 7 x 6 walking trials on different surfaces. 40 strides (concatenated) | OG, outdoor | None | Not mentioned | T: ? (AMI)<br>dE: ? (GFNN) | 0-1, 4-10 ? From references |
| S6 | 12-camera 3D motion capture system (220 Hz). 3D inertial sensor fixed to the lower back | joint angles (sagittal, frontal, transverse), and CoM and trunk accelerations | 4 x 2 min walking trials. Number of strides not mentioned | TW | Motion-capture data were low-pass filtered at 14 Hz and IMU data at 20 Hz (fourth-order Butterworth) | Not mentioned | T: ? (AMI)<br>dE: ? (GFNN) | ? likely standard 0-1 and 4-10 |

* Study using detrended fluctuation analysis (DFA) to quantify the long-range correlation structure of stride intervals (gait complexity).

COPYRIGHT 2026 THE AUTHORS. THIS FILE IS LICENSED UNDER A CREATIVE COMMONS ATTRIBUTION 4.0 INTERNATIONAL (CC BY 4.0) LICENSE.

#### D. Study aims, result interpretation, and conclusion

| ID | Study aims | DE results | DE interpretation | Key conclusion | Clinical relevance |
|---|---|---|---|---|---|
| 1 | To test if diabetic neuropathy patients have increased local instability compared to controls and if slower walking speeds are a compensatory strategy for stability. | Lower long-term DE in neuropathic patients | Slower walking speeds in neuropathic patients enhance local dynamic stability (lower DE = higher stability). | Reduced walking speed is an adaptive strategy for maintaining upper body stability in neuropathic patients, not a direct result of sensory loss. | Highlights the importance of considering speed as a compensatory strategy in clinical assessments of gait stability. |
| 2 | Compare local dynamic stability of normal and neuropathic gait under overground vs. treadmill conditions | TW reduced long-term DE by 3–18%; diabetic neuropathic (NP) patients showed lower DE values. | Lower long-term DE indicates greater local stability; NP patients adopt slower walking speeds as a compensatory strategy. | Variability and local dynamic stability are distinct concepts; NP patients improve local stability by slowing down | Insights into compensatory strategies in neuropathic gait can guide fall prevention and rehabilitation programs. |
| 3 | Demonstrate that local dynamic stability (DE) emerges over multiple consecutive strides and differs from traditional variability measures | Show that small perturbations continue to diverge for over ten strides, with TW reducing both short- and long-term DE | Indicate that DEs capture different and more nuanced aspects of locomotor control compared to standard gait variability measures | TW can artificially decrease variability and increase stability, potentially obscuring true gait differences relative to OG | Clinicians and researchers should be cautious when interpreting treadmill-based gait data in populations with altered neuromuscular control |
| 4 | Investigate how a 3-hour free-walking task affects gait, muscle activity (EMG), heart rate, and local dynamic stability under fatigue. | Long-term DE decreased in group exhibiting higher subjective fatigue but increased in group with lower fatigue. | Decrease in DE = reduced dynamic stability under fatigue. Increase in DE = distinct adaptation mechanism in less fatigue-prone individuals. | Fatigue-induced muscle changes drive instability, then slower gait to regain stability. Low fatigue group is better to maintains stability, showing different fatigue adaptation. | A new physical fatigue index (combining gait + physiological data) reliably predicts subjective fatigue. Useful for monitoring fatigue in daily life. |
| 5 | Examine how kinematic variability and local dynamic stability vary across walking speeds to resolve paradox between slow walking and stability. | Short-term DE increased linearly with speed in all directions<br>Long-term DE increased linearly with speed in AP and VT directions. | Lower DE values indicate better local dynamic stability<br>Measures resilience of locomotor system to natural kinematic variability. | Slower walking improves local dynamic stability despite increased movement variability, resolving the speed-stability paradox. | Explains why slower walking may be a beneficial adaptation despite increased variability |
| 6 | Determine minimal walking duration required for reliable local dynamic stability measures. | Short-term DEs showed good reliability with 2–3 minutes; long-term DE reliability remained low with 5 minutes. | Short-term DE reflects local dynamic stability; Long-term DE requires longer or multiple trials for reliable assessment | 2–3 minutes suffice for short-term stability metrics, but 5 minutes not enough for long-term | Short-term metrics can be collected quickly in clinical settings, aiding assessments for patients unable to walk for long periods |
| 7 | Compare DEs and orbital (Floquet multipliers) stability during treadmill vs. overground walking. | Positive DEs in all conditions, indicating persistent local instability. | DEs are sensitive to small perturbations but does not predict overall (orbital) gait stability | Gait remains orbitally stable despite local instability; treadmill walking slightly improves orbital measures. | None |
| 8 | Examine whether older adults increase gait stability by walking slower.<br>Assess roles of strength and range of motion in stability. | Older adults: higher short-term DE at all speeds, no difference for long-term DE.<br>Both groups reduced short- and long-term DEs when walking slower. | Higher DEs indicate greater local instability.<br>Slower speeds decreased these instability measures in both age groups. | Older adults remained more unstable even after controlling for strength and flexibility.<br>Slower walking reduced instability but did not eliminate age effects. | Gait stability measures may serve as sensitive indicators of fall risk.<br>Speed alone is insufficient to predict future fall risk. |
| 9 | Examine effects of peripheral neuropathy (PN) on gait variability and local instability at different speeds | Short-term DE significantly higher for PN group at fast walking speed. No significant difference for long-term DE. | Elevated DEs reflect increased local instability and heightened sensitivity to perturbations. | PN leads to greater variability and instability at faster speeds, implying reduced adaptability. | Slower walking speeds may serve as a compensatory strategy to minimize fall risk. |

COPYRIGHT 2026 THE AUTHORS. THIS FILE IS LICENSED UNDER A CREATIVE COMMONS ATTRIBUTION 4.0 INTERNATIONAL (CC BY 4.0) LICENSE. 

| ID | Study aims | DE results | DE interpretation | Key conclusion | Clinical relevance |
|---|---|---|---|---|---|
| | | When collapsed across speeds, NP patients exhibited higher short-term DE and lower long-term DE as compared to healthy control (not significant) | | | |
| 10* | Examine relationships between long-range correlations (DFA), local stability (long-term DE), and relative phase variability during gait transitions. | Head long-term DE increased during slow running vs fast walking. Ankle DE decreased during slow running vs fast walking. Greater long-term DE at head vs ankle across conditions. Significant correlations between long-range correlations (DFA) and long-term DEs. | Higher long-term DE indicates decreased local dynamic stability. Head stability prioritized over ankle stability. Different stability strategies for head vs ankle control. | Relevant relation between long range correlations in the stride interval of walking (DFA) and measures of stability (long-term DE). | None |
| 11 | Determine required data length for precise estimates of dynamic stability measures and assess sensitivity to walking speed and dual-task changes. | DEs increase with longer data series. Short-term DE sensitive to speed changes in AP/VT directions and Stroop task. Long-term DE only sensitive to speed changes. | Higher DEs indicate less stable patterns. Precision improves with longer data series. Different sensitivity between short-term and long-term DEs. | 150+ strides needed for reliable measures. Short-term and long-term DEs respond differently to gait changes. Walking speed affects both measures while cognitive task only affects short-term DEs. | Highlights need for standardized stride counts in clinical gait studies. |
| 12 | Examine how walking speed influences DEs and variability of trunk movements. | Short-term DE decreased linearly with speed in AP direction; showed inverted U-shaped patterns in ML and VT. Long-term DE had distinct linear or quadratic effects per direction. | DE depends on walking speed and direction of measurement. Slow walking is not consistently more stable than fast walking. | Speed-related changes in stability are direction-specific; slow gait is not unequivocally more stable. | Challenges assumption that slower walking is always better for balance. Great caution should be exerted to avoid methodological pitfalls. |
| 13 | Compare dynamic stability of trunk vs. pelvis, thigh, shank, foot. Examine age differences between younger and older adults. | Older adults show higher short-term DEs than younger. The effect is less evident for long-term DE. Trunk has smaller DEs than distal segments. | Larger DEs in older adults mean reduced dynamic stability. | Trunk stability is prioritized; trunk motion most sensitive to age-related declines in dynamic stability. | Assessing trunk dynamics may detect early gait impairments in older populations. |
| 14 | Compare continuous multi-muscle activation dynamics (EMG) and kinematics in young vs. older adults at different walking speeds. | Older adults showed larger short-term DEs than younger controls. Not significant difference for long-term DE. Both long-term and short-term DEs measured from EMG and kinematics were highly correlated. | Greater values reflect heightened instability or sensitivity to small perturbations in muscle activation. | Neuromuscular activation patterns are mirrored in the observed kinematics, indicating a coupling between muscle activity and gait dynamics. | Early detection of subtle neuromuscular instability may help identify older adults headed toward gait-related functional declines. |
| 15 | Examine effects of acute plantar sensation loss on stability-related gait kinematics at different speeds | Ice-induced desensitization increased short-term DEs and long-term DEs. Higher speeds led to greater DEs. | Higher DEs indicated a diminished ability to attenuate small-scale gait perturbations. | Acute plantar sensory loss raises local instability but does not alter magnitude of spatial or temporal variability. | Supports importance of plantar feedback for maintaining stable gait control; relevant for neuropathic populations |
| 16 | Examine effects of side-by-side walking on stride-to-stride variability and kinematic coordination between pairs of individuals synchronizing their gait, voluntarily or not. | While short-term divergence DE was consistent across conditions, long-term DE rose significantly during unintentional synchronization. | Suggests that changes in DEs might represent alterations in the locomotor attractor’s properties rather than direct changes in stability | Side-by-side walking affects gait stability and coordination patterns. | Potential therapeutic application for gait rehabilitation through interpersonal synchronization. |
| 17 | Determine kinematic variability of lower extremity joints using chaos | Long-term DE values: Males: 0.035 ± 0.016 (right ankle) to 0.073 ± | Larger DEs indicate more divergence and variability. Positive DE | Significant LE differences between joints except hip- | Normative long-term DE values for lower extremity joints. |

COPYRIGHT 2026 THE AUTHORS. THIS FILE IS LICENSED UNDER A CREATIVE COMMONS ATTRIBUTION 4.0 INTERNATIONAL (CC BY 4.0) LICENSE. 

| ID | Study aims | DE results | DE interpretation | Key conclusion | Clinical relevance |
|---|---|---|---|---|---|
| | theory in healthy young adults during normal walking. Test hypothesis of variability differences between joints, genders, and sides. | 0.023 (left knee). Females: 0.028 ± 0.014 (left ankle) to 0.065 ± 0.028 (right hip). | suggests chaotic characteristics. Lower positive DEs imply less sensitivity to perturbations. | knee. No significant right-left differences. Only left knee showed significant male-female differences. | Potential for assessing gait abnormalities, evaluating recovery progress, fall risk assessment. Sex have little effects on long-term DEs. |
| 18 | Investigate effects of arm swing and walking speed on gait stability using nonlinear measures. | Arm swing slightly increases short-term DE and decreases long-term DE (not significant). Walking speed affected DEs differently, lowering short-term values but increasing long-term ones. | Short-term DE quantifies the system's ability to respond to small perturbations during walking, with higher values indicating decreased local dynamic stability. | Arm swing enhances gait stability through improved responses to perturbations and more efficient recovery strategies. | Understanding arm swing's role in gait stability could inform rehabilitation strategies and fall prevention programs. |
| 19 | Compare gait stability measures from optoelectronic and inertial sensor systems during treadmill walking. | Short-term DE decreased with speed, long-term DE increased, high correlation (R > 0.85) between systems for DEs. | Short-term DE reflects short-term local stability; long-term DE possibly linked to stride variability. | Inertial sensors are a valid alternative to optoelectronic systems for assessing dynamic gait stability. | Use of inertial sensors enables large-scale studies on gait stability in natural walking conditions. |
| 20* | Compare ability of gait measures (stride interval dynamics, DEs) to detect changes between overground and compliant surface walking. | Long-term DE showed significant differences between compliant surface and overground conditions. Lower values on compliant surface. | Unexpected lower long-term DE suggests improved local dynamic stability, possibly due to increased stride intervals and cautious walking strategy. | Long-term DE can distinguish compliant surface from overground walking, unlike traditional gait variability measures. | None |
| 21 | Investigate knee kinematics' stability and variability during gait in knee osteoarthritis patients pre- and post-replacement surgery. | Short-term DE decreased with speed and long-term DE increased with speed for both patients and controls.<br>Pre-operatively, patients had higher short-term DE at the unaffected side compared to healthy controls, and lower long-term DE at the affected side compared to healthy controls. Post-operatively, no change in long-term DE was observed, while short-term DE decreased at unaffected side over time. | Higher unaffected short-term DE in patients suggests reduced local stability on the unaffected side, possibly due to altered gait patterns to reduce demands on the affected leg.<br>Lower affected long-term DE in patients may indicate a compensatory strategy to improve global stability on the affected side. | Knee replacement surgery improves both stability and variability of knee kinematics during gait in OA patients. | Highlights the importance of assessing dynamic stability for evaluating surgical outcomes and rehabilitation strategies. |
| 22 | Determine if continuous pseudo-random perturbations (mechanical and visual) induce measurable changes in dynamic stability measures during walking. | Long-term DE increased during all perturbation conditions<br>Long-term DE decreased during all perturbation conditions | Short-term DE measure of local dynamic instability of the gait. Long-term DE may indicate how quickly movements reach maximum divergence limits. | Subjects showed decreased orbital and short-term local dynamic stability specifically in directions of applied perturbations, while long-term stability appeared to improve. | Metrics like DE could help assess fall risk and inform rehabilitation strategies. |
| 23* | Examine effects of resistance training-induced fatigue on nonlinear dynamics of lower limb kinematics during treadmill walking | No significant changes in short-term and long-term DEs for ankle angle, knee angle, and vertical ankle movement post-fatigue. | DE values interpreted as measures of local dynamic stability of gait patterns | Control of walking is robust to moderate muscle fatigue in healthy individuals | None |
| 24 | Quantify local stability of neck and upper extremities during walking using long-term DE. Compare linear (ROM) and nonlinear (DE) measures and analyze differences between upper and lower extremities. | Long-term DE values: Neck 0.037±0.0230. R Shoulder 0.043±0.0210. L Shoulder 0.045±0. 0300.R<br>Elbow 0.032±0.0210. L Elbow 0.034±0.0260. No significant differences between joints or genders. | DE reflects local stability of joint motion, with higher values indicating lower stability | Long-term DE for upper extremities is lower than for lower extremities, suggesting greater stability in upper extremity joints during walking. DE and ROM provide complementary | None |

COPYRIGHT 2026 THE AUTHORS. THIS FILE IS LICENSED UNDER A CREATIVE COMMONS ATTRIBUTION 4.0 INTERNATIONAL (CC BY 4.0) LICENSE. 

| ID | Study aims | DE results | DE interpretation | Key conclusion | Clinical relevance |
|---|---|---|---|---|---|
| | | | | insights into joint motion analysis. No differences between genders | |
| 25* | Compare treadmill vs overground walking using variability measures: kinematic variability (SD), fractal dynamics (DFA), local dynamic stability (LDS) | Treadmill significantly decreased short-term DE in all directions and long-term DE in ML and V directions but not AP. | Higher stability (lower DEs) on treadmill may be due to reduced degrees of freedom in constrained environment. | Treadmill modifies non-linear gait characteristics (DE, long-range correlations) but not kinematic variability | Choice of walking condition (treadmill vs overground) important to consider in protocol design |
| 26 | Test sensitivity of trunk variability and stability measures to experimentally induced balance impairments using galvanic vestibular stimulation (GVS) during treadmill walking. | Short-term DE increased significantly with GVS across all walking speeds, while long-term DEs decreased. | Short-term DE reflects reduced local stability due to induced perturbations. Long-term DE unexpectedly decreased, possibly reflecting compensatory mechanisms or insensitivity to short-term perturbations. | Short-term DEs and variability measures are reliable indicators of gait stability impairments. Long-term DEs and Floquet multipliers are not suitable for assessing balance impairments during gait. | Portable system using trunk acceleration variability and short-term DE could diagnose gait stability issues |
| 27 | Assess impact of voluntary step width/length modifications on gait variability and trunk dynamic stability | Wide steps increased short-term DE in all directions; narrow steps decreased ML but increased AP and V short-term DE; long steps increased AP/VT short-term DE, but decreased it in ML axis. Short steps increased ML and V short-term DE but decreased it in AP direction. Wide and long steps increased ML long-term DE; Narrow steps increased long-term DE in AP axis. Shorter steps decreased AP and V long-term DE. Longer steps increased long-term DE in all axis. | Higher DE values denote greater local dynamic instability and sensitivity to small perturbations. | Voluntary gait alterations do not improve stability; step modifications increase variability and can worsen trunk dynamic stability | Cautious gait patterns (wider/shorter steps) may not protect against falls. |
| 28 | Compare Wolf's and Rosenstein's algorithms for DE estimation using small gait datasets. | Short-term and long-term DE revealed higher local instability in older adults only for the 3-minute time series. | With Rosenstein algorithm, averaging of convergent and divergent trajectories led to underestimation of instability, especially in short data. | Wolf's algorithm is more sensitive for short gait data; Rosenstein's needs longer data to detect group differences in stability | Extended data collection is important when using Rosenstein's method to detect subtle gait stability changes |
| 29* | Examine how auditory, visual, and tactile cues modify fractal gait dynamics and dynamic stability in healthy adults. | No significant changes in short-term DEs under any cue conditions. Auditory and combined cues significantly lowered long-term DEs. | Lower long-term exponents suggest more stable gait over extended stride sequences. Observe that change in long-term DEs coincide with alterations in stride-to-stride fluctuations (lower scaling exponent, DFA). | Auditory cues have the strongest impact, reducing stride variability and fractal persistence simultaneously. | Visual cues could be harnessed in rehabilitation without detrimental effects on normal fractal rhythms. |
| 30 | Determine how orbital and local dynamic stability metrics change in response to visual and surface perturbations of different amplitudes | Short-term DE increased significantly, while long-term DE decreased significantly, for all perturbation conditions compared to unperturbed walking. | Short-term increase indicates greater local instability; long-term decrease reflects bounded divergence. | Perturbation type affects stability more than perturbation magnitude. | Nonlinear metrics can assess fall risk without inducing falls; useful for rehabilitation and fall prevention. |
| 31 | Explore collinearity of gait variability and stability measures. Assess association between gait | Short-term DE positively associated with fall history. Significant in | Short-term DE quantifies divergence rate after small perturbations within 1 | Both increased gait variability and short-term instability are | Short-term DE may help identify fall risk in relatively young elderly (50-75 years). |

COPYRIGHT 2026 THE AUTHORS. THIS FILE IS LICENSED UNDER A CREATIVE COMMONS ATTRIBUTION 4.0 INTERNATIONAL (CC BY 4.0) LICENSE.

| ID | Study aims | DE results | DE interpretation | Key conclusion | Clinical relevance |
|---|---|---|---|---|---|
| | variability, local dynamic stability, and fall history in elderly | multivariate model with gait variability. Long-term DE Not significantly associated with fall history. | step. Long-term DE quantifies divergence between 4-10 strides, less relevant for fall risk. | associated with fall history in elderly. | Could contribute to selection for fall prevention programs if verified prospectively. |
| 32 | Apply local dynamic stability measures to load carrying tasks. Investigate adaptive gait stability changes during load carrying | Higher short-term DE in AP axis only with load. No differences in ML and VT axes. Long-term DEs significantly higher in all axes with load. | Decreased local dynamic stability when carrying load, particularly for long-term dynamics. | Load carrying reduces dynamic stability across all movement axes over longer time periods. | Method can assess fall risk during occupational load carrying using wearable sensors. |
| 33* | Compare gait behavior during paired walking with synchronized and desynchronized patterns. | Short-term DE: Increased in DeSYNC condition for ankle vertical displacement and shank/foot relative phase. Long-term DE: No significant differences between SOLO, PAIRED, and DeSYNC conditions. | Increased DE indicates greater attractor divergence and reduced local dynamic stability. May reflect active search for synchronization with partner | Greater differences in walking patterns between partners lead to more gait modifications Paired walking affects gait variability and complexity, especially when patterns differ. | May inform gait rehabilitation strategies using partner-based interventions. |
| 34 | Investigate effects of music and TV on gait dynamics in healthy adults. | No significant differences across conditions for short-term and long-term DEs. | The lack of significant differences in Lyapunov exponents suggests that music and TV do not alter the local stability of gait | Listening to music did not significantly affect intrinsic walking dynamics. Treadmill walking while watching TV with subtitles resulted in a less persistent gait with lower variability compared to watching TV with sound. | The study provides insights into how external stimuli might affect gait in healthy populations, which could inform rehabilitation strategies for individuals with gait impairments. |
| 35* | To analyze the effect of rhythmic auditory cueing (RAC) on dynamic stability in healthy individuals during treadmill walking. To compare the responsiveness of short-term and long-term DE to RAC. To evaluate the correlation between LDS and statistical persistence in gait parameters. | RAC slightly decreased short-term DE, particularly at slower speeds. RAC significantly decreased long-term LDS across all speeds and directions. | Short-term DE reflects rapid, automated motor processes for managing small perturbations. Long-term DE indicates a more cautious, voluntary control of gait, likely due to the conscious effort to synchronize with RAC. | RAC induces a more dampened dynamics in gait, lowering long-term DE and reducing statistical persistence in stride-to-stride fluctuations. Both phenomena may be manifestations of a more conscious/voluntary gait control. | RAC could be a valuable and safe treatment in gait rehabilitation, potentially reducing fall risk by improving gait stability. |
| 36 | Investigate intra-/intersession reliability of treadmill gait stability using DEs; compare measurement lengths (35 vs 70 strides) and time scales (short-term and long-term DEs) | Short-term DE assessed at the step scale demonstrated higher reliability than stride-scale measures, particularly in the medio-lateral direction. Intrasession agreement was strong, while intersession reliability was moderate. Long-term DE exhibits a 40% increase when measurement length doubles from 35 to 70 strides, Long-term DE exhibited poorer reliability across sessions, even with longer recordings (70 strides). | Short-term DE (step scale) seems a robust indicator of gait stability due to superior repeatability. Long-term LDS very likely requires prolonged gait data and is prone to higher intra and inter session variability. | The findings highlight short-term DE assessed over one step as the most practical metric for clinical applications due to its superior reliability, while long-term LDS remains less reliable for individual assessments. | Short-term DE may be able to detect clinically meaningful stability changes (e.g., fall risk) in vulnerable populations. Repeated short tests offer practical utility for patients with limited mobility. |
| 37 | Determine minimum strides & within-session reliability of 11 gait variability/stability | Short-term DEs showed average reliability (ICC=0.68-0.69). Long-term DEs showed poor reliability, requiring | Short-term DE captures short-term local stability, moderately reliable with ≥85 strides. Long-term DE reflects | Multiscale entropy (MSE) & recurrence analysis (RQA) demonstrated excellent reliability | Recommend MSE/RQA for fall risk assessment using wearable sensors. |

COPYRIGHT 2026 THE AUTHORS. THIS FILE IS LICENSED UNDER A CREATIVE COMMONS ATTRIBUTION 4.0 INTERNATIONAL (CC BY 4.0) LICENSE. 

| ID | Study aims | DE results | DE interpretation | Key conclusion | Clinical relevance |
|---|---|---|---|---|---|
| | measures from trunk accelerations. | >110 strides. ICC=0.41-0.53. | long-term divergence, unreliable even with long trials. | with <10-30 strides - preferable for clinical gait analysis over DE-based measures. | |
| 38 | Assess the influence of directional changes & sampling frequency on gait measures | Turns did not influence short-term DE. Long-term DEs not analyzed (high variability for the small number of strides) | Short-term DE robustness suggests local dynamic stability maintained during turns. | Only Harmonic Ratio affected by turns & sampling frequency; other metrics are reliable. | Short-term DE/MSE/RQA applicable in free-walking fall risk assessment |
| 39 | Assess predictive power of stability indices for slip-induced falls in older adults. | Short-term DE and long-term DE did not differ significantly between fallers and recoverers | Both short- and long-term DEs failed to predict slip-induced falls, likely due to their inability to capture large-scale perturbations. | Feasible-Stability-Region (FSR) measurement was the best predictor of falls, while step width variability showed moderate predictive capacity. | FSR measurement and step width variability could help identify older adults at risk of falling, guiding early interventions. |
| 40 | Evaluate joint stability responses to segment load perturbations during treadmill walking; assess spatial and temporal effects. | Short-term DE increased when a load was added proximal to a joint. Long-term DEs were highly variable; their results offered limited interpretability and were not analyzed in detail. | Elevated DEs signify decreased dynamic stability and less effective neuromuscular control. | Segmental loads proximal to a joint destabilize it; distal loads maintain or stabilize it. | Implications for weight training, orthotics/prosthetics design, and load carriage techniques to maintain/enhance joint stability. |
| 41* | Investigate age-related reliance on visual feedback for balance during walking using virtual-reality (VR) perturbations and dynamic gait metrics. | Old adults: higher short-term DE during perturbations vs. young. No age difference in normal walking. Lower long-term DE during perturbations, but not significant. | Higher short-term DE in old indicates reduced resilience to perturbations and heightened visual reliance. | Older adults depend more on visual feedback for gait stability, compensating for degraded somatosensation. | VR-based perturbations could diagnose sensory deficits; informs balance training targeting visual reliance in elderly. |
| 42 | Evaluate reliability and minimal detectable change (MDC) of gait stability measures during cognitive, physical, visual perturbations. | Short-term DE: higher reliability (ICC 0.59–0.88) and lower MDC (3.9–7.8% of mean). Long-term DE: Lower reliability (ICC 0.49–0.91) and higher MDC (16.6–49.1% of mean). | Short-term DE detect immediate instability; long-term less reliable for tracking changes in individuals. | Temporal-spatial measures and short-term DE reliable for tracking gait changes across perturbations. | Validates measures to identify instability in clinical populations (e.g., elderly, injured) during perturbed walking. |
| 43 | Assess dynamic stability during blindfolded treadmill walking under safe conditions. | Marginal short-term DE increase (1%) during blindfolded vs. eyes-open at same speed. Significant long-term DE decrease (20%) during blindfolded walking vs. control, but marginal not significant change of blindfolded vs. eyes-open at same speed. | Visual deprivation did not impair gait stability | Safe environments enable alternative sensory strategies to stabilize gait without vision. | Highlights importance of safety/context in gait stability assessments for low-vision patients. |
| 44 | Investigate associations between step duration variability and IMU-derived gait characteristics. Assess agreement between resultant and vertical acceleration-derived metrics. | Short-term DE negatively associated with MSE (entropy) at coarseness level 2. Long-term DE not significantly associated with other gait characteristics. | Higher Lyapunov exponents indicate poorer gait stability. | MSE captures multiple gait characteristics, including variability and stability. Resultant acceleration is a valid alternative to vertical acceleration for gait analysis. | Resultant acceleration simplifies IMU-based gait assessments, enabling practical use in clinical settings without requiring sensor orientation estimation or gyroscope data. |
| 45 | Validate wearable accelerometer method for ecological gait assessment in chronic pain patients. | Short-term DE slightly higher in patients (4% difference). Long-term DE substantially lower in patients (19% difference) | Short-term DE: reflects dynamic stability; Long-term DE: likely linked to gait automaticity and control, with lower values indicating a loss of automaticity. (antalgic gait). | Chronic pain patients show altered gait patterns; method is valid and reliable for group-level studies. | Gait monitoring with accelerometers provide complementary data to clinical tests. |
| 46 | Compare gait stability/variability between overground/treadmill | Overground walking shows elevated short- | Higher short-term DE during overground walking suggests a | Both linear and advanced computational | Understanding differences between OG and TW is crucial |

COPYRIGHT 2026 THE AUTHORS. THIS FILE IS LICENSED UNDER A CREATIVE COMMONS ATTRIBUTION 4.0 INTERNATIONAL (CC BY 4.0) LICENSE.

| ID | Study aims | DE results | DE interpretation | Key conclusion | Clinical relevance |
|---|---|---|---|---|---|
| | walking in young/older women using linear/nonlinear measures | term DE in all directions across both age groups. Lower long-term DE in overground in V/ML direction in young. Only in ML axis in older. Higher long-term DE in older during overground as compared to young in V direction. | greater need to respond to immediate perturbations. Lower long-term LE during overground walking might be due to visual anticipation of movement. | techniques are sensitive to walking conditions (TW), especially in older adults. | for accurate gait assessment and rehabilitation, particularly when using advanced computational techniques. Findings highlight the importance of considering ecological validity in gait analysis. |
| 47 | Investigate whether local dynamic stability during unperturbed walking predicts perturbation response; examine aging/falls history effects. | Optical flow perturbations markedly increased short-term DE; older fallers showed higher short-term DE than young adults. Long-term DE remained mostly unchanged with perturbations; group differences were minimal. However, trend to lower long-term DE with increased visual perturbations. | Short-term DE reflects immediate response to perturbations. Long-term DE might not be sensitive to perturbations due to treadmill constraints. | Local dynamic stability during unperturbed walking does not predict response to perturbations in older adults, especially fallers. Optical flow perturbations reveal effects of aging and falls history not apparent during normal walking. | Optical flow perturbations could be used to identify subtle balance deficits not visible during normal walking. Understanding perturbation responses can guide the development of fall prevention strategies. |
| 48* | Investigate the sensitivity of short-term and long-term DEs to the noise structure of stride intervals. | Short-term DE not substantially affected by noise structure. Slight variations observed but not noise-type dependent. Long-term DE highly sensitive to noise structure, correlated with scaling exponents. | Short-term DE reflects local dynamic stability. Long-term DE likely indicates stride interval complexity Proposal to rename long-term DE as "Attractor Complexity Index" (ACI). | Short and long-term DE measure different aspects of gait. Long-term DE primarily reflects gait complexity, not stability | Long-term DE could reveal gait features related to automaticity or cautiousness. Potential use in assessing gait in clinical settings without step detection. |
| 49 | To evaluate the potential of trunk local dynamic stability (DEs) combined with clinical measures for fall risk prediction in elderly over one year. | Higher trunk mediolateral short-term DE in multiple fallers vs non-fallers. No significant differences in long-term DE between groups (all directions). Trend of lower DE in multiple fallers in vertical and AP directions. | Increased ML short-term DE reflects decreased local stability to perturbations during gait, linked to fall occurrence. | Trunk ML dynamic stability (short-term DE) combined with Tinetti score improves fall risk prediction, even in low-risk elderly. | DE from trunk accelerometry + clinical balance tests may enhance identification of high-risk elderly for targeted interventions. |
| 50 | Quantify gait variability differences between knee osteoarthritis (OA) patients and controls using nonlinear dynamics at 2 different walking speeds. | Short-term DE significantly higher in OA vs. controls. Lower long-term DE in OA vs. controls, but not significant. | Higher short-term DE in OA reflects reduced local stability. Long-term DE near zero in OA is interpreted as semi-periodic, more rigid signal. | OA gait is less stable, more rigid, and velocity changes minimally affect variability metrics. | Nonlinear measures like DEs can detect instability in OA gait, aiding fall risk assessment and therapy design. |
| 51* | Validate attractor complexity index (ACI=long-term DE) as a gait complexity measure from continuous signals; compare sensitivity to auditory/visual cueing vs. DFA. | Short-term DE unchanged except ML-direction auditory cueing (decrease) Long-term DE strongly decreased under cueing. | Short-term DE reflects gait stability; long-term DE (ACI) quantifies gait complexity/automaticity, linked to stride-interval noise structure (statistical persistence). | ACI correlates strongly with DFA and is equally predictive of cueing, enabling gait complexity assessment via continuous signals. | ACI enables gait complexity monitoring in clinical/real-world settings using inertial sensors (e.g., accelerometers), improving assessments of motor control. |
| 52 | Investigate gait stability and Preferred Walking Speed (PWS) in real and virtual environments (VR). Determine new proportionality coefficient for calculating PWS in VR. | Short-term DE: No significant differences across gait velocities in VR or real environments. Long-term DE: Significant differences | Short-term and long-term DEs indicate gait stability. | PWS is lower in VR than in real environments; a new proportionality coefficient is required for VR- | Accurate PWS determination in VR is critical for safe and effective rehabilitation using immersive technologies. |

COPYRIGHT 2026 THE AUTHORS. THIS FILE IS LICENSED UNDER A CREATIVE COMMONS ATTRIBUTION 4.0 INTERNATIONAL (CC BY 4.0) LICENSE.

| ID | Study aims | DE results | DE interpretation | Key conclusion | Clinical relevance |
|---|---|---|---|---|---|
| | | between speeds observed; lowest values at 0.9 PWS and 1.0 PWS, increasing with speed. | | based PWS calculations. | |
| 53 | Investigate impact of knee assistive device (KAD) on gait stability using nonlinear metrics and biomechanical measures, comparing control strategies (AIT: Automatic Impedance Tuning; FSM: Finite State Machine) | Lower short-term DE in AIT vs normal condition for left leg (unassisted), indicating improved local stability. Right leg (assisted) showed slight increase but no statistical significance.<br>Long-term DE decreased in AIT for left hip, knee, ankle; right hip/ankle as compared to normal condition. | Short-term DE correlates with fall probability. Positive exponents indicate local instability, with larger values suggesting greater sensitivity to perturbation. | Unilateral knee assistance triggers bilateral adaptation, with unassisted leg showing improved stability. AIT control strategy provides better stability than FSM through smooth impedance transitions between gait phase. | AIT's stability benefits support safer robot-assisted gait training. |
| 54* | Examine differential effects of head-worn smart glasses versus smartphone and paper-based displays on lateral stepping dynamics and stability during dual-task (DT) gait using gait variability measures from the GEM framework and local dynamic stability quantified by maximum Lyapunov exponent analysis. | Short-term DEs increased significantly in AP and V directions during DT walking with smartphone and paper-based systems compared to single-task and smart glasses.<br>Long-term DEs largely invariant across display conditions in most movement directions during DT walking. | Higher short-term DEs in DT walking indicates reduced stability, suggesting handheld displays impose greater destabilization. Non-significant long-term DE aligns with literature prioritizing short-term metrics for stability assessment. | Smart glasses caused less gait destabilization than paper/smartphone but uniquely disrupted lateral position control, reflecting a trade-off between stability and lateral stepping dynamics. | Smart glasses may reduce fall risk vs traditional displays but require design refinements to address lateral control deficits. Highlights need to monitor lateral position variability in dual-task gait assessments. |
| 55 | Investigate effects of mobile phone use on motor variability during gait, focusing on lower limb joint angles, muscle activation, and COM stability via short- and long-term DEs under six walking conditions. | Significant increases in short-term DEs for hip and pelvis sagittal angles during screen-focused tasks; no changes in ankle/knee/COM or muscle activation patterns.<br>Increased long-term DEs for hip and pelvis sagittal angles during screen use; other joints/COM/muscles showed no significant differences across conditions. | Higher hip/pelvis DEs indicates adaptive joint-angle variability to stabilize gait under visual/cognitive load; unchanged COM/muscle variability suggests compensatory neuromuscular control preserving whole-body stability. | Mobile phone use, particularly when involving visual distraction, selectively amplifies hip and pelvis joint variability without altering global dynamic stability, indicating that gait adjustments are task-specific compensatory responses rather than a state of general instability. | Screen viewing during walking may increase fall risk in complex environments despite maintained stability in controlled conditions. Long-term hip/pelvis adaptations could lead to excessive muscle activation and reduced gait economy |
| 56 | Investigate effects of load carriage and experience on gait variability (spatiotemporal, joint kinematic, Lyapunov exponents) among military cadets, assessing experience-related differences across three load conditions. | Hip short-term DE increased, knee short-term DE decreased in high load. No significant ankle short-term DE changes.<br>Lower ankle long-term DE in inexperienced cadets for heavy load. Hip/knee long-term DE showed no load/experience effects. | DEs quantify the divergence rate of gait trajectories, where higher short-term values indicate increased sensitivity to local perturbations and lower long-term stability reflects adaptive control. | Gait variability measures better reflect experience-related differences than traditional mean gait measures. Load and experience significantly affect movement strategies, with experienced cadets showing more controlled adaptations | Findings can guide military training programs and equipment design. Understanding experience-based movement strategies may help reduce injury risk during load carriage tasks |
| S1 | Compare the sensitivity of multiple dynamic balance measures — including short- and long-term DEs, gait variability, margin of stability, and whole-body angular momentum — for detecting differences between normal, dual-task (Stroop), and arm-restricted walking. | Short-term DE significantly higher during arm-restricted vs. normal walking for COM velocity,trunk velocity, hip joint angle, and also thigh, shank, foot velocities and knee joint angle. Short-term DE of hip joint angle was the only measure significantly higher during dual-task vs. normal walking. Long- | Higher short-term DE indicates greater local instability. The high sensitivity of trunk and hip measures reflects the dominance of trunk (~50%) and upper leg (~20%) body mass in dynamic balance control. | Short-term DE — particularly of COM velocity, trunk velocity, and hip joint angle — outperforms other balance measures in detecting gait instability under small continuous perturbations. Long-term DE showed no discriminatory power. | Provides guidance for selecting sensitive dynamic balance measures in clinical gait assessment; short-term DE of trunk and hip signals is recommended over traditional gait variability or biomechanical measures. |

COPYRIGHT 2026 THE AUTHORS. THIS FILE IS LICENSED UNDER A CREATIVE COMMONS ATTRIBUTION 4.0 INTERNATIONAL (CC BY 4.0) LICENSE. 

| ID | Study aims | DE results | DE interpretation | Key conclusion | Clinical relevance |
|---|---|---|---|---|---|
| | | term DE showed no significant differences across any condition or signal. | | | |
| S2* | Validate the attractor complexity index (ACI = long-term DE) as an indicator of gait automaticity and cautiousness in older adults. Compare responsiveness to metronome walking of short-term DE (LDS), long-term DE (ACI), and scaling exponent (DFA). Assess intrasession reliability of ACI versus DFA. | Short-term DE unchanged during metronome walking (ES close to zero across all axes). Long-term DE (ACI) significantly decreased during metronome walking. Significant correlations between ACI and DFA scaling exponents. ACI showed moderate intrasession reliability superior to DFA scaling exponent. | Short-term DE reflects gait stability and resilience to perturbations, unaffected by metronome synchronization. Long-term DE (ACI) captures the correlation structure of stride-to-stride variability, indicating the level of voluntary gait control and cautiousness. ACI and DFA scaling exponents reflect analogous dimensions of gait dynamics. | ACI is validated as a viable and more reliable alternative to DFA for assessing gait automaticity in older adults. ACI requires fewer consecutive strides than DFA and can be computed from a single lumbar accelerometer signal without step detection. | ACI enables continuous, unobtrusive gait quality monitoring in older adults using a single lumbar accelerometer, potentially improving early fall risk identification in unsupervised real-world settings. |
| S3 | Compare three methods for computing DEs and Floquet multipliers (FM) from short walking trials: Method A (GFNN + Rosenstein's numerical approximation), Method B (GFNN + semi-analytical technique), and Method C (SVD + semi-analytical technique). Determine which approach best matches observed gait behavior. | Short-term DE (Method A) indicating chaotic gait for all subjects. Long-term DE , chaos for ~half of subjects; Method C non-chaotic for most subjects. Long-term DE from Methods A and B were statistically similar; all other DE pairings differed significantly. | Short-term DE from Rosenstein's algorithm overestimates chaos because of numerical approximation errors. Positive DE values in healthy non-fallers are considered artefactual. | The SVD and semi-analytical approach (Method C) yielded DE and FM values most closely matching observed gait patterns. Rosenstein's algorithm introduces approximation errors, particularly for short-term DE. SVD-based dimensionality reduction enables reliable dynamical systems analysis from short walking trials. | Provides a methodological framework for applying dynamical systems theory to short walking trials, which is important for patients unable to walk for extended durations. SVD-based noise reduction may improve the reliability of DE estimation in clinical gait assessments. |
| S4* | Assess ACI's (long-term DE) sensitivity to attentional demands during gait control and potential for characterizing age-related gait changes. Compare ACI with classical gait metrics (movement intensity, regularity, short-term DE). Test two hypotheses: (1) unique sensitivity of ACI to metronome walking compared to other lumbar accelerometer metrics; (2) ACI discriminates age-related gait pattern changes as effectively as other gait metrics. | Short-term DE (LDS-ML): small but not significant age difference. Metronome walking did not affect short-term DE. Long-term DE (ACI): Significant age differences. Both groups showed substantial decreases during metronome walking. ACI-AP demonstrated strongest age discrimination and was speed-independent, while ACI-V and ACI-N correlated with walking speed. | Short-term DE reflects local dynamic stability and resilience to perturbations. Long-term DE (ACI) captures stride-to-stride correlation structure, shifting from persistent (fractal) patterns during normal walking to anti-persistent patterns during metronome walking. This shift indicates altered gait automaticity and increased attentional control. Lower ACI values in older adults suggest reduced gait complexity and automaticity compared to younger adults. | Long-term DE demonstrates specific sensitivity to metronome walking, distinct from other gait metrics. Long-term DE effectively discriminates age-related gait changes performing comparably to step/stride regularity measures. Long-term DE provides complementary information to traditional variability metrics, potentially revealing aspects of gait decline unrelated to walking speed reduction. | Long-term DE ease of measurement from a single lumbar accelerometer makes it promising for continuous gait quality monitoring in unsupervised (free-living) conditions. Could serve as sensitive marker for identifying older adults at fall risk through assessment of gait automaticity. May help evaluate interventions targeting gait automaticity restoration. |
| S5 | Investigate effects of 7 outdoor surfaces on nonlinear gait dynamics in healthy adults. Examine sex differences irrespective of surface. Investigate interaction effects of surface and sex on movement predictability (sample entropy), smoothness, symmetry (step/stride regularity), and | Short-term DEs: Cobblestone significantly decreased short-term DE. Sloped-up increased short-term DE. Other surfaces showed no significant changes. Long-term DE: All surfaces significantly increased long-term DE compared to flat-even No significant sex main | Short-term LyE reflects local dynamic stability and step-to-step variability, with higher values indicating lower stability and greater sensitivity to perturbations. Long-term LyE quantifies long-term divergence and complexity of gait attractor dynamics. | Surface type significantly influenced both short- and long-term DEs independent of sex. Despite successful navigation without falls, all challenging surfaces prompted deteriorations in movement | Even in healthy young adults successfully navigating familiar outdoor terrains, surface characteristics substantially alter gait dynamics. Findings emphasize need for outdoor gait assessment in clinical populations. |

COPYRIGHT 2026 THE AUTHORS. THIS FILE IS LICENSED UNDER A CREATIVE COMMONS ATTRIBUTION 4.0 INTERNATIONAL (CC BY 4.0) LICENSE.

| ID | Study aims | DE results | DE interpretation | Key conclusion | Clinical relevance |
|---|---|---|---|---|---|
| | stability (short-/long-term DEs). | effects, though males showed non-significant 13.71% decrease in DEs. No significant surface-sex interactions for DEs. | | smoothness, altered symmetry, and increased stride-to-stride fluctuations. | |
| S6 | Identify the most effective combination of data modality (trunk acceleration & joint kinematics) and nonlinear dynamic metric (DE and multiscale entropy) for distinguishing fallers from non-fallers in older women across three treadmill speed conditions (preferred, slow, fast). | Short-term DE of ankle sagittal-plane angle significantly higher in fallers at slow and fast speeds. Short-term DE of trunk AP acceleration significantly higher in fallers at preferred and slow speeds. CoM vertical DE showed no significant group difference. ROC analysis: ankle sagittal DE achieved best discrimination, trunk AP DE fair discrimination (AUC up to 0.77), CoM vertical DE poor discrimination (AUC = 0.53). Age and self-selected speed did not confound the results. | Higher short-term DE in fallers reflects reduced local dynamic stability and greater sensitivity to perturbations at both distal (ankle) and central (trunk) segments. | Short-term DE from ankle sagittal motion and trunk AP acceleration are the most effective nonlinear dynamic measures for discriminating fall history in older women. Results were consistent across Rosenstein's and Wolf's algorithms and across IMU and motion capture modalities. Long-term DE and MSE contributed to PCA variance but did not emerge as primary discriminators in logistic regression. | Comparable performance of IMU-based and motion capture-based short-term DE supports the feasibility of scalable, wearable fall risk screening using a single lumbar sensor. Short-term DE of ankle and trunk segments could enhance objective fall risk identification in community or clinical settings, complementing existing clinical assessments. |


COPYRIGHT 2026 THE AUTHORS. THIS FILE IS LICENSED UNDER A CREATIVE COMMONS ATTRIBUTION 4.0 INTERNATIONAL (CC BY 4.0) LICENSE. 

[10] K. Jordan, J.H. Challis, J.P. Cusumano, K.M. Newell, Stability and the time-dependent structure of gait variability in walking and running, Human Movement Science 28 (2009) 113–128. https://doi.org/10.1016/j.humov.2008.09.001.
[11] S.M. Bruijn, J.H. Van Dieën, O.G. Meijer, P.J. Beek, Statistical precision and sensitivity of measures of dynamic gait stability, Journal of Neuroscience Methods 178 (2009) 327–333. https://doi.org/10.1016/j.jneumeth.2008.12.015.
[12] S.M. Bruijn, J.H. Van Dieën, O.G. Meijer, P.J. Beek, Is slow walking more stable?, Journal of Biomechanics 42 (2009) 1506–1512. https://doi.org/10.1016/j.jbiomech.2009.03.047.
[13] H.G. Kang, J.B. Dingwell, Dynamic stability of superior vs. inferior segments during walking in young and older adults, Gait & Posture 30 (2009) 260–263. https://doi.org/10.1016/j.gaitpost.2009.05.003.
[14] H.G. Kang, J.B. Dingwell, Dynamics and stability of muscle activations during walking in healthy young and older adults, Journal of Biomechanics 42 (2009) 2231–2237. https://doi.org/10.1016/j.jbiomech.2009.06.038.
[15] B. Manor, P. Wolenski, A. Guevaro, L. Li, Differential effects of plantar desensitization on locomotion dynamics, Journal of Electromyography and Kinesiology 19 (2009) e320–e328. https://doi.org/10.1016/j.jelekin.2008.06.006.
[16] J.A. Nessler, C.J. De Leone, S. Gilliland, Nonlinear time series analysis of knee and ankle kinematics during side by side treadmill walking, Chaos: An Interdisciplinary Journal of Nonlinear Science 19 (2009) 026104. https://doi.org/10.1063/1.3125762.
[17] K. Son, J. Park, S. Park, Variability analysis of lower extremity joint kinematics during walking in healthy young adults, Medical Engineering & Physics 31 (2009) 784–792. https://doi.org/10.1016/j.medengphy.2009.02.009.
[18] S.M. Bruijn, O.G. Meijer, P.J. Beek, J.H. Van Dieën, The effects of arm swing on human gait stability, Journal of Experimental Biology 213 (2010) 3945–3952. https://doi.org/10.1242/jeb.045112.
[19] S.M. Bruijn, W.R.Th. Ten Kate, G.S. Faber, O.G. Meijer, P.J. Beek, J.H. Van Dieën, Estimating Dynamic Gait Stability Using Data from Non-aligned Inertial Sensors, Ann Biomed Eng 38 (2010) 2588–2593. https://doi.org/10.1007/s10439-010-0018-2.
[20] M.D. Chang, E. Sejdić, V. Wright, T. Chau, Measures of dynamic stability: Detecting differences between walking overground and on a compliant surface, Human Movement Science 29 (2010) 977–986. https://doi.org/10.1016/j.humov.2010.04.009.
[21] H.R. Yakhdani, H.A. Bafghi, O.G. Meijer, S.M. Bruijn, N.V.D. Dikkenberg, A.B. Stibbe, B.J. Van Royen, J.H. Van Dieën, Stability and variability of knee kinematics during gait in knee osteoarthritis before and after replacement surgery, Clinical Biomechanics 25 (2010) 230–236. https://doi.org/10.1016/j.clinbiomech.2009.12.003.
[22] P.M. McAndrew, J.M. Wilken, J.B. Dingwell, Dynamic stability of human walking in visually and mechanically destabilizing environments, Journal of Biomechanics 44 (2011) 644–649. https://doi.org/10.1016/j.jbiomech.2010.11.007.
[23] J.A. Nessler, H. Huynh, M. MCDougal, A single bout of resistance exercise does not affect nonlinear dynamics of lower extremity kinematics during treadmill walking, Gait & Posture 34 (2011) 285–287. https://doi.org/10.1016/j.gaitpost.2011.04.003.
[24] K. Son, J. Park, S. Park, Kinematic analysis of the neck and upper extremities during walking in healthy young adults, J Bionic Eng 8 (2011) 305–312. https://doi.org/10.1016/S1672-6529(11)60025-5.
[25] P. Terrier, O. Dériaz, Kinematic variability, fractal dynamics and local dynamic stability of treadmill walking, J NeuroEngineering Rehabil 8 (2011) 12. https://doi.org/10.1186/1743-0003-8-12.
[26] K.S. Van Schooten, L.H. Sloot, S.M. Bruijn, H. Kingma, O.G. Meijer, M. Pijnappels, J.H. Van Dieën, Sensitivity of trunk variability and stability measures to balance

COPYRIGHT 2026 THE AUTHORS. THIS FILE IS LICENSED UNDER A CREATIVE COMMONS ATTRIBUTION 4.0 INTERNATIONAL (CC BY 4.0) LICENSE. 

impairments induced by galvanic vestibular stimulation during gait, Gait & Posture 33 (2011) 656–660. https://doi.org/10.1016/j.gaitpost.2011.02.017.
[27] P.M. McAndrew Young, J.B. Dingwell, Voluntarily changing step length or step width affects dynamic stability of human walking, Gait Posture 35 (2012) 472–477. https://doi.org/10.1016/j.gaitpost.2011.11.010.
[28] F. Cignetti, L.M. Decker, N. Stergiou, Sensitivity of the Wolf's and Rosenstein's Algorithms to Evaluate Local Dynamic Stability from Small Gait Data Sets, Ann Biomed Eng 40 (2012) 1122–1130. https://doi.org/10.1007/s10439-011-0474-3.
[29] E. Sejdić, Y. Fu, A. Pak, J.A. Fairley, T. Chau, The Effects of Rhythmic Sensory Cues on the Temporal Dynamics of Human Gait, PLoS ONE 7 (2012) e43104. https://doi.org/10.1371/journal.pone.0043104.
[30] E.H. Sinitksi, K. Terry, J.M. Wilken, J.B. Dingwell, Effects of perturbation magnitude on dynamic stability when walking in destabilizing environments, Journal of Biomechanics 45 (2012) 2084–2091. https://doi.org/10.1016/j.jbiomech.2012.05.039.
[31] M.J.P. Toebes, M.J.M. Hoozemans, R. Furrer, J. Dekker, J.H. Van Dieën, Local dynamic stability and variability of gait are associated with fall history in elderly subjects, Gait & Posture 36 (2012) 527–531. https://doi.org/10.1016/j.gaitpost.2012.05.016.
[32] J. Liu, T.E. Lockhart, Local Dynamic Stability Associated with Load Carrying, Safety and Health at Work 4 (2013) 46–51. https://doi.org/10.5491/SHAW.2013.4.1.46.
[33] J.A. Nessler, D. McMillan, M. Schoulten, T. Shallow, B. Stewart, C. De Leone, Side by Side Treadmill Walking With Intentionally Desynchronized Gait, Ann Biomed Eng 41 (2013) 1680–1691. https://doi.org/10.1007/s10439-012-0657-6.
[34] E. Sejdić, B. Findlay, C. Merey, T. Chau, The effects of listening to music or viewing television on human gait, Computers in Biology and Medicine 43 (2013) 1497–1501. https://doi.org/10.1016/j.compbiomed.2013.07.019.
[35] P. Terrier, O. Dériaz, Non-linear dynamics of human locomotion: effects of rhythmic auditory cueing on local dynamic stability, Front. Physiol. 4 (2013). https://doi.org/10.3389/fphys.2013.00230.
[36] F. Reynard, P. Terrier, Local dynamic stability of treadmill walking: Intrasession and week-to-week repeatability, Journal of Biomechanics 47 (2014) 74–80. https://doi.org/10.1016/j.jbiomech.2013.10.011.
[37] F. Riva, M.C. Bisi, R. Stagni, Gait variability and stability measures: Minimum number of strides and within-session reliability, Computers in Biology and Medicine 50 (2014) 9–13. https://doi.org/10.1016/j.compbiomed.2014.04.001.
[38] F. Riva, E. Grimpampi, C. Mazzà, R. Stagni, Are gait variability and stability measures influenced by directional changes?, BioMed Eng OnLine 13 (2014) 56. https://doi.org/10.1186/1475-925X-13-56.
[39] F. Yang, Y.-C. Pai, Can stability really predict an impending slip-related fall among older adults?, Journal of Biomechanics 47 (2014) 3876–3881. https://doi.org/10.1016/j.jbiomech.2014.10.006.
[40] S.M. Beaudette, T.A. Worden, M. Kamphuis, L. Ann Vallis, S.H.M. Brown, Local Dynamic Joint Stability During Human Treadmill Walking in Response to Lower Limb Segmental Loading Perturbations, Journal of Biomechanical Engineering 137 (2015) 091006. https://doi.org/10.1115/1.4030944.
[41] J.R. Franz, C.A. Francis, M.S. Allen, S.M. O'Connor, D.G. Thelen, Advanced age brings a greater reliance on visual feedback to maintain balance during walking, Human Movement Science 40 (2015) 381–392. https://doi.org/10.1016/j.humov.2015.01.012.
[42] C.A. Rábago, J.B. Dingwell, J.M. Wilken, Reliability and Minimum Detectable Change of Temporal-Spatial, Kinematic, and Dynamic Stability Measures during Perturbed Gait, PLoS ONE 10 (2015) e0142083. https://doi.org/10.1371/journal.pone.0142083.

COPYRIGHT 2026 THE AUTHORS. THIS FILE IS LICENSED UNDER A CREATIVE COMMONS ATTRIBUTION 4.0 INTERNATIONAL (CC BY 4.0) LICENSE. 

#### References (Update screening)

[S1] S. Yamagata, T. Yamaguchi, M. Shinya, M. Milosevic, K. Masani, Comparison of sensitivity among dynamic balance measures during walking with different tasks, ROYAL SOCIETY OPEN SCIENCE. 11 (2024). https://doi.org/10.1098/rsos.230883.

COPYRIGHT 2026 THE AUTHORS. THIS FILE IS LICENSED UNDER A CREATIVE COMMONS ATTRIBUTION 4.0 INTERNATIONAL (CC BY 4.0) LICENSE. 

[S2] S. Piergiovanni, P. Terrier, Effects of metronome walking on long-term attractor divergence and correlation structure of gait: a validation study in older people, SCIENTIFIC REPORTS. 14 (2024). https://doi.org/10.1038/s41598-024-65662-5.
[S3] S.G. Bhat, K.R. Kaufman, Dynamical systems theory applied to short walking trials, JOURNAL OF BIOMECHANICS. 176 (2024). https://doi.org/10.1016/j.jbiomech.2024.112331.
[S4] S. Piergiovanni, P. Terrier, Validity of Linear and Nonlinear Measures of Gait Variability to Characterize Aging Gait with a Single Lower Back Accelerometer, SENSORS. 24 (2024). https://doi.org/10.3390/s24237427.
[S5] J. Emmerzaal, P. Ippersiel, P.C. Dixon, Non-Linear Gait Dynamics Are Affected by Commonly Occurring Outdoor Surfaces and Sex in Healthy Adults, SENSORS. 25 (2025). https://doi.org/10.3390/s25134191.
[S6] A. Amirpourabasi, S.E. Lamb, J.Y. Chow, G.K.R. Williams, Using nonlinear dynamic analysis to differentiate fall status in older women, GAIT & POSTURE. 124 (2026). https://doi.org/10.1016/j.gaitpost.2025.110032.

COPYRIGHT 2026 THE AUTHORS. THIS FILE IS LICENSED UNDER A CREATIVE COMMONS ATTRIBUTION 4.0 INTERNATIONAL (CC BY 4.0) LICENSE.
