## Supplemental file 2 (divergence curves) for "From Stability to Complexity: A Systematic Review of Long-term Divergence Exponents in Nonlinear Gait Analysis"

¶ Equal contribution

This supplementary document compiles divergence curve figures from the 30 studies (48% of the 62 included studies) that published graphical representations of logarithmic divergence curves in their manuscripts. Study IDs correspond to those listed in S1 Text. Divergence curves plot the average logarithmic divergence of neighboring trajectories against time, demonstrating how short-term (0-1 stride) and long-term (4-10 strides) divergence exponents are extracted as linear slopes from distinct temporal regions. These visual representations serve two functions: they illustrate the computational methodology for calculating divergence exponents using Rosenstein's algorithm, and their shape reveals when divergence ceases to grow as trajectories reach bounded regions of phase space. We systematically categorized each study's divergence curve presentation by purpose: methodological illustration (demonstrating the calculation procedure without comparing conditions), comparative results (highlighting differences between experimental groups or conditions), or both.

The document is organized in two parts. The first part lists studies published behind paywalls, for which only the original figure legends and category classifications are provided. The second part presents studies published under open-access licenses, for which the divergence curve figures have been reproduced along with their complete legends. Within each part, entries are ordered by Study ID, corresponding to study IDs of S1_Text.

COPYRIGHT 2025 THE AUTHORS. THIS FILE IS LICENSED UNDER A CREATIVE COMMONS ATTRIBUTION 4.0 INTERNATIONAL (CC BY 4.0) LICENSE. 

**ID 1.** J.B. Dingwell, J.P. Cusumano, D. Sternad, P.R. Cavanagh, Slower speeds in patients with diabetic neuropathy lead to improved local dynamic stability of continuous overground walking, Journal of Biomechanics 33 (2000) 1269–1277. https://doi.org/10.1016/S0021-9290(00)00092-0.

"Fig. 2. Schematic representation of local dynamic stability analysis."

Category: Methodological illustration

**ID 2.** J.B. Dingwell, J.P. Cusumano, Nonlinear time series analysis of normal and pathological human walking, Chaos: An Interdisciplinary Journal of Nonlinear Science 10 (2000) 848–863. https://doi.org/10.1063/1.1324008.

"FIG. 7. Logarithmic divergence curves for a typical subject for both OG and TM walking."

Category: Methodological illustration & Comparative results

**ID 3.** J.B. Dingwell, J.P. Cusumano, P.R. Cavanagh, D. Sternad, Local Dynamic Stability Versus Kinematic Variability of Continuous Overground and Treadmill Walking, Journal of Biomechanical Engineering 123 (2001) 27–32. https://doi.org/10.1115/1.1336798.

"Fig. 2 Schematic representation of local stability analysis."
"Fig. 5 Representative plots of the average logarithmic divergence,"

Category: Methodological illustration & Comparative results

**ID 5.** J.B. Dingwell, L.C. Marin, Kinematic variability and local dynamic stability of upper body motions when walking at different speeds, Journal of Biomechanics 39 (2006) 444–452. https://doi.org/10.1016/j.jbiomech.2004.12.014.

"Fig. 3. Schematic representation of local dynamic stability analysis."
"Fig. 5. […] (B) Logarithmic divergence curves obtained from dE ¼ 5 state spaces"

Category: Methodological illustration

**ID 6.** H.G. Kang, J.B. Dingwell, Intra-session reliability of local dynamic stability of walking, Gait & Posture 24 (2006) 386–390. https://doi.org/10.1016/j.gaitpost.2005.11.004.

"Fig. 1. Schematic representation of state space construction and local dynamic stability analysis for a single trial."
"Fig. 2. Mean linear divergence curves from a typical subject."

Category: Methodological illustration & Comparative results

COPYRIGHT 2025 THE AUTHORS. THIS FILE IS LICENSED UNDER A CREATIVE COMMONS ATTRIBUTION 4.0 INTERNATIONAL (CC BY 4.0) LICENSE.

**ID 7.** J.B. Dingwell, H.G. Kang, Differences Between Local and Orbital Dynamic Stability During Human Walking, Journal of Biomechanical Engineering 129 (2007) 586–593. https://doi.org/10.1115/1.2746383.

Fig 3C. Schematic representation, legend is missing for divergence subplot.

Category: Methodological illustration

**ID 8.** H.G. Kang, J.B. Dingwell, Effects of walking speed, strength and range of motion on gait stability in healthy older adults, Journal of Biomechanics 41 (2008) 2899–2905. https://doi.org/10.1016/j.jbiomech.2008.08.002.

"Fig. 1. Schematic representation of state-space construction."
"Fig. 3. (A) Sample local divergence curves"

Category: Methodological illustration & Comparative results

**ID 9.** B. Manor, P. Wolenski, L. Li, Faster walking speeds increase local instability among people with peripheral neuropathy, Journal of Biomechanics 41 (2008) 2787–2792. https://doi.org/10.1016/j.jbiomech.2008.07.006.

"Fig. 2. Representation of state-space reconstruction and divergence analysis."

Category: Methodological illustration

**ID 10.** K. Jordan, J.H. Challis, J.P. Cusumano, K.M. Newell, Stability and the time-dependent structure of gait variability in walking and running, Human Movement Science 28 (2009) 113–128. https://doi.org/10.1016/j.humov.2008.09.001.

"Fig. 1. The average logarithmic divergence of all pairs of neighboring trajectories in state space over time for a representative trial."

Category: Methodological illustration

**ID 12.** S.M. Bruijn, J.H. Van Dieën, O.G. Meijer, P.J. Beek, Is slow walking more stable?, Journal of Biomechanics 42 (2009) 1506–1512. https://doi.org/10.1016/j.jbiomech.2009.03.047.

"Fig. 1. Schematic representation of the calculation of maximum time finite Lyapunov exponents."

Category: Methodological illustration

**ID 13.** H.G. Kang, J.B. Dingwell, Dynamic stability of superior vs. inferior segments during walking in young and older adults, Gait & Posture 30 (2009) 260–263. https://doi.org/10.1016/j.gaitpost.2009.05.003.

COPYRIGHT 2025 THE AUTHORS. THIS FILE IS LICENSED UNDER A CREATIVE COMMONS ATTRIBUTION 4.0 INTERNATIONAL (CC BY 4.0) LICENSE. 

"Fig. 1. Schematic representation of state space construction. […] Rates of divergence, […] (local divergence exponents), were calculated from the slopes of the mean log divergence curve."

Category: Methodological illustration

**ID 14.** H.G. Kang, J.B. Dingwell, Dynamics and stability of muscle activations during walking in healthy young and older adults, Journal of Biomechanics 42 (2009) 2231–2237. https://doi.org/10.1016/j.jbiomech.2009.06.038.

"Fig. 2. Schematic representation of state space embedding and dynamic stability analyses of EMG signals."

Category: Methodological illustration

**ID 15.** B. Manor, P. Wolenski, A. Guevaro, L. Li, Differential effects of plantar desensitization on locomotion dynamics, Journal of Electromyography and Kinesiology 19 (2009) e320–e328. https://doi.org/10.1016/j.jelekin.2008.06.006.

"Fig. 2. Representation of two-dimensional state space reconstruction and divergence analysis."

Category: Methodological illustration

**ID 16.** J.A. Nessler, C.J. De Leone, S. Gilliland, Nonlinear time series analysis of knee and ankle kinematics during side by side treadmill walking, Chaos: An Interdisciplinary Journal of Nonlinear Science 19 (2009) 026104. https://doi.org/10.1063/1.3125762.

"FIG. 2. Graphical description of the procedure for estimating short term and long term maximal Lyapunov exponents."
"FIG. 5. Mean divergence curves for the solo walking condition, paired condition, forced condition, and surrogate data…"
"FIG. 6. Sample data for an individual who demonstrated signatures of chaotic behavior during treadmill walking for ankle $Y$ data."

Category: Methodological illustration & Comparative results

**ID 17.** K. Son, J. Park, S. Park, Variability analysis of lower extremity joint kinematics during walking in healthy young adults, Medical Engineering & Physics 31 (2009) 784–792. https://doi.org/10.1016/j.medengphy.2009.02.009.

"Fig. 8. LE estimates of a typical subject M1 calculated by linear slopes at intervals of 4–10 s..."

Category: Methodological illustration

COPYRIGHT 2025 THE AUTHORS. THIS FILE IS LICENSED UNDER A CREATIVE COMMONS ATTRIBUTION 4.0 INTERNATIONAL (CC BY 4.0) LICENSE.

**ID 21.** H.R. Yakhdani, H.A. Bafghi, O.G. Meijer, S.M. Bruijn, N.V.D. Dikkenberg, A.B. Stibbe, B.J. Van Royen, J.H. Van Dieën, Stability and variability of knee kinematics during gait in knee osteoarthritis before and after replacement surgery, Clinical Biomechanics 25 (2010) 230–236. https://doi.org/10.1016/j.clinbiomech.2009.12.003.

"Fig. 1. The natural logarithm of divergence as it developed over time in a representative subject,"

Category: Methodological illustration

**ID 22.** P.M. McAndrew, J.M. Wilken, J.B. Dingwell, Dynamic stability of human walking in visually and mechanically destabilizing environments, Journal of Biomechanics 44 (2011) 644–649. https://doi.org/10.1016/j.jbiomech.2010.11.007.

"Fig. 2. Average mean log divergence (MLD) curves for ML movements of the C7 marker for all perturbation conditions."

Category: Comparative results

**ID 24.** K. Son, J. Park, S. Park, Kinematic analysis of the neck and upper extremities during walking in healthy young adults, J Bionic Eng 8 (2011) 305–312. https://doi.org/10.1016/S1672-6529(11)60025-5.

"Fig. 6 LE values of a typical male subject, as calculated by the linear slopes from the plots of y(i) vs. i…"

Category: Methodological illustration

**ID 28.** F. Cignetti, L.M. Decker, N. Stergiou, Sensitivity of the Wolf's and Rosenstein's Algorithms to Evaluate Local Dynamic Stability from Small Gait Data Sets, Ann Biomed Eng 40 (2012) 1122–1130. https://doi.org/10.1007/s10439-011-0474-3.

"FIGURE 2. Illustration of the attractor reconstruction using the time delay method. […] Rate of divergence, k1 (largest Lyapunov exponent), was calculated with the R-algorithm from the slope of the mean log divergence curve between 0–1 stride and 4–10 strides."

Category: Methodological illustration

**ID 40.** S.M. Beaudette, T.A. Worden, M. Kamphuis, L. Ann Vallis, S.H.M. Brown, Local Dynamic Joint Stability During Human Treadmill Walking in Response to Lower Limb Segmental Loading Perturbations, Journal of Biomechanical Engineering 137 (2015) 091006. https://doi.org/10.1115/1.4030944.

"Fig. 1 Schematic depiction of the angular joint time series data processing steps used in the estimation of lumbar spine, hip (shown), knee, and ankle LDS. […] (d) creation of an average maximal logarithmic divergence ({lndj(i)}) curve."

Category: Methodological illustration

COPYRIGHT 2025 THE AUTHORS. THIS FILE IS LICENSED UNDER A CREATIVE COMMONS ATTRIBUTION 4.0 INTERNATIONAL (CC BY 4.0) LICENSE. 

**ID 41.** J.R. Franz, C.A. Francis, M.S. Allen, S.M. O'Connor, D.G. Thelen, Advanced age brings a greater reliance on visual feedback to maintain balance during walking, Human Movement Science 40 (2015) 381–392. https://doi.org/10.1016/j.humov.2015.01.012.

"Fig. 5. Local dynamic stability results. (A) Mean (standard error) divergence curves […] for old and young adults during normal (black line) and visually perturbed (blue line) walking."

Category: Comparative results

**ID 50.** N. Matinazad, A. Esteki, H. Ghomashchi, Comprehensive characterization of gait variability in patients with knee osteoarthritis for altered velocities, J. Mech. Med. Biol. 18 (2018) 1850041. https://doi.org/10.1142/S0219519418500410.

"Fig. 1. The utilized nonlinear tools, […] the LyE graph represents the average logarithmic rate of divergence,"

Category: Methodological illustration

COPYRIGHT 2025 THE AUTHORS. THIS FILE IS LICENSED UNDER A CREATIVE COMMONS ATTRIBUTION 4.0 INTERNATIONAL (CC BY 4.0) LICENSE.

**ID 25.** P. Terrier, O. Dériaz, Kinematic variability, fractal dynamics and local dynamic stability of treadmill walking, J NeuroEngineering Rehabil 8 (2011) 12. https://doi.org/10.1186/1743-0003-8-12.

Category: Methodological illustration

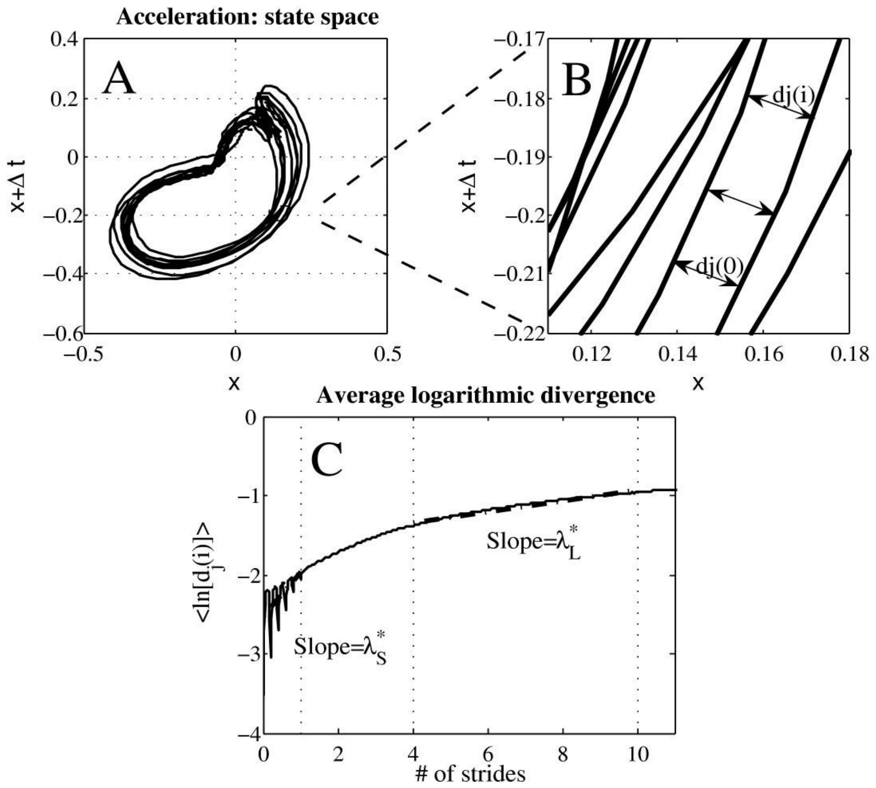


**Figure 3. Method: dynamic stability, maximal Lyapunov exponent**. **A:** Two dimensional state space of the antero-posterior acceleration signal (5s) reconstructed from the original data set and its time delayed copy (Δt = 11 samples). **B:** Magnification of the state space. An initial local perturbation at dj(0) diverge across i time steps as measured by dj(i). **C:** Short term ($\lambda_S$*) and long term ($\lambda_L$*) finite-time maximal Lyapunov exponent computed from average logarithmic divergence.

COPYRIGHT 2025 THE AUTHORS. THIS FILE IS LICENSED UNDER A CREATIVE COMMONS ATTRIBUTION 4.0 INTERNATIONAL (CC BY 4.0) LICENSE. 

**ID 32.** J. Liu, T.E. Lockhart, Local Dynamic Stability Associated with Load Carrying, Safety and Health at Work 4 (2013) 46–51. https://doi.org/10.5491/SHAW.2013.4.1.46.

Category: Comparative results

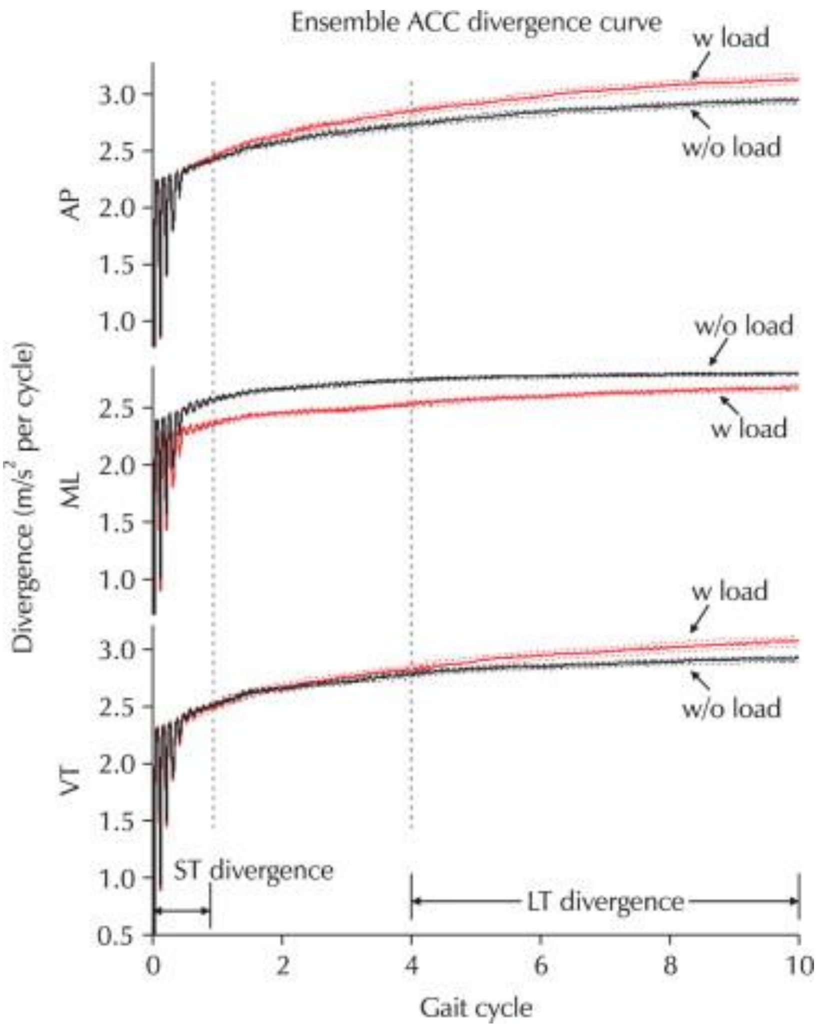


Fig. 2. Ensemble divergence curve of state space reconstructed from acceleration signals for each axis. ST and LT maxLE were calculated for ST and LT divergence ranges, respectively. The dashed line around the ensemble average curve indicated ± 3 standard error. ACC: accelerometer, w: with, w/o: without, AP: anterio-posterior, ML: mediolateral, VT: vertical, ST: short-term, LT: long-term.

COPYRIGHT 2025 THE AUTHORS. THIS FILE IS LICENSED UNDER A CREATIVE COMMONS ATTRIBUTION 4.0 INTERNATIONAL (CC BY 4.0) LICENSE. 

**ID 35** P. Terrier, O. Dériaz, Non-linear dynamics of human locomotion: effects of rhythmic auditory cueing on local dynamic stability, Front. Physiol. 4 (2013). https://doi.org/10.3389/fphys.2013.00230.

Category: Comparative results

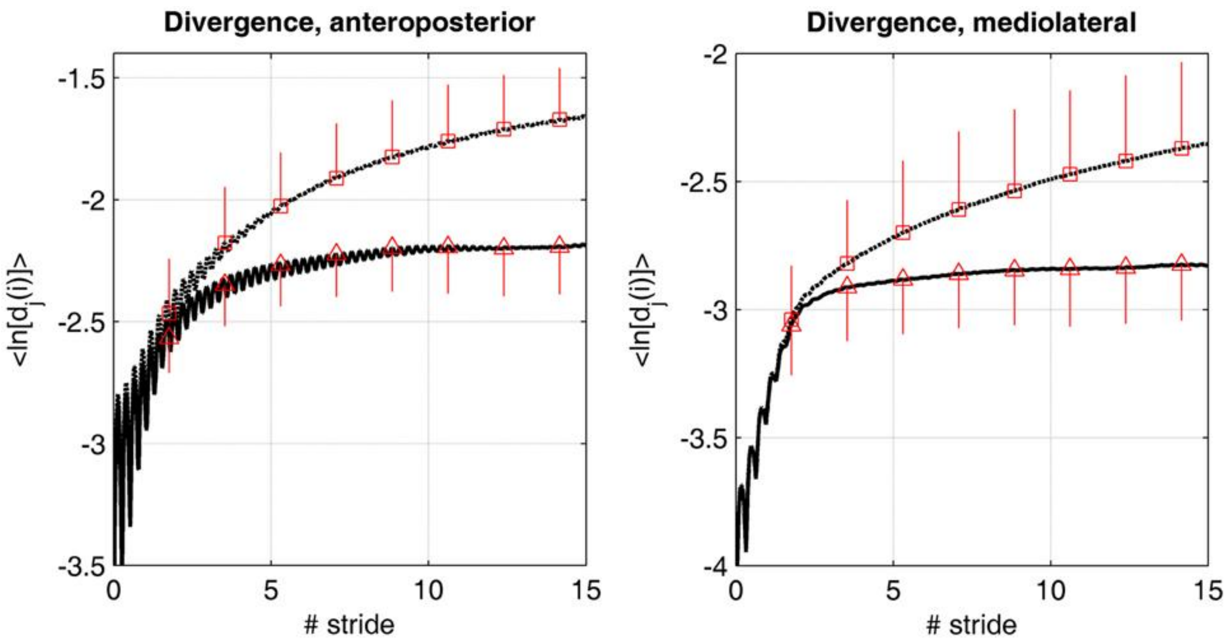


**Figure 2. Divergence curves.** The average logarithmic divergence (<ln[$d_j$(i)]>) in anteroposterior and mediolateral directions was measured in the reconstructed state space of the center of pressure trajectory (50 Hz sampling rate), in 20 individuals walking at preferred walking speed. 175 consecutive strides were analyzed, normalized at 10,000 samples. The value at each time (50 Hz) was averaged across the subjects (*N* = 20). Time was normalized by the average stride time (1.14 s). Discontinuous lines (squares) are the results for the treadmill only condition. Continuous lines (triangles) are the results for the dual cueing condition (treadmill + rhythmic auditory cueing). Mean value at 100, 200, 300, 400, 500, 600, 700, and 800 samples are shown (squares and triangles) with the corresponding SD (vertical lines, *N* = 20).

COPYRIGHT 2025 THE AUTHORS. THIS FILE IS LICENSED UNDER A CREATIVE COMMONS ATTRIBUTION 4.0 INTERNATIONAL (CC BY 4.0) LICENSE. 

**ID 48.** P. Terrier, F. Reynard, Maximum Lyapunov exponent revisited: Long-term attractor divergence of gait dynamics is highly sensitive to the noise structure of stride intervals, Gait & Posture 66 (2018) 236–241. https://doi.org/10.1016/j.gaitpost.2018.08.010.

Category: Comparative results

Note: An open-access preprint of this study is available (arXiv:1802.03223, https://doi.org/10.48550/arXiv.1802.03223). The figure below was extracted from this preprint, which is not subject to the publisher's copyright restrictions on the final published version.

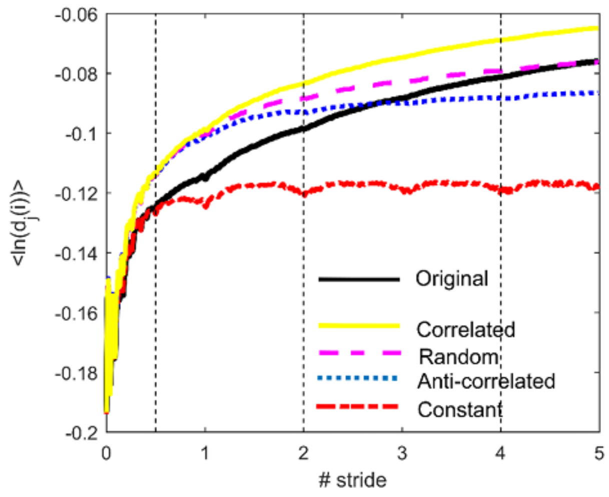


**Fig 2.** Divergence curves. A 5-dimensional attractor was built through the delay-embedding theorem from the trunk acceleration signal. Divergence between neighboring trajectories in the attractor is shown. For each noise type, 109 curves from 69 individuals were aggregated. Original stride intervals were changed to four different noise structures (hybrid signals). X-axis = time $i$ normalized by stride. Y-axis = the logarithm of the $i^{th}$ Euclidian distance $d$ downstream of the $j^{th}$ pair of the nearest neighbors in the attractor, averaged over all the pairs: $<\ln[d_j(i)]>$.

COPYRIGHT 2025 THE AUTHORS. THIS FILE IS LICENSED UNDER A CREATIVE COMMONS ATTRIBUTION 4.0 INTERNATIONAL (CC BY 4.0) LICENSE. 

**ID 49.** L. Bizovska, Z. Svoboda, M. Janura, M.C. Bisi, N. Vuillerme, Local dynamic stability during gait for predicting falls in elderly people: A one-year prospective study, PLoS ONE 13 (2018) e0197091. https://doi.org/10.1371/journal.pone.0197091.

Category: Methodological illustration

**Fig**

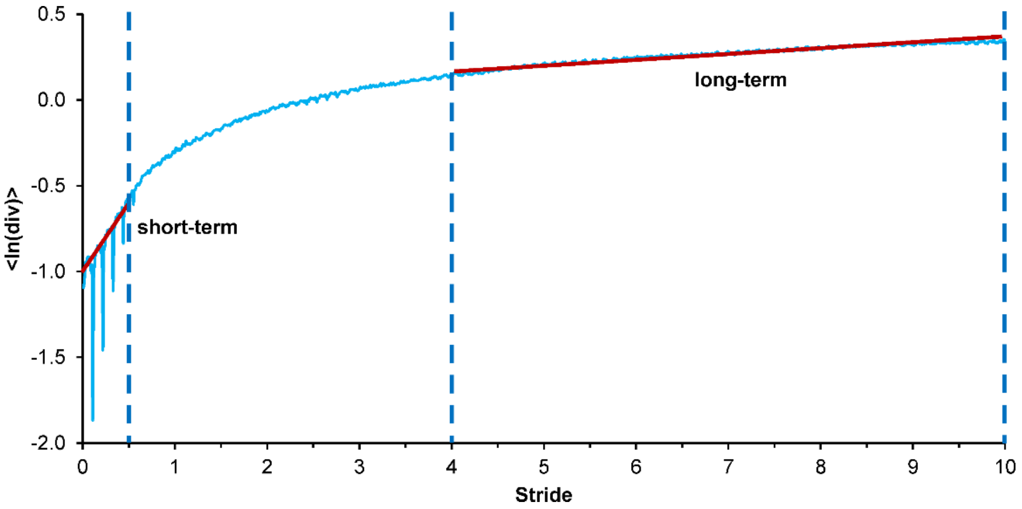


**1. Representation of short- and long-term LE computation.**

LE are computed as slopes of mean log divergence curve between 0 and 0.5 stride (short-term) and 4 and 10 strides (long-term).

COPYRIGHT 2025 THE AUTHORS. THIS FILE IS LICENSED UNDER A CREATIVE COMMONS ATTRIBUTION 4.0 INTERNATIONAL (CC BY 4.0) LICENSE. 

**ID 51.** P. Terrier, Complexity of human walking: the attractor complexity index is sensitive to gait synchronization with visual and auditory cues, PeerJ 7 (2019) e7417. https://doi.org/10.7717/peerj.7417.

Category: Comparative results

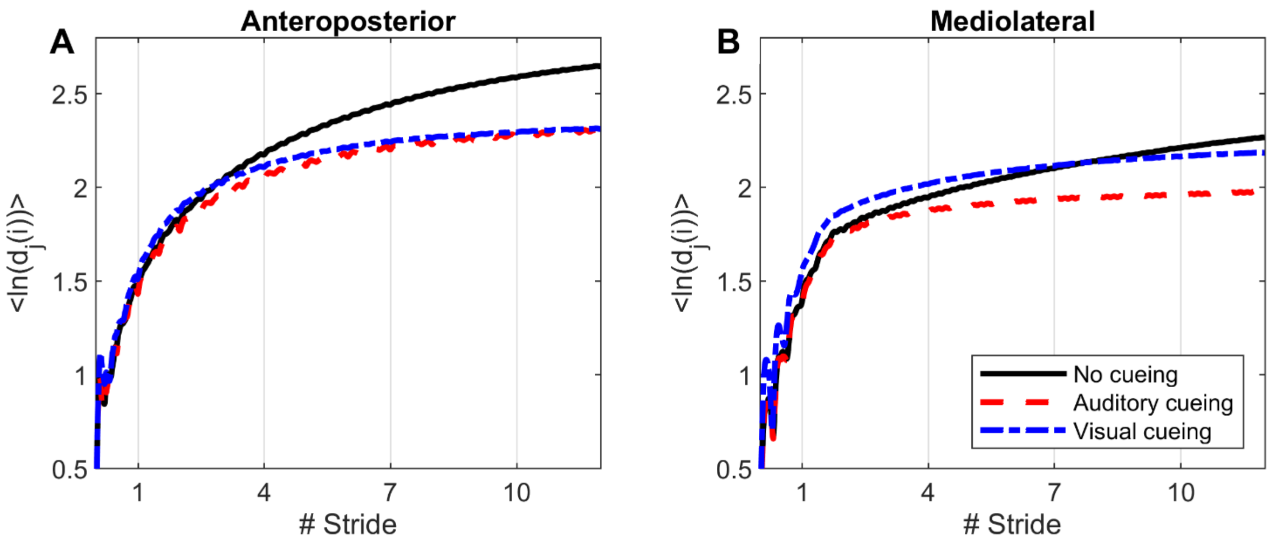


**Figure 1: Divergence curves.**

Using time-delay embedding, 5-dimensional attractors were reconstructed from the anteroposterior and mediolateral coordinates of a center-of-pressure trajectory. The logarithmic divergence from neighbor trajectories (*y*-axis) was averaged across trajectories and participants ($N$ = 36), and drawn against normalized time (strides, *x*-axis). Three curves are shown, one for each experimental condition.

COPYRIGHT 2025 THE AUTHORS. THIS FILE IS LICENSED UNDER A CREATIVE COMMONS ATTRIBUTION 4.0 INTERNATIONAL (CC BY 4.0) LICENSE.

**ID 55.** J. Sarvestan, P. Aghaie Ataabadi, Z. Svoboda, F. Alaei, R.B. Graham, The effects of mobile phone use on motor variability patterns during gait, PLoS ONE 17 (2022) e0267476. https://doi.org/10.1371/journal.pone.0267476.

Category: Methodological illustration

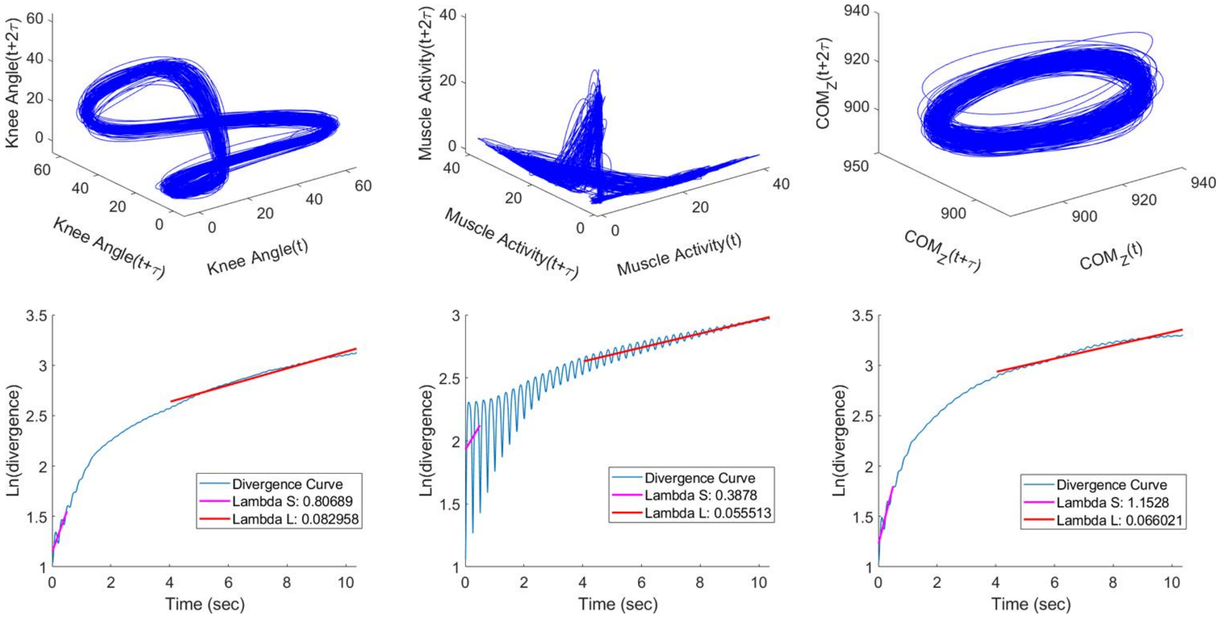


**Fig 1**

Schematic representation of the estimation of the maximum finite-time Lyapunov exponent for the knee sagittal plane (left), rectus femoris activities (centre) and COM trajectories (right). The 3D reconstruction of the phase-space and the expanded view of the reconstructed phase-space (up), and the average logarithmic rate of divergence for $\lambda_S$ and $\lambda_L$ (down).

COPYRIGHT 2025 THE AUTHORS. THIS FILE IS LICENSED UNDER A CREATIVE COMMONS ATTRIBUTION 4.0 INTERNATIONAL (CC BY 4.0) LICENSE.

**ID S2.** S. Piergiovanni, P. Terrier, Effects of metronome walking on long-term attractor divergence and correlation structure of gait: a validation study in older people, SCIENTIFIC REPORTS. 14 (2024). https://doi.org/10.1038/s41598-024-65662-5.

Category: Comparative results

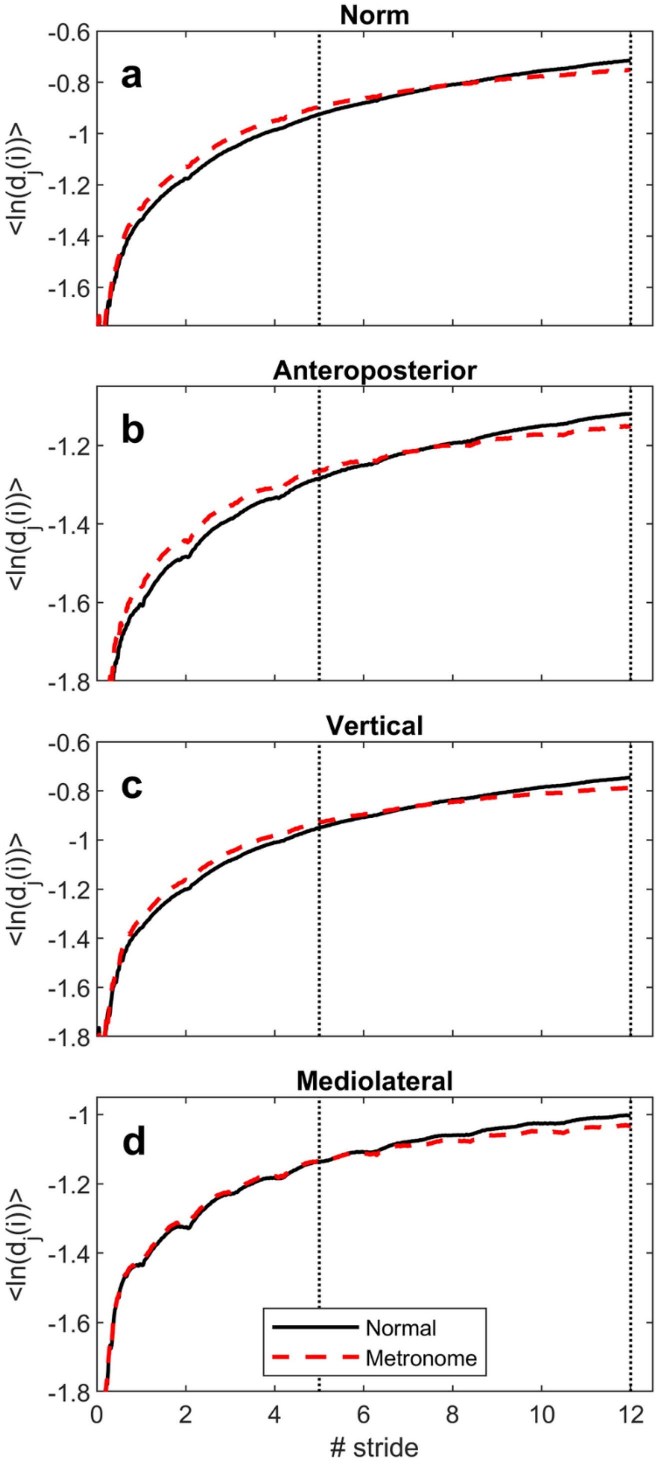


**Figure 1. Divergence curves.** The figure displays the average logarithmic divergence curves of gait dynamics obtained from the triaxial accelerometer attached to the lower back. The first subplot (**a**) shows the outcome of the vector norm (vector magnitude), while other subplots (**b**–**d**) represent each axis of the accelerometer. The curves are the averaged results of 58 participants for both normal and metronome walking. The figures display time on the x-axis, normalized by stride duration, and logarithmic divergence on the y-axis. Logarithmic divergence represents the natural log of the distance (*d*) between the *i*-th point downstream of the *j*-th pair of nearest neighbors in the state space (or attractor), averaged over all neighbors. The vertical dotted lines show the range over which the long-term divergence (or attractor complexity index) was computed.

COPYRIGHT 2025 THE AUTHORS. THIS FILE IS LICENSED UNDER A CREATIVE COMMONS ATTRIBUTION 4.0 INTERNATIONAL (CC BY 4.0) LICENSE. 
